# Controls of spatial grain size and environmental variables on observed beta diversity of molluscan assemblage at a regional scale

**DOI:** 10.1101/2022.11.02.514806

**Authors:** Madhura Bhattacherjee, Devapriya Chattopadhyay

**Affiliations:** Indian Institute of Science Education and Research Pune

**Keywords:** Scale of sampling, oceanographic variables, Arabian sea, live-dead fidelity

## Abstract

Beta diversity, which quantifies the compositional variation among communities, is one of the fundamental partitions of biodiversity and is associated with abiotic and biotic drivers. Unveiling these drivers is essential for understanding various ecological processes in past and recent faunal communities. Although the quantification of measures of beta diversity has improved over the years, the potential dependence of beta diversity on methodological choices are relatively understudied. Here, we investigate the effect of the variable scale of sampling on different measures of beta diversity at a regional scale. The west coast of India bordering the eastern margin of the Arabian sea, presents a coastal stretch of approximately 6100km from 8–21°N. We used marine bivalve distribution data, consisting of live occurrence data from literature reports and abundance data from death assemblages collected from localities representing latitude bins. We tested if the observed variation in beta diversity is explained by variable sampling scales due to differences in bin sizes and unequal coastline length. We developed a null model to generate a beta diversity pattern with an increase in spatial scale of sampling by increasing the spatial grain size along the 14 latitude bins progressively. Our null model demonstrates that for the both live and dead dataset, the total beta diversity measured by Bray-Curtis, Whittaker and Sorenson indices decreases with increasing sampling scale. The species replacement (turnover) evaluated by Simpson index decreases and the species loss (nestedness) measured by Sorenson index increases with increasing sampling scale. A comparison between the simulated and observed beta diversity distribution using K-S test demonstrated that the observed pattern of beta diversity is significantly different from the pattern generated from the null model in both live and death assemblages. This implies that sampling alone does not generate the spatial variation in beta diversity in this region. The results show that environmental parameters such as salinity, productivity, and cyclones play a significant role in shaping the regional beta diversity along the west coast. Our study provides an approach for evaluating the effect of variable sampling scale on comparing regional beta diversity. It also highlights the importance of spatial standardization while inferring about processes driving spatial diversity changes.

## Introduction

Biological diversity is spatially heterogeneous across the globe and understanding the causes of spatial variation in marine diversity is one of the major focus of ecological and paleoecological research (Kowalewski 1996; Olszewski and Patzkowsky 2001; Kidwell and Holland 2002; Huntley and Kowalewski 2007; Melo et al. 2009; Tittensor et al. 2010; Brown 2014; Tyler and Kowalewski 2017). The measures of spatial differences in diversity has three main partitions: alpha, beta, and gamma diversity (Whittaker 1960). Alpha and gamma diversity represents diversity at finest and largest scale of observation respectively (Patzkowsky and Holland 2012). Beta diversity, defined originally as the within-habitat diversity (Whittaker 1960), is used to quantify the spatial variation in community composition among localities (Harrison et al. 1992; Gray 2000; Anderson et al. 2011). Evaluating within-habitat differences in composition helps in understanding different aspects of ecosystem functioning (Legendre 2014), including drivers of community assembly and are considered essential for conservation based studies (Purvis and Hector 2000; Cleary 2003; Tuomisto et al. 2003; Baselga 2010).

Unlike the directly measurable alpha and gamma diversities, however, beta diversity is a derived quantity. It can be measured in numerous ways with no general consensus on which measure is suitable for particular ecological question making it a complex metric to interpret (Whittaker 1960; Anderson et al. 2006, 2011; Baselga 2010; Beck et al. 2013; Barwell et al. 2015). Beta diversity can be partitioned into two major components: turnover and nestedness (Harrison et al. 1992; Baselga 2007, 2010; Anderson et al. 2011). Turnover can be explained as the replacement of some species by others between assemblages along a gradient due to environment sorting and/or historical constraints such as dispersal barriers due to geographic isolation (Qian et al. 2005; Leprieur et al. 2011). In contrast, nestedness reflects a spatial pattern where assemblage of some sites with lower species richness are subsets of those sites with higher species richness as a result of processes such as selective extinction or colonization (Wright and Reeves 1992; Ulrich and Gotelli 2007). These components are not mutually exclusive and the resulting assemblages can be a mix of both components. Exploring these components across a gradient can reveal the role of different processes in shaping the patterns of assemblage composition along that gradient, which will in turn help in designing strategies for protecting the diversity of a landscape (Leprieur et al. 2011; Qian et al. 2020).

The patterns and processes influencing beta diversity has been an area of considerable research interest and the model organisms are dominated by terrestrial communities such as plants (Fournier and Loreau, 2001; Kraft et al., 2011; Qian et al., 2005; Qian and Ricklefs, 2007; Qian and Xiao, 2012; Wagner et al., 2000), insects (Fleishman et al. 2003; Gering et al. 2003; Summerville et al. 2003; Lindo and N. Winchester 2008), birds (Fleishman et al. 2003; Jankowski et al. 2009), mammals (Gabriel et al. 2006; Soininen et al. 2007; Melo et al. 2009; Svenning et al. 2011; Peixoto et al. 2017) and freshwater fauna (Stendera and Johnson 2005). In contrast, the marine communities are relatively poorly studied with the exception of reefal communities such as fishes and benthic invertebrates (Hewitt et al. 2005; Harborne et al. 2006; Josefson 2009; Belley and Snelgrove 2016; Roden et al. 2020; Souza et al. 2021). Large scale patterns in beta diversity is linked to latitudinal and altitudinal gradients (Soininen et al. 2007; Jankowski et al. 2009; Kraft et al. 2011). A combination of abiotic factors (such as temperature, habitat heterogeneity, biogeographic isolation events) and biotic factors (such as dispersal limitation, competitive exclusion) are attributed as important drivers of taxonomic and phylogenetic beta diversity in both terrestrial and marine realm (Becking et al. 2006; Qian and Ricklefs 2007; Arias-González et al. 2008; Leprieur et al. 2011; Baselga et al. 2012; Segre et al. 2014; Hattab et al. 2015; Klompmaker and Finnegan 2018; Fluck et al. 2020; Qian et al. 2020; Maxwell et al. 2022).

Identifying the drivers of beta diversity is highly dependent on spatial scale and resolution of the study (Mac Arthur and Wilson 1967; Hewitt et al. 2005; Tokeshi 2009). The factors that will determine variability in composition at a small spatial scale (site-scale or point-based studies) will be different from the determinant processes at larger scales. Typically, beta diversity increases rapidly at local scales as new sampling units are incorporated due high variation in stochastic species occupancy pattern among sites (Rosenzweig 1995; Barton et al. 2013). At regional scales, beta diversity increases more slowly as fewer newer species are encountered between sites as compared to local scales. At larger scales again beta diversity increases as new species are encountered between sites across bio-geographic regions with different geological and evolutionary histories.

Consequently similar patterns of beta diversity observed at different scales may not imply causative similarities (Whittaker et al. 2001; Hortal et al. 2010). Conceptually, beta diversity is expected to increase with increasing area with increasing spatial scale of individual units of observation (grain size) considering all individual units of observation (Barton et al. 2013). The choice of sample grain sizes even within a constant extent of study area, however, has a significant effect on the variability in species composition (Steinbauer et al. 2012). Barton et al (2013) proposed that a ‘sliding window’ perspective, in which both spatial grain size and extent varies would be an informative way for understanding compositional variation across scales. Uncertainties produced due to unequal sampling and variable geographic configuration further complicates the comparison of measured beta diversity (Womack et al. 2020). In spite of acknowledging the potential scale dependence, only a few studies attempted spatial scaling of beta diversity (Kraft et al. 2011; Barton et al. 2013; Womack et al. 2020). Moreover, the patterns of beta diversity and the sensitivity to sampling can differ among time-averaged death assemblages (DA) and live assemblages (LA) residing in various environments (Tyler and Kowalewski 2017). Understanding the effect of increasing spatial grain size implying increasing sample size per bin within a constant extent on observed beta diversity of both live and death assemblages will provide a unique insight into the spatial patterns of beta diversity.

The diverse ecosystem of tropical shallow marine environments is characterized by large number of co-existing species within habitats and high rates of species turnover between habitats (Gray 2000). Although these are important factors impacting beta-diversity (Segre et al., 2014; Klompmaker and Finnegan, 2018), only a handful of studies explored the regional patterns along tropical shallow marine environments. Using the marine bivalve distribution over a regional stretch of environmentally heterogeneous coastline of India, we evaluated the beta diversity and its dependence on the scale of study. Specifically, we tried to address the following questions:

i. If the variation in the beta diversity can be explained by unequal spatial grain size of sampling for LA and DA?
ii. What is the effect of the choice of beta diversity index on the observed pattern?
iii. If variations due to unequal sampling can be rejected, which environmental parameter contributes maximum to the observed beta diversity pattern?

## Materials and methods

### Locality and sampling

The study was conducted along the west coast of India. The west coast of India bordering the eastern Arabian Sea represent a latitudinal spread of 14° (8–23°N) spanning approximately 6100km from Kanyakumari in the south to Koteshwar in the north. The coast is characterized by high degree of environmental heterogeneity consisting of coral reefs, lagoons, seagrass habitats, sandy beaches. The northern part of Arabian sea has low siliciclastic input and high productivity associated with upwelling during winter cooling. The southern region has a well-developed reefal system with moderate variation in salinity (Parulekar and Wagh 1975; Slater 1984; Madhupratap et al. 1996; Levin et al. 2000; Sarkar et al. 2019). For collecting time-averaged death assemblage, a total of 25 sampling sites representing 14 latitudinal bins were selected. Each bin is represented by at least one sampling locality and with a gap of minimum five km between two consecutive localities within a bin. From each locality, all visible molluscan specimens were collected from a traverse of ~1 km along the sea shore. The procedure was repeated twice for each sampling site. The sampling has been done over a period of five and a half years from July 2010 to December 2015 in both post and pre-monsoon and a minimum of 200 individuals were collected from each latitudinal bin. Each latitudinal bin was represented by a minimum of 200 individuals. Taxonomic nomenclature was primarily based on the published work by Rao (2017) and the World Register of Marine Species (WoRMS Editorial Board, 2020). This is followed by detailed documentation and identification of the bivalve specimens from death assemblage (DA) (Chattopadhyay et al. 2021). For constructing live assemblage (LA) dataset, the occurrence data on marine bivalves was obtained from a marine biodiversity database reported from various published literature, maintained by the Bioinformatics Centre, National Institute of Oceanography, Goa, India (Sarkar et al., 2019). The database provided scientific name of the bivalves, along with taxonomic details, feeding habit, habitat, size and location. Location data are often supplemented by Google Earth for acquiring correct latitude and longitude.

### Oceanographic variables

Data on oceanographic variables (productivity, sea surface temperature, and salinity) were collected for each latitudinal bin from <u>Ocean Productivity database</u>. Diversity of shallow marine fauna is also known to be dependent on the area of the habitat (Smith and Benson 2013) and therefore we use shelf area and coastline length as a proxy for the habitat area. The coastal length and shelf width data are obtained from <u>GEBCO Compilation Group</u> (2020). Because high-energy storm events are known to affect the distribution of molluscan death assemblages (Bhattacherjee et al. 2021), we included cyclone frequency data from the global-tropical-extratropical cyclone climatic atlas from the United States Navy National Climate Data Center cyclone records. The details of the processing for converting the raw cyclone data into cyclone frequency are discussed at Bhattacherjee et al. (2021).

### Diversity estimates

Taxonomic beta diversity can be measured in a number of different ways. According to the concept of additive partitioning (Lande 1996), the gamma diversity (γ) in an area with multiple samples equals the sum of the average diversity within each of the samples (α) and among the samples (β), therefore γ = α + β, and β is given by γ – α (Crist et al. 2003). We report results using both classical additive metrics which are derived directly from the relationship between the alpha diversity and gamma diversity, such as Whittaker index (Lande 1996) and pairwise metric which are based on similarity between a pair of sites or an average of all pairs, and quantify turnover (Anderson et al. 2011). The pairwise metrices used are Sørensen (Sørensen,1948), Nestedness component of Sørensen, Simpson (Simpson 1943) and Bray-Curtis (Pairwise proportional dissimilarity) (Bray and Curtis 1957; Koleff et al. 2003; Anderson et al. 2006) indices. Sørensen dissimilarity measures the compositional dissimilarity component that is arising from species replacement and species loss (nestedness). The component of dissimilarity cause by species replacement is explained by Simpson dissimilarity (Simpson 1943). The nested component of Sørensen can be calculated by simply subtracting the Simpson dissimilarity from the Sørensen dissimilarity measure (Baselga 2010). The presence-absence version of the Bray-Curtis indices or pairwise proportional dissimilarity (PPD) been shown to be relatively insensitive to uneven sample sizes (Wolda 1981; Ferrier et al. 2007).

All calculations were performed on both the datasets (LA vs DA). The abundance data is transformed to presence-absence data prior to measurement of beta diversity. Classical beta diversity measures like Whittaker’s beta diversity are calculated in R using the “betadiver” function from the package Vegan and pairwise measures are calculated using the “beta.pair” function from the package betapart (Baselga and Orme 2012).

### Null model

The null hypothesis that we tested states that the variation in beta diversity along the coast is explained by unequal sampling due to differences in bin sizes and unequal coastline length. To test it, we developed a null model (Ulrich and Gotelli 2007; Astorga et al. 2014; Loiseau et al. 2017) with two versions: 1) Combined bin method and 2) Individual bin method (Fig 2). These versions allow prediction of the pattern of variation in beta diversity with increasing spatial grain and extent of observation.

**Figure 1:**
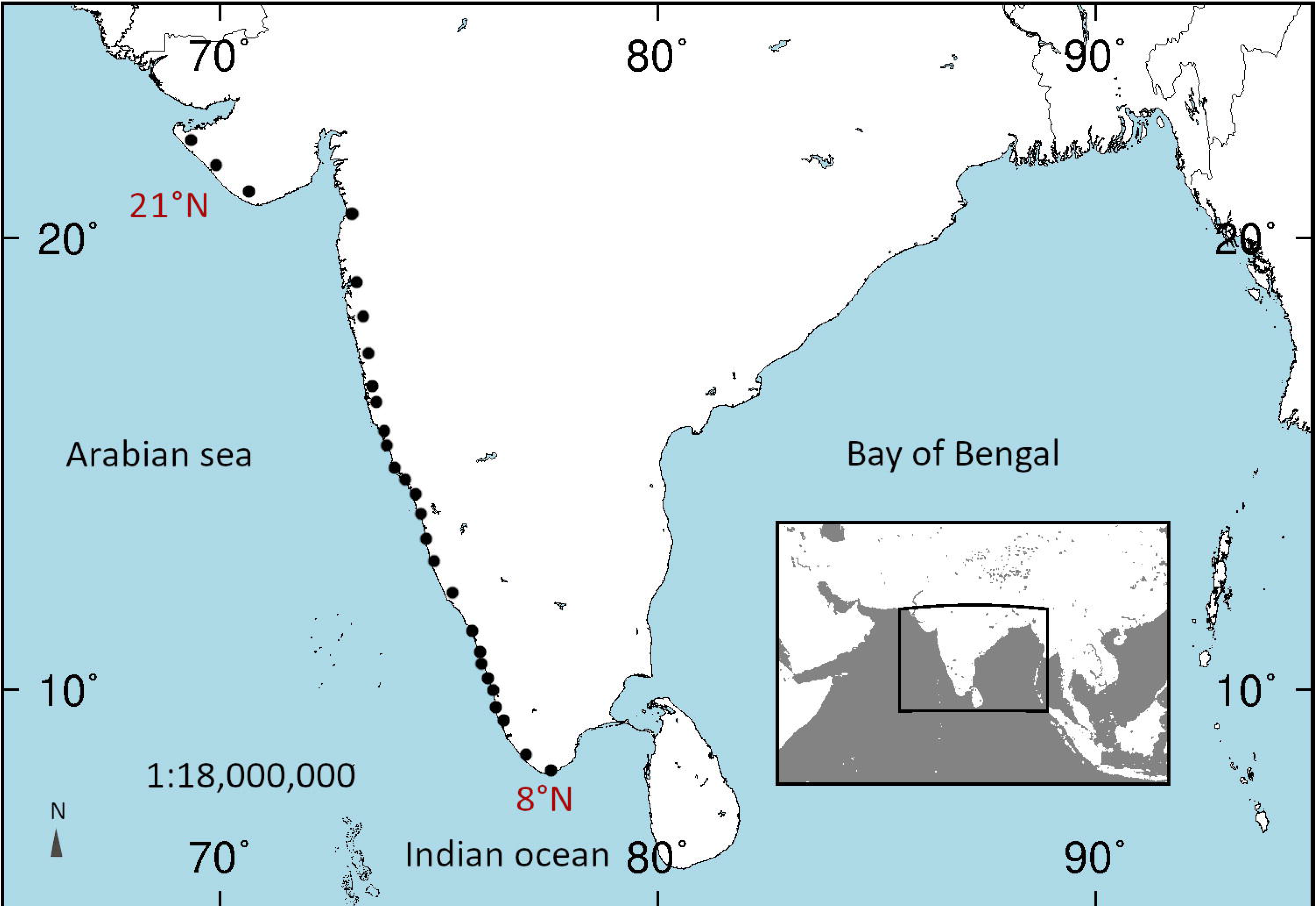
Map of India showing the sampling locations.

**Figure 2:**
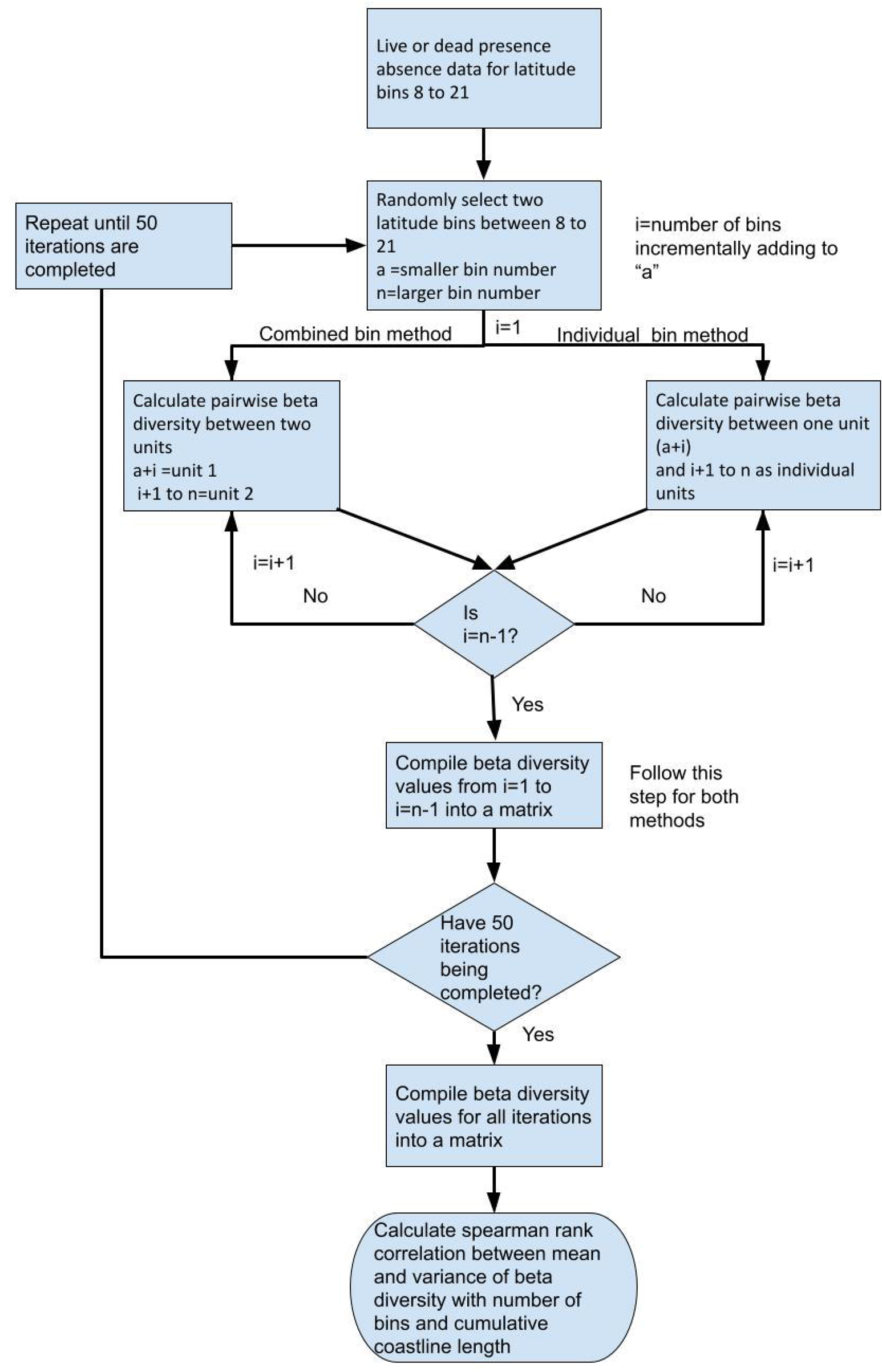
Flowchart describing the general framework for the null model

In both the variations, for each iteration, we randomly choose two latitude bins between 8 to 21, 8 being the southernmost bin and 21 being the northernmost bin. Each of these bins are of unequal sizes spanning variable coastline lengths. We consider each of these bins as grains or individual units of observation. Therefore, bin sizes or coastline length is considered as the measure of sampling scale in our study.

In the “Combined bin method” we incrementally increase the grain size from the smaller latitude bin towards the larger bin, by adding one bin in each step. The remaining latitudinal bins at that step are also clubbed together into a single unit. At each step, the beta diversity is calculated between that grain with the other unit containing the rest of the latitude bins combined together. The grain size from the smaller bin is increased at each step until it reaches the bin prior to the maximum latitude bin value of that iteration. This process is repeated for 50 iterations (Fig. 2). The beta diversity is calculated at each step of every iteration.

In the “Individual bin method”, we increase the grain size at each step from the smaller latitude bin by adding one bin in the same way as described in the “Combined bin method”. However, in contrast to “Combined bin method” where the rest of the latitude bins are clubbed together as one single unit, we consider the remaining latitudinal bins as individual units; the beta diversity at each step is calculated between that grain with the other individual latitude bins. The remaining steps within the first iteration are common to the “Combined bin method” and is followed in the same sequence as explained previously (Fig. 2). To evaluate the effect of choice of the beta diversity measure, both versions of the null model were performed using various beta diversity measures such as Whittaker (β_whit_), Bray Curtis (β_ppd_), Simpson (β_sim_), Sorenson (β_sor_) and nestedness component of Sorenson (β_sne_). Spearman rank-order coefficient is used to measure the correlation of beta diversity values (β_Null_) of each indices with varying bin sizes and coastline length. The model was used for both the LA and DA dataset and the results are compared.

To check the effect of unequal grain sizes on the observed beta diversity distribution we checked whether β_Obs_ (β_Obs_LA_ and β_Obs_DA_) can be generated from the distribution of null model values β_Null_ (β_Null_LA_ and β_Null_DA_). A resampling method similar to Bhattacherjee et al, 2021 was performed for simulating a distribution of β values (β_simulated_) by randomly sampling from the distribution of null model values (β_Null_). We resampled 14 values with replacement corresponding to 14 latitude bins from distribution of β_Null_ to generate a simulated distribution (β_simulated_). We calculated the K-S distance (D) and p-value between the distribution of simulated β values (β_simulated_) and observed β values (β_Obs_LA_ and β_Obs_DA_) using the ks.test () function in R. We repeated this step 10,000 times to get Bootstrap densities of K-S distances and p-values. This process is performed for all the β diversity indices. If β_Obs_ can be generated from β_Null_ then the K-S test will generate p values >0.005 implying the observed difference in beta diversity can be created by the scale-dependent sampling strategy. We can reject the null hypothesis if p<0.005 implying β_Obs_ cannot be generated from distribution of β_Null_. Therefore, that the variation in beta diversity cannot be explained by methodological issues such as sampling strategy alone and probably demonstrating the natural variation.

### Statistical analyses

To evaluate the relationship between β diversity and physical factors (such as latitude, coastline length and other environmental variables), we used Spearman rank-order correlation test. We used Bray-Curtis (PPD) dissimilarity for evaluating the correlation of β_Obs_ with environmental variables. We also used multiple generalized linear models (GLMs) to analyze the effect of environmental variables by taking all parameters simultaneously and evaluating their individual contributions to the total variation in diversity (Quinn and Keough 2002). To assess the change in species composition with environmental variables, a canonical correspondence analysis (CCA) and Redundancy Analysis (RDA) was conducted (ter Braak 1986). CCA uses a site-by-species matrix and a site by-environment matrix to extract orthogonal ordination axes that represent linear combinations of environmental variables. RDA is a canonical extension of principal component analysis (PCA), where ordination vectors are constrained by multiple regression to be linear combinations of the original explanatory variables (Legendre and Legendre 1998).

All statistical tests were performed in R version 4.2.0 (R Core Development Team, 2012).

## Results

The DA consists of a total of 13757 bivalve specimens collected from 25 localities over 14 latitude bins representing 167 species from 28 families. The LA consists of 177 species representing 37 families. Mean beta diversity values in LA varies from 0.156 for β_obs_sne_ to 0.864 for for β_obs_ppd_ (Table 1). Mean beta diversity values in DA varies from 0.151 for β_obs_sne_ to 0.851 for _β obs_ppd_ (Table 1).

**Table 1.** Mean of observed beta diversity values of different indices from LA and DA.

| **$\beta$ diversity measure** | **LA** | **DA** |
| --- | --- | --- |
| Bray-Curtis ($\beta_{\text{obs\_ppd}}$) | 0.851 | 0.864 |
| Whittaker ($\beta_{\text{obs\_whit}}$) | 0.753 | 0.629 |
| Simpson ($\beta_{\text{obs\_simp}}$) | 0.602 | 0.473 |
| Sorenson ($\beta_{\text{obs\_sor}}$) | 0.753 | 0.629 |
| Nestedness component of Sorenson ($\beta_{\text{obs\_sne}}$) | 0.151 | 0.156 |

### Predicted effect of sampling and choice of index on beta diversity

In the live assemblages (LA), the null model generated beta diversity values did not show any consistent pattern and the correlation was dependent on the unit of spatial grain (bins and coastline length) and the method used (Table 2). Bray-Curtis dissimilarity (β_ppd_) although not significantly correlated with coastline length was negatively correlated with number of bins in individual bin method. The negative correlation with coastline length is however significant in combined bin method (Figure 3A, S1A). The total dissimilarity component, is negatively correlated with coastline length. While the Simpson index (β_sim_) values shows a negative correlation with both coastline length and number of bins in individual bin method (Figure 3F, S1F), the nestedness component of Sorenson (β_sne_) is positively correlated with coastline length and number of bins in individual bin method. In combined bin method however, β_sne_ is negatively correlated with number of bins (Figure 3J, S1J).

**Table 2.** Results of Spearman rank correlation test between beta diversity and spatial scale of sampling (grain size) for LA.

| Metric | LA |  |  |  |  |  |  |  |  |  |  |  |  |  |  |  |
| --- | --- | --- | --- | --- | --- | --- | --- | --- | --- | --- | --- | --- | --- | --- | --- | --- |
|  | Number of bins |  |  |  |  |  |  |  | Coastline length |  |  |  |  |  |  |  |
|  | Combined bin method |  |  |  | Individual bin method |  |  |  | Combined bin method |  |  |  | Individual bin method |  |  |  |
|  | Mean |  | Variance |  | Mean |  | Variance |  | Mean |  | Variance |  | Mean |  | Variance |  |
|  | p | rho | p | rho | p | rho | p | rho | p | rho | p | rho | p | rho | p | rho |
| $\beta_{ppd}$ | 0.636 | -0.032 | **0.000** | **0.335** | **0.034** | **0.148** | 0.087 | -0.545 | **0.004** | **-0.191** | 0.341 | 0.382 | 0.296 | 0.074 | 0.141 | -0.473 |
| $\beta_{whit}$ | 0.496 | -0.046 | 0.503 | 0.227 | 0.388 | 0.056 | 0.354 | -0.293 | 0.106 | -0.109 | 0.356 | 0.309 | 0.975 | -0.002 | 0.451 | -0.254 |
| $\beta_{sim}$ | 0.519 | 0.047 | 0.232 | -0.418 | **0.000** | **-0.339** | 0.880 | 0.066 | 0.735 | -0.024 | 0.145 | -0.469 | **0.000** | **-0.277** | 0.945 | 0.030 |
| $\beta_{sor}$ | **0.012** | **-0.169** | 0.968 | -0.018 | 0.085 | 0.118 | 0.457 | -0.237 | 0.053 | -0.131 | 0.451 | -0.254 | 0.321 | 0.068 | 0.654 | -0.150 |
| $\beta_{sne}$ | **0.056** | **-0.139** | 0.299 | -0.345 | **0.000** | **0.471** | 0.225 | -0.400 | 0.252 | -0.084 | **0.021** | **-0.700** | **0.000** | **0.335** | 0.946 | -0.027 |

**Figure 3:**
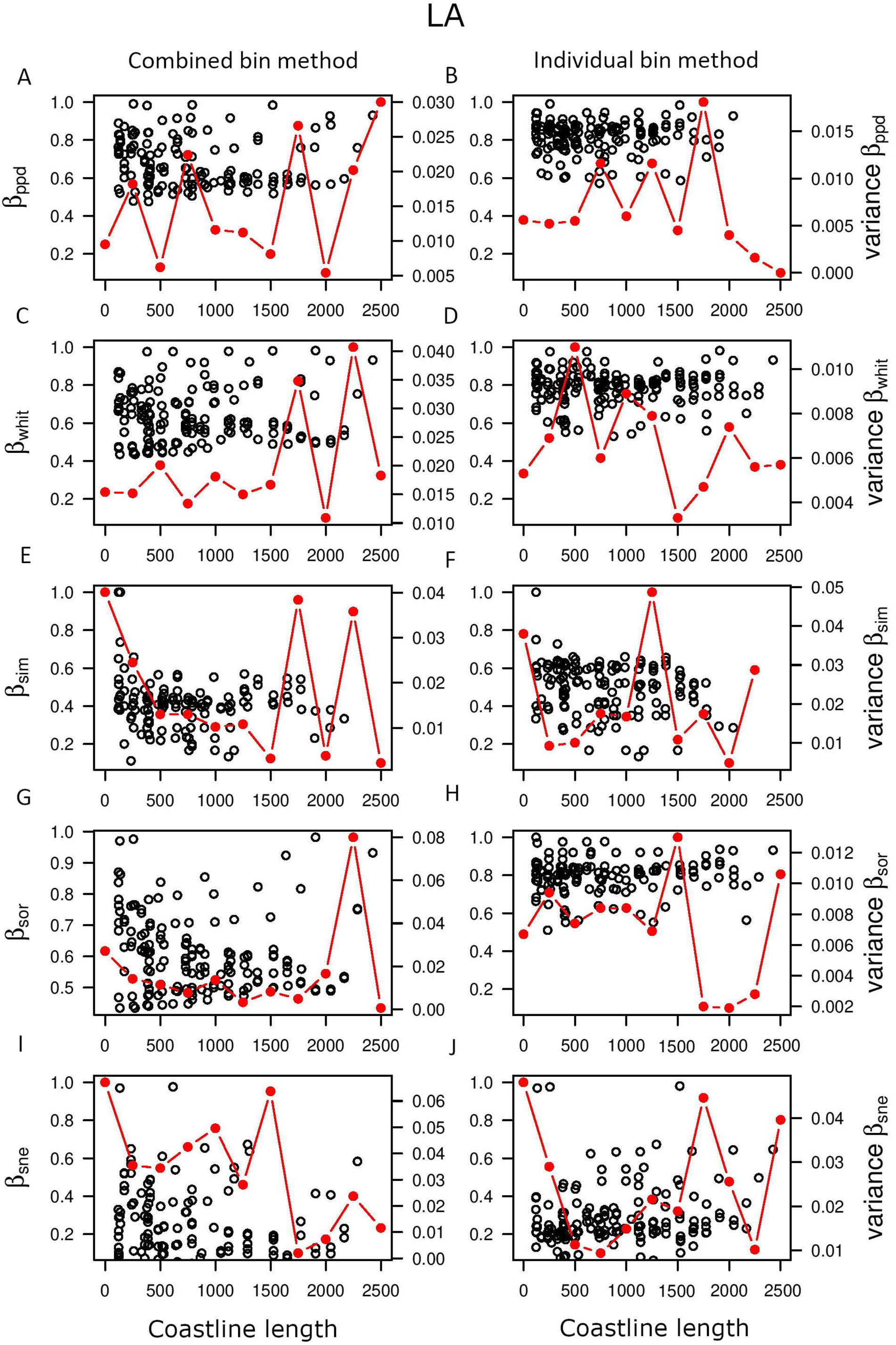
Null model predicted mean (black circles) and variance of beta diversity (red dash) with coastline length based on LA data. The left column represents “combined bin method” and the right column represents “individual bin method”. The indices of beta diversity used here include Bray-Curtis (β_ppd_) (A-B), Whittaker index (β_whit_) (C-D), Simpson index (β_sim_) (E-F), Sorenson index (β_sor_) (G-H), Nestedness component of Sorenson (β_sne_) (I-J).

In the DA’s, beta diversity of all indices from the null model, was significantly negatively correlated with number of bins and coastline length in combined bin method, except β_sim_ where correlation wasn’t significant with coastline length (Figure 4, S1; Table 3). Only Bray-Curtis (β_ppd_) was positively correlated with coastline length and number of bins for both methods (Figure 4A-B, S2A-B). Whittaker’s beta diversity (β_sim_) is negatively correlated with coastline length and number of bins combined bin method and only with bins in individual bin method (Figure 4C, S2C-D). The Simpson index (β_sim_) demonstrates a consistent negative correlation with coastline length in individual bin method and with number of bins in both methods (Figure 4E, S2E-F). Sorenson (β_sor_) shows a similar pattern to β_sim_ being negatively correlated with coastline length in combined bin method and with number of bins in both methods (Figure 4G, S2G, S2H). The variance in β_sim_ and β_sor_ is also negatively correlated with number of bins in both methods and coastline length in combined bin method (Figure 4E, 4G, S2E-H; Table 3). The nestedness component of Sorenson (β_sne_) on the other hand, shows a positive correlation with coastline length in individual bin method, although it is negatively correlated in combined bin method (Figure 4I-J, S2I). All the correlations mentioned before were significant, if not mentioned otherwise (Table 3).

**Table 3.** Results of Spearman rank correlation test between beta diversity and spatial scale of sampling (grain size) for DA.

| Metric | DA |  |  |  |  |  |  |  |  |  |  |  |  |  |  |  |
| --- | --- | --- | --- | --- | --- | --- | --- | --- | --- | --- | --- | --- | --- | --- | --- | --- |
|  | Number of bins |  |  |  |  |  |  |  | Coastline length |  |  |  |  |  |  |  |
|  | Combined bin method |  |  |  | Individual bin method |  |  |  | Combined bin method |  |  |  | Individual bin method |  |  |  |
|  | Mean |  | Variance |  | Mean |  | Variance |  | Mean |  | Variance |  | Mean |  | Variance |  |
|  | p | rho | p | rho | p | rho | p | rho | p | rho | p | rho | p | rho | p | rho |
| $\beta_{ppd}$ | **0.000** | **0.315** | 0.758 | -0.115 | **0.001** | **0.221** | 0.095 | -0.563 | **0.000** | **0.292** | 0.341 | -0.318 | **0.001** | **0.227** | 0.236 | -0.391 |
| $\beta_{whit}$ | **0.000** | **-0.436** | 0.967 | 0.018 | **0.002** | **-0.214** | 0.707 | -0.139 | **0.000** | **-0.353** | 0.145 | -0.472 | 0.752 | 0.220 | 0.802 | -0.091 |
| $\beta_{sim}$ | **0.045** | **-0.141** | **0.001** | **-0.872** | **0.000** | **-0.474** | 0.100 | -0.527 | 0.072 | -0.126 | **0.004** | **-0.809** | **0.000** | **-0.263** | 0.327 | 0.327 |
| $\beta_{sor}$ | **0.000** | **-0.428** | 0.427 | -0.284 | **0.000** | **-0.271** | **0.033** | **-0.654** | **0.000** | **-0.327** | **0.016** | **-0.718** | 0.658 | 0.029 | 0.192 | -0.427 |
| $\beta_{sne}$ | **0.000** | **-0.354** | 0.743 | 0.133 | 0.097 | 0.119 | 0.503 | 0.227 | **0.001** | **-0.222** | 0.349 | -0.333 | **0.022** | **0.163** | 0.313 | -0.336 |

**Figure 4:**
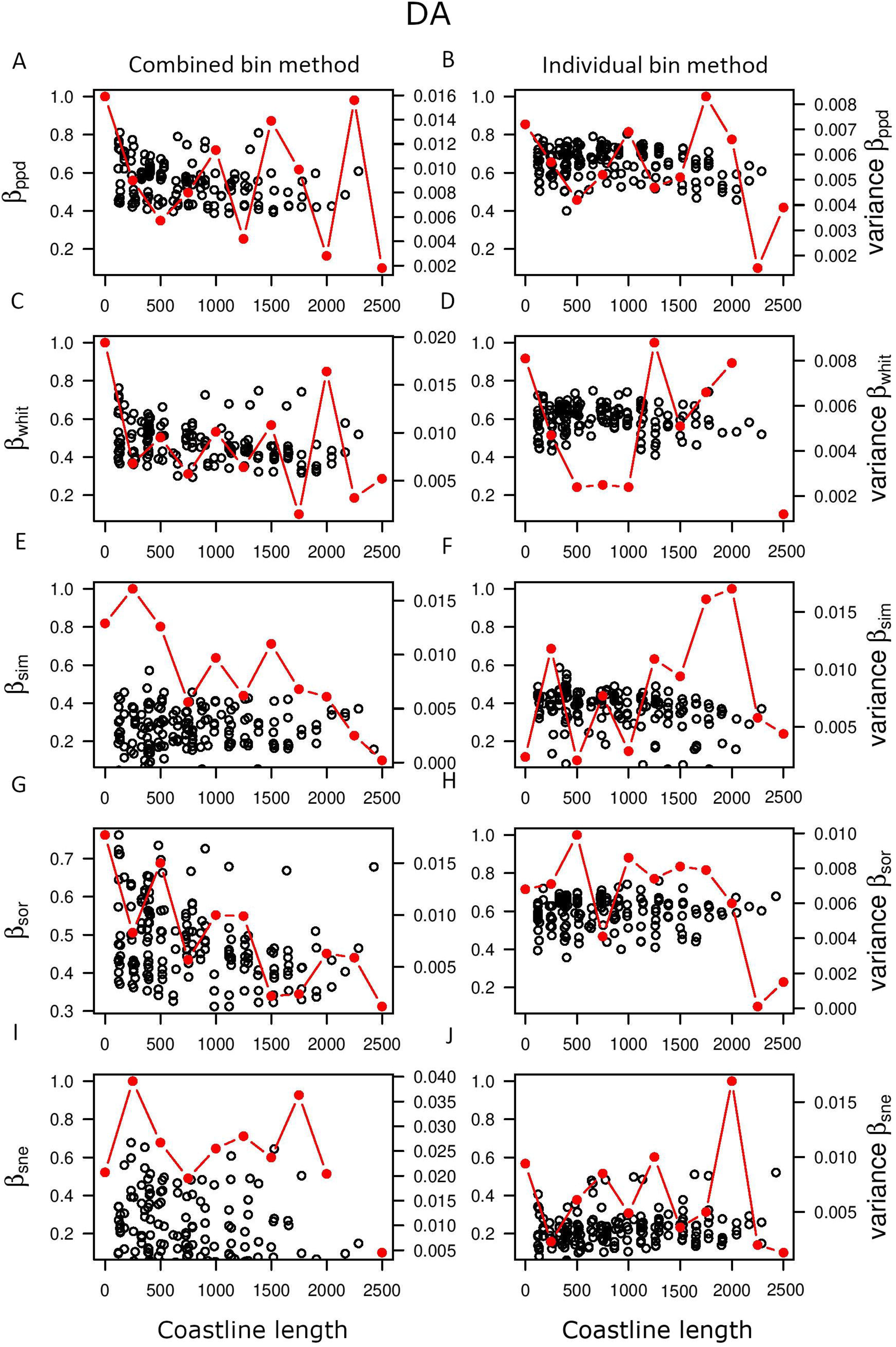
Null model predicted mean (black circles) and variance of beta diversity (red dash) with coastline length based on DA data. The left column represents “combined bin method” and the right column represents “individual bin method”. The indices of beta diversity used here include Bray-Curtis (β_ppd_) (A-B), Whittaker index (β_whit_) (C-D), Simpson index (β_sim_) (E-F), Sorenson index (β_sor_) (G-H), Nestedness component of Sorenson (β_sne_) (I-J).

### Effect of sampling scale and choice of index on observed beta diversity pattern

The observed variation pattern of beta diversity along the west coast also does not show any significant correlation with coastline length in both LA and DA (Figure 5). The distribution of β_Obs_ is significantly different from β_Null_ in K-S test for all beta diversity indices except nestedness component of Sorenson (β_sne_) in combined bin method (Figure 6). In individual bin method, the difference is significant only for β_sor_ distribution in live assemblages and for β_ppd_ in live assemblages (Figure 6 D, N). β_Null_ and β_Obs_ in nestedness component of Sorenson(β_sne_) is never significantly different in both methods (Figure 6Q-T). Most of the results for results for total dissimilarity indices β_sor_ and β_ppd_ are significant (Figure 6 A, C,D,M,N,O), which implies that they are sensitive proxies that can be used to evaluate methodological influence. Since β_ppd_ is showing a consistent pattern, it is a good index for determining the effect of sampling scale.

**Figure 5:**
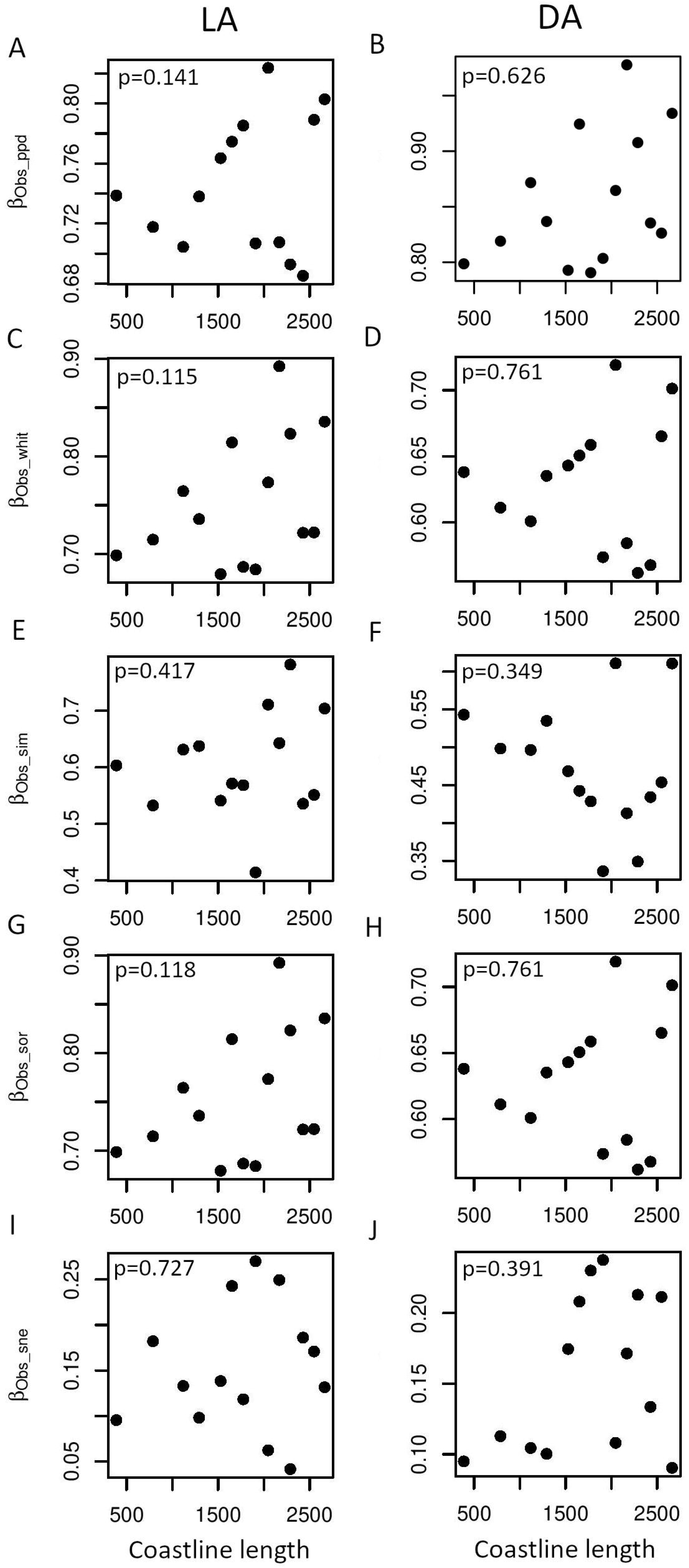
Relationship between observed mean beta diversity and coastline length. The left column represents LA and the right column represents DA. The indices of beta diversity used here include Bray-Curtis (β_ppd_) (A-B), Whittaker index (β_whit_) (C-D), Simpson index (β_sim_) (E-F), Sorenson index (β_sor_) (G-H), Nestedness component of Sorenson (β_sne_) (I-J).

**Figure 6:**
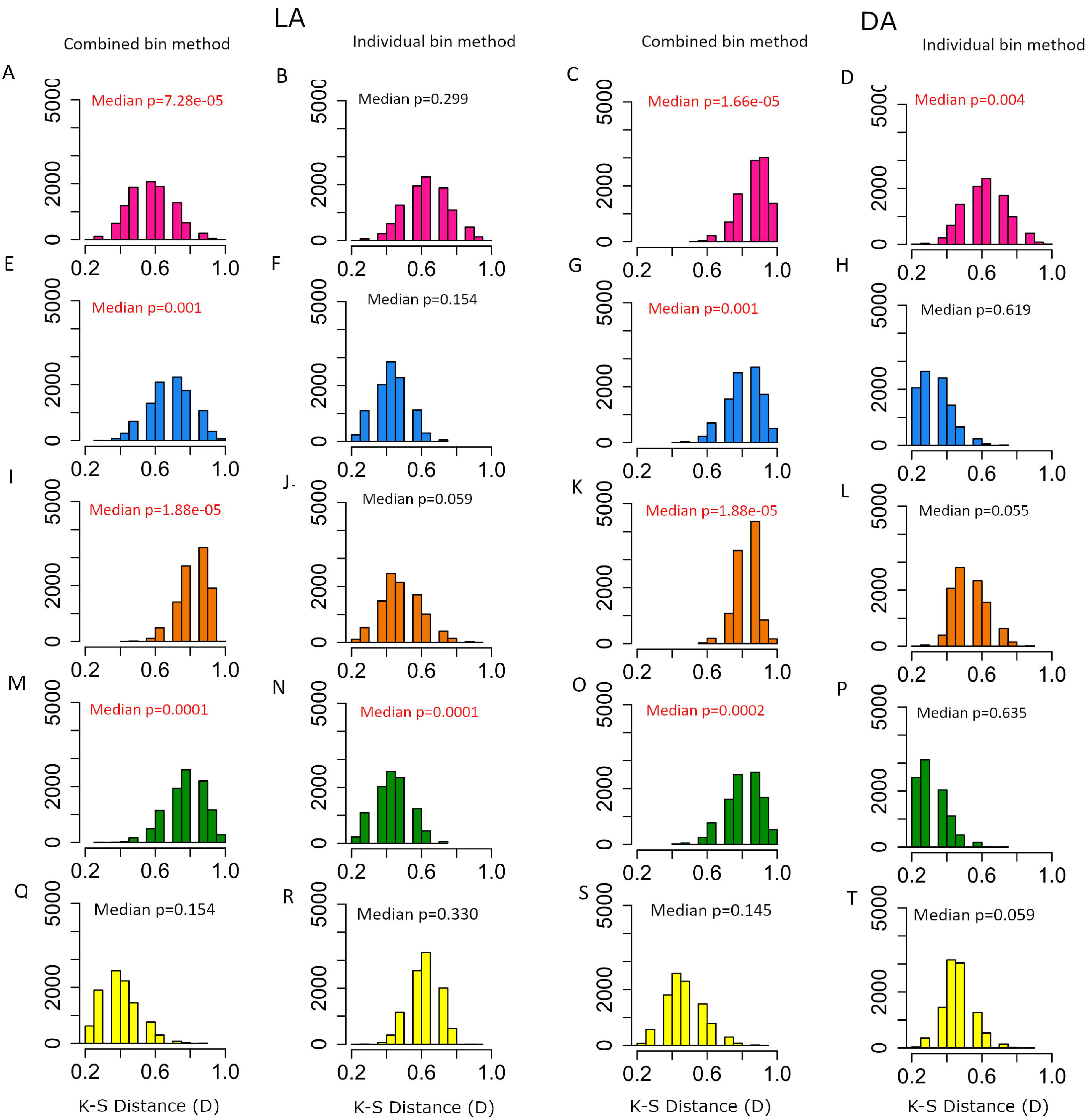
Histograms of D-values produced by K-S test between simulated (combined and individual method) and observed beta diversity distributions. The first two columns represent LA and the right two columns represent DA. The indices of beta diversity used here include Bray-Curtis (β_ppd_) (A-D), Whittaker index (β_whit_) (E-H), Simpson index (β_sim_) (I-L), Sorenson index (β_sor_) (M-P), Nestedness component of Sorenson (β_sne_) (Q-T). The significant p-values are marked in red.

### Overlapping and non-overlapping patterns in LA and DA

For LA and DA, beta diversity patterns were from the null model were mostly different especially when the combined bin method was used. While LA did not show a significant correlation with number of bins in β_ppd_ β_whit_ β_sim_, DAs were strongly negatively correlated with number of bins for all indices (Table 2-3). The patterns in LA were not as consistent as the patterns observed in the DA. β_ppd_ shows a significant positive correlation with bins in individual bin method in both the LA and DA (Figure S1B, S2B). The turnover component (β_sim_) shows a negative correlation with coastline length and number of bins in individual bin method in both the datasets (Figure 3F,4F, S1F, S2F; Table 2-3). The nestedness component (β_sne_) on the other hand positively correlated with coastline length in individual bin method and negatively correlated with number of bins in combined bin method in both LA and DA (Figure 3J, 4J, S1I, S2I). Overall, both LAs and DAs showed a decreasing pattern in the total dissimilarity components as well as turnover components with increase in sampling scale, except for the nestedness component which showed an increasing pattern. Β_Null_LA_ and β_Null_DA_ produced by both the LA and DA datasets were significantly different from the β_Obs_LA_ and β_Obs_DA_ of respective LA and DA datasets, in combined bin method. In individual bin method, however β_Null_ and β_Obs_ difference was not significant for most indices in both LA and DA except for β_sor_ in LA and β_ppd_ in DA which were significantly different. Except for two instances (Figure 6B-D, 6N-P), LA and DA behaved the same for all treatments (index, type of null model). This implies that the sensitivity to sampling scale is similar for both live assemblages and time-averaged death assemblages.

### Effect of environmental variables on beta diversity

Because of the robustness of β_ppd_ (Figure 6), we selected this index to evaluate the influence of the environmental variables on beta diversity. Only oxygen concentration shows a significant negative correlation with Bray-Curtis dissimilarity (β_ppd_) in LAs (Figure 7M). Salinity (range) shows a significant relationship with other environmental variables (Table 4). After excluding salinity (range) based on autocorrelation, none of the explanatory variables show significant effect on the beta diversity in single and multiple or single GLM for LA and DA (Table 5).

**Table 4.**
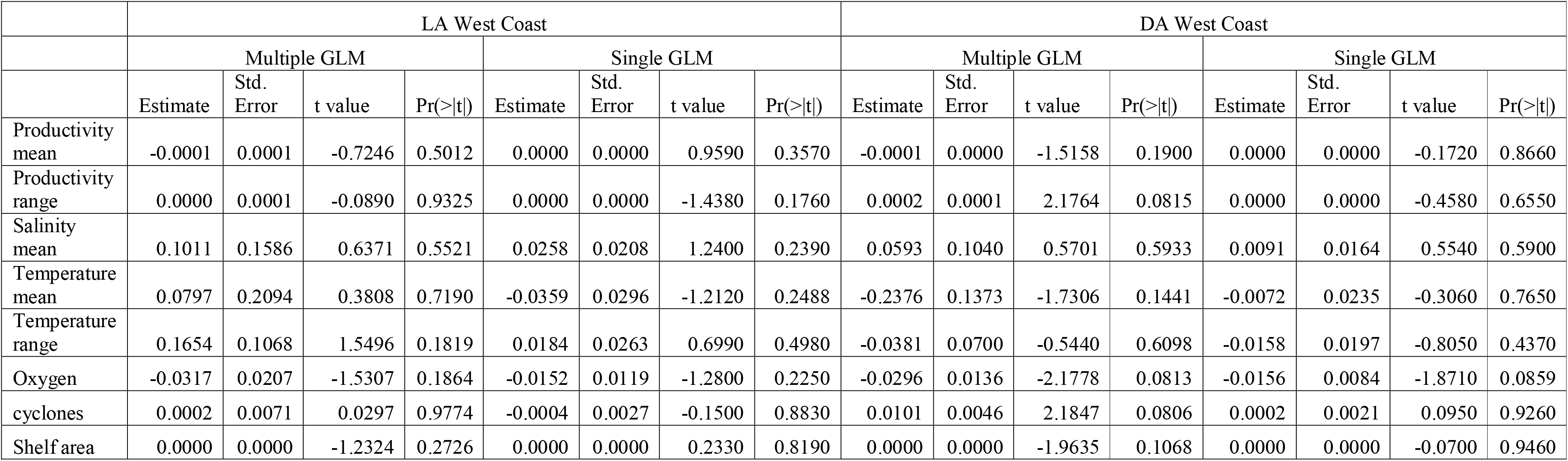
Results of multiple and single GLM analyses to assess contribution of environmental variables in determining observed Bray-Curtis dissmilarity (β_obs_ppd_). The significant results are in red.

|  | LA West Coast |  |  |  |  |  |  |  | DA West Coast |  |  |  |  |  |  |  |
| --- | --- | --- | --- | --- | --- | --- | --- | --- | --- | --- | --- | --- | --- | --- | --- | --- |
|  | Multiple GLM |  |  |  | Single GLM |  |  |  | Multiple GLM |  |  |  | Single GLM |  |  |  |
|  | Estimate | Std. Error | t value | Pr(>\|t\|) | Estimate | Std. Error | t value | Pr(>\|t\|) | Estimate | Std. Error | t value | Pr(>\|t\|) | Estimate | Std. Error | t value | Pr(>\|t\|) |
| Productivity mean | -0.0001 | 0.0001 | -0.7246 | 0.5012 | 0.0000 | 0.0000 | 0.9590 | 0.3570 | -0.0001 | 0.0000 | -1.5158 | 0.1900 | 0.0000 | 0.0000 | -0.1720 | 0.8660 |
| Productivity range | 0.0000 | 0.0001 | -0.0890 | 0.9325 | 0.0000 | 0.0000 | -1.4380 | 0.1760 | 0.0002 | 0.0001 | 2.1764 | 0.0815 | 0.0000 | 0.0000 | -0.4580 | 0.6550 |
| Salinity mean | 0.1011 | 0.1586 | 0.6371 | 0.5521 | 0.0258 | 0.0208 | 1.2400 | 0.2390 | 0.0593 | 0.1040 | 0.5701 | 0.5933 | 0.0091 | 0.0164 | 0.5540 | 0.5900 |
| Temperature mean | 0.0797 | 0.2094 | 0.3808 | 0.7190 | -0.0359 | 0.0296 | -1.2120 | 0.2488 | -0.2376 | 0.1373 | -1.7306 | 0.1441 | -0.0072 | 0.0235 | -0.3060 | 0.7650 |
| Temperature range | 0.1654 | 0.1068 | 1.5496 | 0.1819 | 0.0184 | 0.0263 | 0.6990 | 0.4980 | -0.0381 | 0.0700 | -0.5440 | 0.6098 | -0.0158 | 0.0197 | -0.8050 | 0.4370 |
| Oxygen | -0.0317 | 0.0207 | -1.5307 | 0.1864 | -0.0152 | 0.0119 | -1.2800 | 0.2250 | -0.0296 | 0.0136 | -2.1778 | 0.0813 | -0.0156 | 0.0084 | -1.8710 | 0.0859 |
| cyclones | 0.0002 | 0.0071 | 0.0297 | 0.9774 | -0.0004 | 0.0027 | -0.1500 | 0.8830 | 0.0101 | 0.0046 | 2.1847 | 0.0806 | 0.0002 | 0.0021 | 0.0950 | 0.9260 |
| Shelf area | 0.0000 | 0.0000 | -1.2324 | 0.2726 | 0.0000 | 0.0000 | 0.2330 | 0.8190 | 0.0000 | 0.0000 | -1.9635 | 0.1068 | 0.0000 | 0.0000 | -0.0700 | 0.9460 |

**Figure 7:**
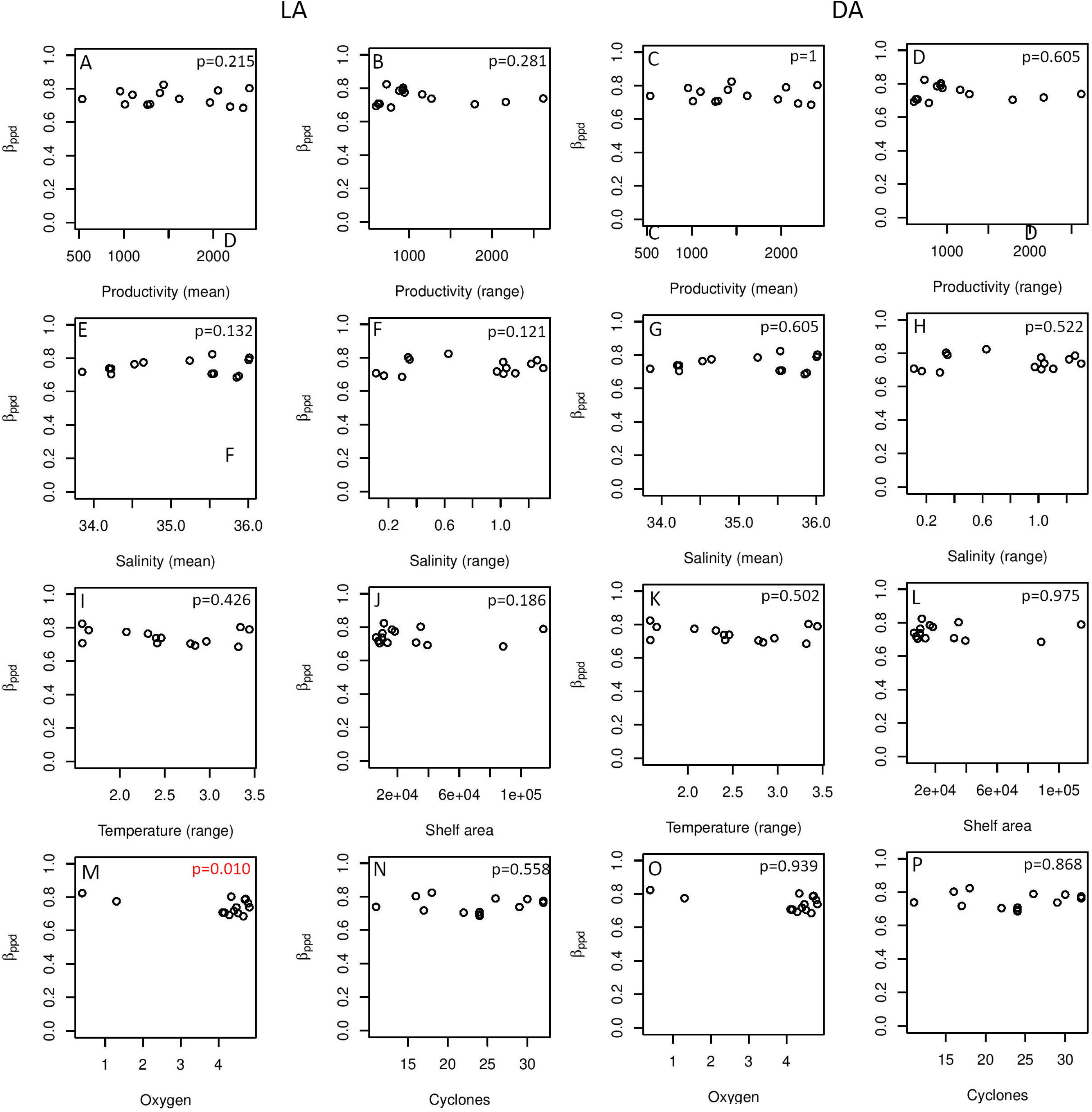
Relationship between β_ppd_ and different oceanographic parameters. The first two columns represent LA and the right two columns represent DA.

In Canonical correspondence analysis (CCA), 58% variation in species composition in LAs was explained by the environmental variables of salinity mean, productivity mean, productivity range, temperature mean, shelf area, oxygen concentration and cyclones (Figure 8A). Of the three ordination axes, axis 1 explained 12% of the total variation in the dataset and 42% of the variation explained by all three axes. The same combination of variables was able to explain 6.3% of the total variation in species composition in DAs (Figure 8B). In DAs, out of the three ordination axes, axis 1 explained 17% of the total variation in the dataset and 43.5 % of the variation explained by all three axes.

**Figure 8:**
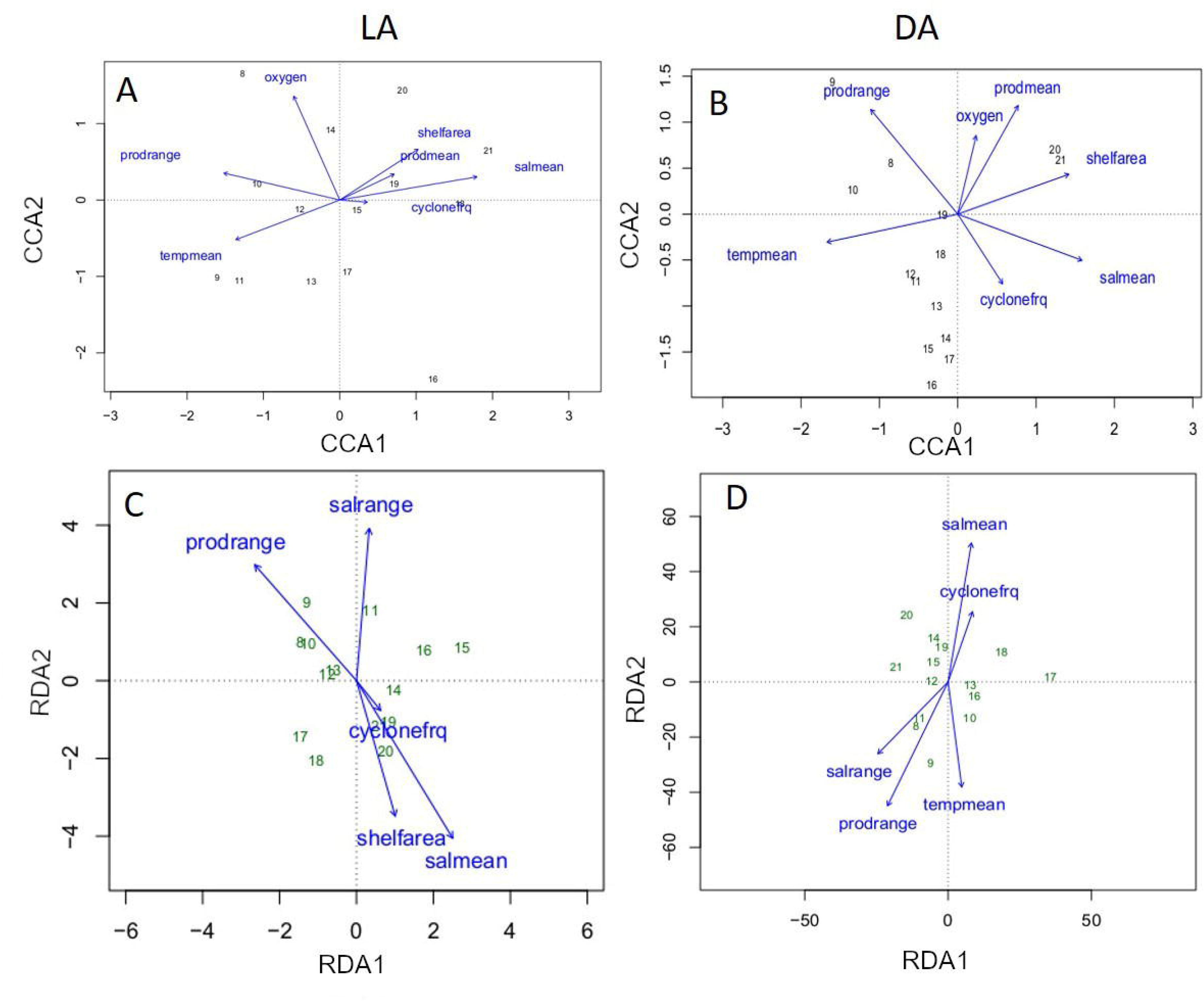
Biplots showing the relationship between β_ppd_ and environmental parameters using canonical correspondence analysis (CCA) (A-B) and redundancy analysis (RDA) (C-D). The left column represents LA and the right column represents DA.

About 50% of the constrained variation in species distribution in LAs is explained by a combination of productivity (range), salinity (mean), temperature (mean) and cyclones using RDA on presence-absence species data (Adjusted R^2^=23.7%) (Figure 8C). With forward selection, only salinity (mean and range) was found to be a significant predictor (p=0.03). The same set of variables along with shelf area were able to explain about 53% of the variation in species distribution in DAs (Adjusted R^2^=23.7%) (Figure 8D). Forward selection to choose a model with fewer variables, however stopped because of limited explanatory power of fewer environmental variables.

## Discussion

The compositional variation among communities captured by beta diversity is influenced by abiotic and biotic drivers. Unveiling these drivers of spatial heterogeneity of diversity requires us to rule out variations arising due to methodological strategies. The high marine diversity of tropical oceans, although studied in detail, their spatial structure is relatively poorly known. The west coast of India bordering the eastern Arabian Sea represents a tropical marine realm with a latitudinal spread of 14° (8–23°N) and is characterized by high degree of environmental heterogeneity. The alpha diversity of the coastal and shelf region of Arabian sea has been fairly well studied (Jayaraj et al. 2008; Joydas and Damodaran 2009, 2014). In contrast the beta diversity of macrobenthic species from this region has been largely unexplored (Sarkar et al. 2019; Sivadas et al. 2020, 2021). Our study attempts to develop a methodological framework to assess how beta diversity is influenced by methodological strategies such as spatial scale and diversity index. Using the regional distribution of LA and DA from a tropical coast with high environmental heterogeneity, it also attempts to identify the oceanographic drivers shaping the distribution.

### Effect of sampling scale

Coastline length is an important predictor of biodiversity of recent marine ecosystems (Tittensor et al. 2010). A longer coastline offers higher availability of important habitat features that positively influences both abundance and richness of coastal species (Rosenzweig 1995). However, variable coastline lengths of each latitude bin might lead to uneven sampling from different spatial bins resulting in increased beta diversity. A similar positive correlation is observed in alpha diversity studies, where alpha diversity increases quickly with increasing scale at smaller spatial scales due to high variation in stochastic species occupancy patterns among sampling units and variation in species responses to habitat heterogeneity (Rosenzweig 1995; Whittaker et al. 2001). At intermediate or regional scales, diversity increase with scale is slower because of fewer addition of new species relative to the regional pool. This pattern is also applicable to beta diversity, wherein dissimilarity is higher at smallest and biggest spatial scales but lower at intermediate scale (when based on “sliding window” with varying grain and extent) (Barton et al. 2013).

The null model provides a ‘sliding window’ perspective wherein the spatial grain size is increasing incrementally within a constant spatial extent. According to the results of our null model, the consistent pattern observed in beta diversity across LAs and DAs was a decreasing trend or negative correlation with increasing sampling scale (Table 2,3). This decreasing pattern is contradictory to the general theory of increasing beta diversity with increasing spatial grain within a constant extent (Barton et al. 2013; Womack et al. 2020). Harborne et al. (2006) observed a positive correlation of beta diversity with environmental conditions within a specific spatial scale across a tropical seascape supporting the importance of multiple scale-studies over single-scale studies for generalizing ecological patterns (Levin 1992). The observed pattern in beta diversity from mollusc LA and DAs from west coast, however doesn’t show any significant correlation with increasing coastline length. The null model generated distribution of beta diversity in this study provides an opportunity to quantitatively evaluate the effect of scale on the regional beta diversity. Our study also demonstrated that slight changes in the null model design may result in differing conclusion of the scale sensitivity. Combined bin method appeared to be more robust in identifying beta diversity variations developed due to non-methodological processes. This affirms that data categorization decisions can influence the observed beta diversity patterns at regional scales.

### Effect of choice of index

Unlike the overall diversity measures (alpha and gamma diversities) beta diversity cannot be measured directly. Because it is a derived quantity, the choice of measure is often debated as there is no general consensus on the suitability of a measure for addressing particular ecological question (Whittaker 1960; Anderson et al. 2006, 2011; Baselga 2010; Beck et al. 2013; Barwell et al. 2015). Moreover, the very concept of beta diversity is scale dependent and hence, the individual measures may differ in their sensitivity of the scale dependence. Our study shows that different measures of beta diversity may have a varying degree of sensitivity to spatial scale of sampling. Multisite pairwise measures of beta diversity (β_ppd_, β_sor_, β_whit_) shows a general negative correlation with increasing sampling scale represented by number of bins/coastline length, contrary to the a priori expectation of increasing beta diversity with increasing scale (Barton et al. 2013). Partitioning beta diversity into nestedness and species replacement components facilitates a greater understanding of patterns in beta diversity. However, we find a difference in their scale sensitivity implying a potential problem in interpreting observed patterns in beta diversity. In our study, the turnover component (β_sim_) decreases with increasing sampling scale whereas the nestedness component (β_sne_) increases, although in some of the analyses the nestedness component also decreases with increasing scale from the null model. Species replacement component or turnover component is the dominant component of variation in compositional dissimilarity and it is supposed to increase with increasing spatial scale, while the nestedness component decreases with increasing scale (Baselga 2007; Womack et al. 2020). The patterns of these components are logical consequences of the effect of environmental or ecological conditions that are operating at different scales. However, these studies have been performed at global scale where the role of dispersal limitation of species is higher and geographical differences in environmental conditions will also increase, which will likely increase the beta diversity. Our study has been performed at an intermediate scale in tropics, with a latitudinal range of 14 where such large scale geographical and dispersal limitation are less likely to occur. While comparing the simulated and observed pattern in beta diversity, the nestedness component (β_ppd_) did not show significant difference between observed and simulated patterns in any of the results, indicating that this index is not a reliable index in this context, as it cannot tell apart the methodological influence from the biological influence. Whereas, in total dissimilarity indices like β_sor_ and _βppd,_ the simulated and observed distribution are significantly different in most results. This implies that they are sensitive proxies that can be used to evaluate methodological influence. Therefore, we used β_ppd_ in our subsequent analyses for determining the contribution of environment.

### Patterns observed in LA and DA

Death assemblages showed a consistent pattern of negative correlation of beta diversity with increasing sampling scale from the null with the exception of nestedness component which showed a positive correlation with sampling scale. The live assemblages were also negatively correlated with sampling scale except β_sne_, however the correlation was significant in only very few analyses and indices. The observed beta diversity pattern in both DA and LA was not significantly correlated with coastline length and both showed the same signal of being significantly different from the predicted beta diversity pattern generated from the null model. In comparison of observed and simulated beta diversity, LA and DA behaved the same for all treatments (index, type of null model) except for two instances. This implies that the sensitivity to sampling scale is similar for both datasets. This means that in contrary to the previous observation that time averaging generally reduces the beta diversity in an assemblage (Tomašových and Kidwell 2009), our study demonstrated that time-averaged death assemblages and fossils are no worse than the LA when it comes to beta diversity scaling. Therefore, death assemblages preserve the biological signal that is observed in the live assemblages as observed in other marine assemblages (Tyler and Kowalewski 2017).. Time averaging and post-mortem mixing did not change the spatial fidelity in beta diversity pattern at a regional scale study such as this one.

### Role of environmental factors

Environmental variables of the eastern Arabian sea have a significant influence on beta diversity. Environmental processes often explain beta diversity at regional scales and lower latitudes (Qian and Ricklefs 2007). Studies showing stronger effect of environmental as opposed to spatial variables on community similarity have been reported from tropical forests and marine macrofauna in European marine sediments (Condit et al. 2002; Duivenvoorden et al. 2002; Ellingsen 2002; Ellingsen and Gray 2002; Cleary et al. 2004). In our study, salinity played a major role in determining the variability of the species composition in both northern and southern part of west coast based on the results of CCA and RDA. Salinity is one of the main structuring factor for macrobenthic species turnover at regional scale, as observed in the estuarine species in the northern Baltic sea where beta diversity changed at the same rate as the change in salinity between regions (Bleich et al. 2011; Josefson and Göke 2013). There is a significant variation in salinity in the southern part of west coast because of the influence of rainfall summer monsoons along with mixing of Bay of Bengal waters during winter months. This salinity variation is likely to affects marine benthos in the west coast.

Productivity also plays an important role in shaping up the diversity profile along a coastal region (Sarkar et al. 2019). A study on bacterial community distribution showed that surface water productivity explained 5.1% of the global pelagic community variation and 4.1% in the benthic realm (Zinger et al. 2011). Benthic marine communities showed dissimilarity with increasing scale than pelagic communities, since physical mixing plays a major role in homogenization of species composition (Zinger et al. 2011). In tropical marine reservoirs, diatom beta diversity was found to be negatively correlated to productivity (Zorzal-Almeida et al. 2017). Our study however doesn’t show any significant correlation of beta diversity with productivity. The productivity range plays a significant in controlling the variability of composition in both LA and DA as observed by the proximity of southern latitudinal bins to productivity range in RDA. This is because the west coast experiences an increase productivity due to upwelling processes with the onset of summer monsoon (June-September) (Madhupratap et al. 1996). During winter months, there is a rise in productivity in the surface layer mainly in the northeastern Arabian sea whereas the southern part has low productivity (Kumar and Prasad 1996; Madhupratap et al. 1996). The difference between summer and winter productivity is therefore higher in the southern Arabian sea, resulting in higher productivity range in the south.

Shelf area had a significant effect on the DAs but not LAs which is likely attributable to the fact that LAs have habitat specific patchy occurrences whereas DAs are more prone to post-mortem mixing. A greater shelf area indicates gentler slopes which causes lower rates of mixing whereas lower shelf area means a steeper slope causing higher rates of post-mortem transportation (Kidwell and Bosence 1991; Donovan 2002).

Our RDA plot (Figure 8C, 8D) shows a higher effect of cyclones on the species composition of the northern part of the west coast in both LA and DA as illustrated by the proximity of northern latitudinal bins to frequency of cyclones. Beta diversity across assemblages increased after storms events at a tidal flat in Brazil, owing to species loss after storms (Corte et al. 2017). There has been an increase in the intensity of pre-monsoon tropical cyclones over the Arabian Sea during recent years owing to an increase in the heat content in the ocean (Rajeevan et al. 2013). The cyclone tracks arriving from the western Arabian sea move in a northwesterly direction from 14°N to 17°N and gradually weakened towards north (Subrahmanyam et al. 2002). These cyclones thereby follow a northwesterly track thereby impacting the northern part of west coast more significantly.

The results of this study suggest that substantial variation in beta diversity can arise from methodological artefacts like uneven sampling and spatial resolution. Unless such variation is identified and accounted for, the true spatial pattern of biodiversity would be obscured and it will not be possible to identify the environmental drivers influencing the ecological processes.

## Conclusion

In conclusion, the present study analyzed the effect of sampling scale on the beta diversity at a regional scale using live and dead bivalve assemblages along west coast of India. The beta diversity pattern generated from null model provides us a reference to assess the effect of sampling scale on regional beta diversity pattern and its sensitivity on the choice of beta diversity index. Our analyses show that the observed beta diversity distribution in west coast cannot be explained by the null model alone implying uneven sampling to be a minor factor in shaping the beta diversity pattern. Among the environmental variables, salinity and productivity are major variables explaining the beta diversity of this region. Consistent patterns were obtained for live and dead datasets indicating that at regional scale, spatial and compositional fidelity have not changed significantly despite time averaging and post-mortem transportation events affecting the death assemblages. However, the consistency in this study should not be generalized to imply that live and death assemblages can be always considered to be congruent at regional scales and evaluation of live-dead fidelity should not be overlooked even at regional scales. A possible caveat of our study is the lack of detailed information on seasonal variation of the live assemblages as we had to mostly rely on snapshots of community data from literature. Because of the large spatial coverage pf our data, it is less likely to be severely impacted by the caveats.

## Supporting information

Supplementary R script (S3)

## Supplementary materials

Figure S1: Null model predicted mean (black circles) and variance of beta diversity (red dash) with number of bins based on LA data. The left column represents “combined bin method” and the right column represents “individual bin method”. The indices of beta diversity used here include Bray-Curtis (β_ppd_)(A-B), Whittaker index (β_whit_) (C-D), Simpson index (β_sim_) (E-F), Sorenson index (β_sor_) (G-H), Nestedness component of Sorenson (β_sne_) (I-J).

Figure S2: Null model predicted mean (black circles) and variance of beta diversity (red dash) with number of bins based on DA data. The left column represents “combined bin method” and the right column represents “individual bin method”. The indices of beta diversity used here include Bray-Curtis (β_ppd_) (A-B), Whittaker index (β_whit_) (C-D), Simpson index (β_sim_) (E-F), Sorenson index (β_sor_) (G-H), Nestedness component of Sorenson (β_sne_) (I-J).

Table S1. Significance (p-values) of Spearman rank correlation test between environmental variables. The significant results are in bold.

S3: This is the R-script used for the present study. The required data is available with the authors and can be shared upon request.

## References

Anderson, M. J., K. E. Ellingsen, and B. H. McArdle. 2006: Multivariate dispersion as a measure of beta diversity. Ecology Letters 9:683–693.

Anderson, M. J., T. O. Crist, J. M. Chase, M. Vellend, B. D. Inouye, A. L. Freestone, N. J. Sanders, H. V. Cornell, L. S. Comita, K. F. Davies, S. P. Harrison, N. J. B. Kraft, J. C. Stegen, and N. G. Swenson. 2011: Navigating the multiple meanings of β diversity: a roadmap for the practicing ecologist. Ecology Letters 14:19–28.

Arias-González, J. E., P. Legendre, and F. A. Rodríguez-Zaragoza. 2008: Scaling up beta diversity on Caribbean coral reefs. Journal of Experimental Marine Biology and Ecology 366:28–36.

Astorga, A., R. Death, F. Death, R. Paavola, M. Chakraborty, and T. Muotka. 2014: Habitat heterogeneity drives the geographical distribution of beta diversity: the case of New Zealand stream invertebrates. Ecology and Evolution 4:2693–2702.

Barton, P. S., S. A. Cunningham, A. D. Manning, H. Gibb, D. B. Lindenmayer, and R. K. Didham. 2013: The spatial scaling of beta diversity. Global Ecology and Biogeography 22:639–647.

Barwell, L. J., N. J. B. Isaac, and W. E. Kunin. 2015: Measuring β-diversity with species abundance data. Journal of Animal Ecology 84:1112–1122.

Baselga, A. 2007: Disentangling Distance Decay of Similarity from Richness Gradients: Response to Soininen et al. 2007. Ecography 30:838–841.

Baselga, A.. 2010: Partitioning the turnover and nestedness components of beta diversity. Global Ecology and Biogeography 19:134–143.

Baselga, A., and C. D. L. Orme. 2012: betapart: an R package for the study of beta diversity. Methods in Ecology and Evolution 3:808–812.

Baselga, A., J. M. Lobo, J.-C. Svenning, P. Aragón, and M. B. Araújo. 2012: Dispersal ability modulates the strength of the latitudinal richness gradient in European beetles. Global Ecology and Biogeography 21:1106–1113.

Beck, J., J. D. Holloway, and W. Schwanghart. 2013: Undersampling and the measurement of beta diversity. Methods in Ecology and Evolution 4:370–382.

Becking, L. E., D. F. R. Cleary, N. J. de Voogd, W. Renema, M. de Beer, R. W. M. van Soest, and B. W. Hoeksema. 2006: Beta diversity of tropical marine benthic assemblages in the Spermonde Archipelago, Indonesia. Marine Ecology 27:76–88.

Belley, R., and P. V. R. Snelgrove. 2016: Relative Contributions of Biodiversity and Environment to Benthic Ecosystem Functioning. Frontiers in Marine Science 3.

Bhattacherjee, M., D. Chattopadhyay, B. Som, A. S. Sankar, and S. Mazumder. 2021: Molluscan Live-Dead fidelity of a storm-dominated shallow-marine setting and its implications. Palaios 36:77–93.

Bleich, S., M. Powilleit, T. Seifert, and G. Graf. 2011: β-diversity as a measure of species turnover along the salinity gradient in the Baltic Sea, and its -consistency with the Venice System. Marine Ecology Progress Series 436:101–118.

Ter Braak, C. J. F. 1986: Canonical Correspondence Analysis: A New Eigenvector Technique for Multivariate Direct Gradient Analysis. Ecology 67:1167–1179.

Bray, J. R., and J. T. Curtis. 1957: An Ordination of the Upland Forest Communities of Southern Wisconsin. Ecological Monographs 27:326–349.

Brown, J. H. 2014: Why are there so many species in the tropics? Journal of Biogeography 41:8–22.

Chattopadhyay, D., D. Sarkar, and M. Bhattacherjee. 2021: The Distribution Pattern of Marine Bivalve Death Assemblage From the Western Margin of Bay of Bengal and Its Oceanographic Determinants. Frontiers in Marine Science 8.

Cleary, D. F. R. 2003: An examination of scale of assessment, logging and ENSO-induced fires on butterfly diversity in Borneo. Oecologia 135:313–321.

Cleary, D. F. R., A. Ø. Mooers, K. A. O. Eichhorn, J. Van Tol, R. De Jong, and S. B. J. Menken. 2004: Diversity and community composition of butterflies and odonates in an ENSO-induced fire affected habitat mosaic: a case study from East Kalimantan, Indonesia. Oikos 105:426–448.

Condit, R., N. Pitman, E. G. Leigh, J. Chave, J. Terborgh, R. B. Foster, P. Núñez, S. Aguilar, R. Valencia, G. Villa, H. C. Muller-Landau, E. Losos, and S. P. Hubbell. 2002: Beta-Diversity in Tropical Forest Trees. Science 295:666–669.

Corte, G. N., T. A. Schlacher, H. H. Checon, C. A. M. Barboza, E. Siegle, R. A. Coleman, and A. C. Z. Amaral. 2017: Storm effects on intertidal invertebrates: increased beta diversity of few individuals and species. PeerJ 5:e3360.

Crist, T. O., J. A. Veech, J. C. Gering, and K. S. Summerville. 2003: Partitioning Species Diversity across Landscapes and Regions: A Hierarchical Analysis of α, β, and γ Diversity. The American Naturalist 162:734–743.

Donovan, S. K. 2002: Island shelves, downslope transport and shell assemblages. Lethaia 35:277–277.

Duivenvoorden, J. F., J.-C. Svenning, and S. J. Wright. 2002: Beta Diversity in Tropical Forests. Science 295:636–637.

Ellingsen, K. E. 2002: Soft-sediment benthic biodiversity on the continental shelf in relation to environmental variability. Marine Ecology Progress Series 232:15–27.

Ellingsen, Karie., and J. s. Gray. 2002: Spatial patterns of benthic diversity: is there a latitudinal gradient along the Norwegian continental shelf? Journal of Animal Ecology 71:373–389.

Ferrier, S., G. Manion, J. Elith, and K. Richardson. 2007: Using generalized dissimilarity modelling to analyse and predict patterns of beta diversity in regional biodiversity assessment. Diversity and Distributions 13:252–264.

Fleishman, E., C. J. Betrus, and R. B. Blair. 2003: Effects of spatial scale and taxonomic group on partitioning of butterfly and bird diversity in the Great Basin, USA. Landscape Ecology 18:675–685.

Fluck, I. E., N. Cáceres, C. D. Hendges, M. do N. Brum, and C. S. Dambros. 2020: Climate and geographic distance are more influential than rivers on the beta diversity of passerine birds in Amazonia. Ecography 43:860–868.

Fournier, E., and M. Loreau. 2001: Respective roles of recent hedges and forest patch remnants in the maintenance of ground-beetle (Coleoptera: Carabidae) diversity in an agricultural landscape. Landscape Ecology 16:17–32.

Gabriel, D., I. Roschewitz, T. Tscharntke, and C. Thies. 2006: Beta Diversity at Different Spatial Scales: Plant Communities in Organic and Conventional Agriculture. Ecological Applications 16:2011–2021.

Gering, J. C., T. O. Crist, and J. A. Veech. 2003: Additive Partitioning of Species Diversity across Multiple Spatial Scales: Implications for Regional Conservation of Biodiversity. Conservation Biology 17:488–499.

Gray, J. S. 2000: The measurement of marine species diversity, with an application to the benthic fauna of the Norwegian continental shelf. Journal of Experimental Marine Biology and Ecology 250:23–49.

Harborne, A. R., P. J. Mumby, K. ZŻychaluk, J. D. Hedley, and P. G. Blackwell. 2006: Modeling the Beta Diversity of Coral Reefs. Ecology 87:2871–2881.

Harrison, S., S. J. Ross, and J. H. Lawton. 1992: Beta Diversity on Geographic Gradients in Britain. Journal of Animal Ecology 61:151–158.

Hattab, T., C. Albouy, F. Ben Rais Lasram, F. Le Loc’h, F. Guilhaumon, and F. Leprieur. 2015: A biogeographical regionalization of coastal Mediterranean fishes. Journal of Biogeography 42:1336–1348.

Hewitt, J. E., S. F. Thrush, J. Halliday, and C. Duffy. 2005: The Importance of Small-Scale Habitat Structure for Maintaining Beta Diversity. Ecology 86:1619–1626.

Hortal, J., N. Roura-Pascual, N. J. Sanders, and C. Rahbek. 2010: Understanding (insect) species distributions across spatial scales. Ecography 33:51–53.

Huntley, J. W., and M. Kowalewski. 2007: Strong coupling of predation intensity and diversity in the Phanerozoic fossil record. Proceedings of the National Academy of Sciences 104:15006–15010.

Jankowski, J. E., A. L. Ciecka, N. Y. Meyer, and K. N. Rabenold. 2009: Beta diversity along environmental gradients: implications of habitat specialization in tropical montane landscapes. Journal of Animal Ecology 78:315–327.

Jayaraj, J., A. Wan, M. Rahman, and Z. K. Punja. 2008: Seaweed extract reduces foliar fungal diseases on carrot. Crop Protection 27:1360–1366.

Josefson, A. B. 2009: Additive partitioning of estuarine benthic macroinvertebrate diversity across multiple spatial scales. Marine Ecology Progress Series 396:283–292.

Josefson, A. B., and C. Göke. 2013: Disentangling the effects of dispersal and salinity on beta diversity in estuarine benthic invertebrate assemblages. Journal of Biogeography 40:1000–1009.

Joydas, T. V., and R. Damodaran. 2009: Infaunal macrobenthos along the shelf waters of the west coast of India, Arabian Sea. IJMS Vol.38(2) [June 2009].

Joydas, T. V., and R. Damodaran. 2014: Infaunal macrobenthos of the oxygen minimum zone on the Indian western continental shelf. Marine Ecology 35:22–35.

Kidwell, S. M., and D. Bosence. 1991: Taphonomy and time averaging of marine shelly faunas. Pp.115–209 *in* Taphonomy: Releasing the Data Locked in the Fossil Record. Plenum, New York.

Kidwell, S. M., and S. M. Holland. 2002: The Quality of the Fossil Record: Implications for Evolutionary Analyses. Annual Review of Ecology and Systematics 33:561–588.

Klompmaker, A. A., and S. Finnegan. 2018: Extreme rarity of competitive exclusion in modern and fossil marine benthic ecosystems. Geology 46:723–726.

Koleff, P., K. J. Gaston, and J. J. Lennon. 2003: Measuring beta diversity for presence– absence data. Journal of Animal Ecology 72:367–382.

Kowalewski, M. 1996: Time-Averaging, Overcompleteness, and the Geological Record. The Journal of Geology 104:317–326.

Kraft, N. J. B., L. S. Comita, J. M. Chase, N. J. Sanders, N. G. Swenson, T. O. Crist, J. C. Stegen, M. Vellend, B. Boyle, M. J. Anderson, H. V. Cornell, K. F. Davies, A. L. Freestone, B. D. Inouye, S. P. Harrison, and J. A. Myers. 2011: Disentangling the Drivers of β Diversity Along Latitudinal and Elevational Gradients. Science 333:1755–1758.

Kumar, S. P., and T. G. Prasad. 1996: Winter cooling in the northern Arabian Sea. Current Science 71:834–841.

Lande, R. 1996: Statistics and Partitioning of Species Diversity, and Similarity among Multiple Communities. Oikos 76:5–13.

Legendre, P. 2014: Interpreting the replacement and richness difference components of beta diversity. Global Ecology and Biogeography 23:1324–1334.

Legendre, P., and L. Legendre. 1998: Numerical Ecology: Volume 20. Elsevier Science & Technology.

Leprieur, F., P. A. Tedesco, B. Hugueny, O. Beauchard, H. H. Dürr, S. Brosse, and T. Oberdorff. 2011: Partitioning global patterns of freshwater fish beta diversity reveals contrasting signatures of past climate changes. Ecology Letters 14:325–334.

Levin, L. A., J. D. Gage, C. Martin, and P. A. Lamont. 2000: Macrobenthic community structure within and beneath the oxygen minimum zone, NW Arabian Sea. Deep Sea Research Part II: Topical Studies in Oceanography 47:189–226.

Levin, S. A. 1992: The Problem of Pattern and Scale in Ecology: The Robert H. MacArthur Award Lecture. Ecology 73:1943–1967.

Lindo, Z., and N. N. Winchester. 2008: Scale dependent diversity patterns in arboreal and terrestrial oribatid mite (Acari: Oribatida) communities. Ecography 31:53–60.

Loiseau, N., G. Legras, M. Kulbicki, B. Mérigot, M. Harmelin-Vivien, N. Mazouni, R. Galzin, and J. c. Gaertner. 2017: Multi-component β-diversity approach reveals conservation dilemma between species and functions of coral reef fishes. Journal of Biogeography 44:537–547.

Mac Arthur, R. H., and E. O. Wilson. 1967: The Theory of Island Biogeography. Princeton University Press, p.

Madhupratap, M., S. P. Kumar, P. M. A. Bhattathiri, M. D. Kumar, S. Raghukumar, K. K. C. Nair, and N. Ramaiah. 1996: Mechanism of the biological response to winter cooling in the northeastern Arabian Sea. Nature 384:549–552.

Maxwell, M. F., F. Leprieur, J. P. Quimbayo, S. R. Floeter, and M. G. Bender. 2022: Global patterns and drivers of beta diversity facets of reef fish faunas. Journal of Biogeography 49:954–967.

Melo, A. S., T. F. L. V. B. Rangel, and J. A. F. Diniz-Filho. 2009: Environmental drivers of beta-diversity patterns in New-World birds and mammals. Ecography 32:226–236.

Olszewski, T. D., and M. E. Patzkowsky. 2001: Measuring Recurrence of Marine Biotic Gradients: A Case Study from the Pennsylvanian-Permian Midcontinent. PALAIOS 16:444–460.

Parulekar, A. H., and A. B. Wagh. 1975: Quantitative studies on benthic macrofauna of north-eastern Arabian Sea shelf. IJMS Vol.04(2) [December 1975].

Patzkowsky, M. E., and S. M. Holland. 2012: 6 The Ecology of Fossil Taxa through Time. Pp.113–130 *in* 6 The Ecology of Fossil Taxa through Time. University of Chicago Press.

Peixoto, F. P., F. Villalobos, A. S. Melo, J. a. F. Diniz-Filho, R. Loyola, T. F. Rangel, and M. V. Cianciaruso. 2017: Geographical patterns of phylogenetic beta-diversity components in terrestrial mammals. Global Ecology and Biogeography 26:573–583.

Purvis, A., and A. Hector. 2000: Getting the measure of biodiversity. Nature 405:212–219.

Qian, H., and R. E. Ricklefs. 2007: A latitudinal gradient in large-scale beta diversity for vascular plants in North America. Ecology Letters 10:737–744.

Qian, H., and M. Xiao. 2012: Global patterns of the beta diversity–energy relationship in terrestrial vertebrates. Acta Oecologica 39:67–71.

Qian, H., R. E. Ricklefs, and P. S. White. 2005: Beta diversity of angiosperms in temperate floras of eastern Asia and eastern North America. Ecology Letters 8:15–22.

Qian, H., Y. Jin, F. Leprieur, X. Wang, and T. Deng. 2020: Geographic patterns and environmental correlates of taxonomic and phylogenetic beta diversity for large-scale angiosperm assemblages in China. Ecography 43:1706–1716.

Quinn, G. P., and M. J. Keough. 2002: Experimental Design and Data Analysis for Biologists. Cambridge University Press, p.

Rajeevan, M., J. Srinivasan, K. Niranjan Kumar, C. Gnanaseelan, and M. M. Ali. 2013: On the epochal variation of intensity of tropical cyclones in the Arabian Sea. Atmospheric Science Letters 14:249–255.

Rao, N. S. 2017: Indian seashells: part-2 bivalvia. Zoological Survey of India Occasional Paper 375, p. 1–568

Roden, V. J., M. Zuschin, A. Nützel, I. M. Hausmann, and W. Kiessling. 2020: Drivers of beta diversity in modern and ancient reef-associated soft-bottom environments. PeerJ 8:e9139.

Rosenzweig, M. L. 1995: Species diversity in space and time. Cambridge University Press, p.

Sarkar, D., M. Bhattacherjee, and D. Chattopadhyay. 2019: Influence of regional environment in guiding the spatial distribution of marine bivalves along the Indian coast. Journal of the Marine Biological Association of the United Kingdom 99:163–177.

Segre, H., R. Ron, N. De Malach, Z. Henkin, M. Mandel, and R. Kadmon. 2014: Competitive exclusion, beta diversity, and deterministic vs. stochastic drivers of community assembly. Ecology Letters 17:1400–1408.

Simpson, G. G. 1943: Mammals and the nature of continents. American Journal of Science 241:1–31.

Sivadas, S. K., D. P. Singh, and R. Saraswat. 2020: Functional and taxonomic (α and β) diversity patterns of macrobenthic communities along a depth gradient (19–2639 m): A case study from the southern Indian continental margin. Deep Sea Research Part I: Oceanographic Research Papers 159:103250.

Sivadas, S. K., G. V. M. Gupta, S. Kumar, and B. S. Ingole. 2021: Trait-based and taxonomic macrofauna community patterns in the upwelling ecosystem of the southeastern Arabian sea. Marine Environmental Research 170:105431.

Slater, R. D. 1984: Controls of dissolved oxygen distribution and organic carbon deposition in the Arabian Sea. Geology and Oceanography of the Arabian Sea and Coastal Pakistan: 305–313.

Smith, A. B., and R. B. J. Benson. 2013: Marine diversity in the geological record and its relationship to surviving bedrock area, lithofacies diversity, and original marine shelf area. Geology 41:171–174.

Soininen, J., J. J. Lennon, and H. Hillebrand. 2007: A Multivariate Analysis of Beta Diversity Across Organisms and Environments. Ecology 88:2830–2838.

Souza, C. A. de, B. E. Beisner, L. F. M. Velho, P. de Carvalho, A. Pineda, and L. C. G. Vieira. 2021: Impoundment, environmental variables and temporal scale predict zooplankton beta diversity patterns in an Amazonian river basin. Science of The Total Environment 776:145948.

Steinbauer, M. J., K. Dolos, B. Reineking, and C. Beierkuhnlein. 2012: Current measures for distance decay in similarity of species composition are influenced by study extent and grain size. Global Ecology and Biogeography 21:1203–1212.

Stendera, S. E. S., and R. K. Johnson. 2005: Additive partitioning of aquatic invertebrate species diversity across multiple spatial scales. Freshwater Biology 50:1360–1375.

Subrahmanyam, B., K. H. Rao, N. Srinivasa Rao, V. S. N. Murty, and R. J. Sharp. 2002: Influence of a tropical cyclone on Chlorophyll-a Concentration in the Arabian Sea. Geophysical Research Letters 29:22-1-22–24.

Summerville, K. S., M. J. Boulware, J. A. Veech, and T. O. Crist. 2003: Spatial Variation in Species Diversity and Composition of Forest Lepidoptera in Eastern Deciduous Forests of North America. Conservation Biology 17:1045–1057.

Svenning, J.-C., C. Fløjgaard, and A. Baselga. 2011: Climate, history and neutrality as drivers of mammal beta diversity in Europe: insights from multiscale deconstruction. Journal of Animal Ecology 80:393–402.

Tittensor, D. P., C. Mora, W. Jetz, H. K. Lotze, D. Ricard, E. V. Berghe, and B. Worm. 2010: Global patterns and predictors of marine biodiversity across taxa. Nature 466:1098–1101.

Tokeshi, M. 2009: Species Coexistence: Ecological and Evolutionary Perspectives. John Wiley & Sons, p.

Tuomisto, H., K. Ruokolainen, and M. Yli-Halla. 2003: Dispersal, Environment, and Floristic Variation of Western Amazonian Forests. Science 299:241–244.

Tyler, C. L., and M. Kowalewski. 2017: Surrogate taxa and fossils as reliable proxies of spatial biodiversity patterns in marine benthic communities. Proceedings of the Royal Society B: Biological Sciences 284:20162839.

Ulrich, W., and N. J. Gotelli. 2007: Null Model Analysis of Species Nestedness Patterns. Ecology 88:1824–1831.

Wagner, H. H., O. Wildi, and K. C. Ewald. 2000: Additive partitioning of plant species diversity in an agricultural mosaic landscape. Landscape Ecology 15:219–227.

Whittaker, R. H. 1960: Vegetation of the Siskiyou Mountains, Oregon and California. Ecological Monographs 30:279–338.

Whittaker, R. J., K. J. Willis, and R. Field. 2001: Scale and species richness: towards a general, hierarchical theory of species diversity. Journal of Biogeography 28:453–470.

Wolda, H. 1981: Similarity indices, sample size and diversity. Oecologia 50:296–302.

Womack, T., M. Hannah, and J. Crampton. 2020: Spatial scaling of beta diversity in the shallow-marine fossil record. Paleobiology.

Wright, D. H., and J. H. Reeves. 1992: On the meaning and measurement of nestedness of species assemblages. Oecologia 92:416–428.

Zinger, L., L. A. Amaral-Zettler, J. A. Fuhrman, M. C. Horner-Devine, S. M. Huse, D. B. M. Welch, J. B. H. Martiny, M. Sogin, A. Boetius, and A. Ramette. 2011: Global Patterns of Bacterial Beta-Diversity in Seafloor and Seawater Ecosystems. PLOS ONE 6:e24570.

Zorzal-Almeida, S., L. M. Bini, and D. C. Bicudo. 2017: Beta diversity of diatoms is driven by environmental heterogeneity, spatial extent and productivity. Hydrobiologia 800:7–16.

