## Supplementary figures and images for "Controls of spatial grain size and environmental variables on observed beta diversity of molluscan assemblage at a regional scale"

### Supplementary Figure (Figure S1)

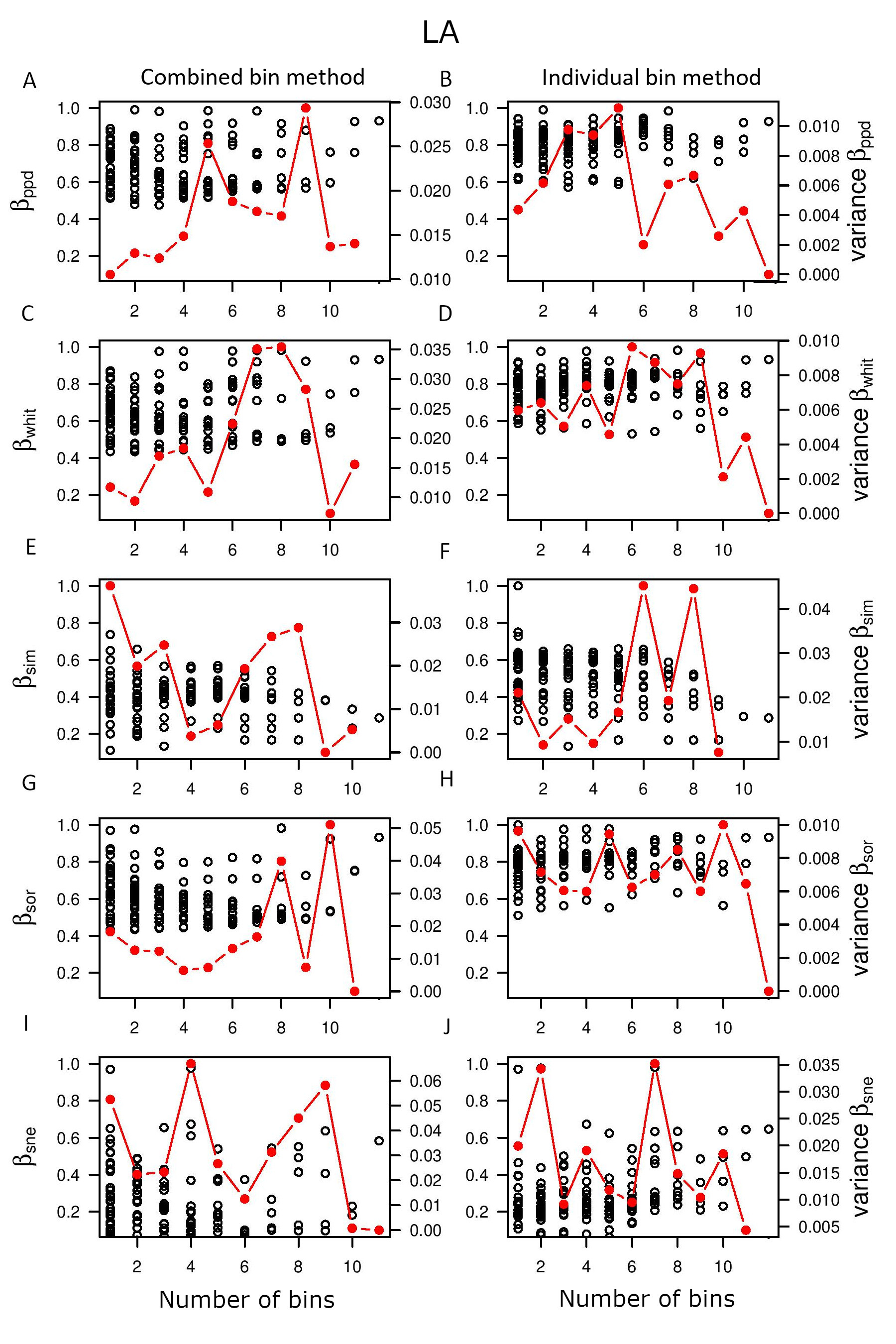

### Supplementary Figure (Figure S2)

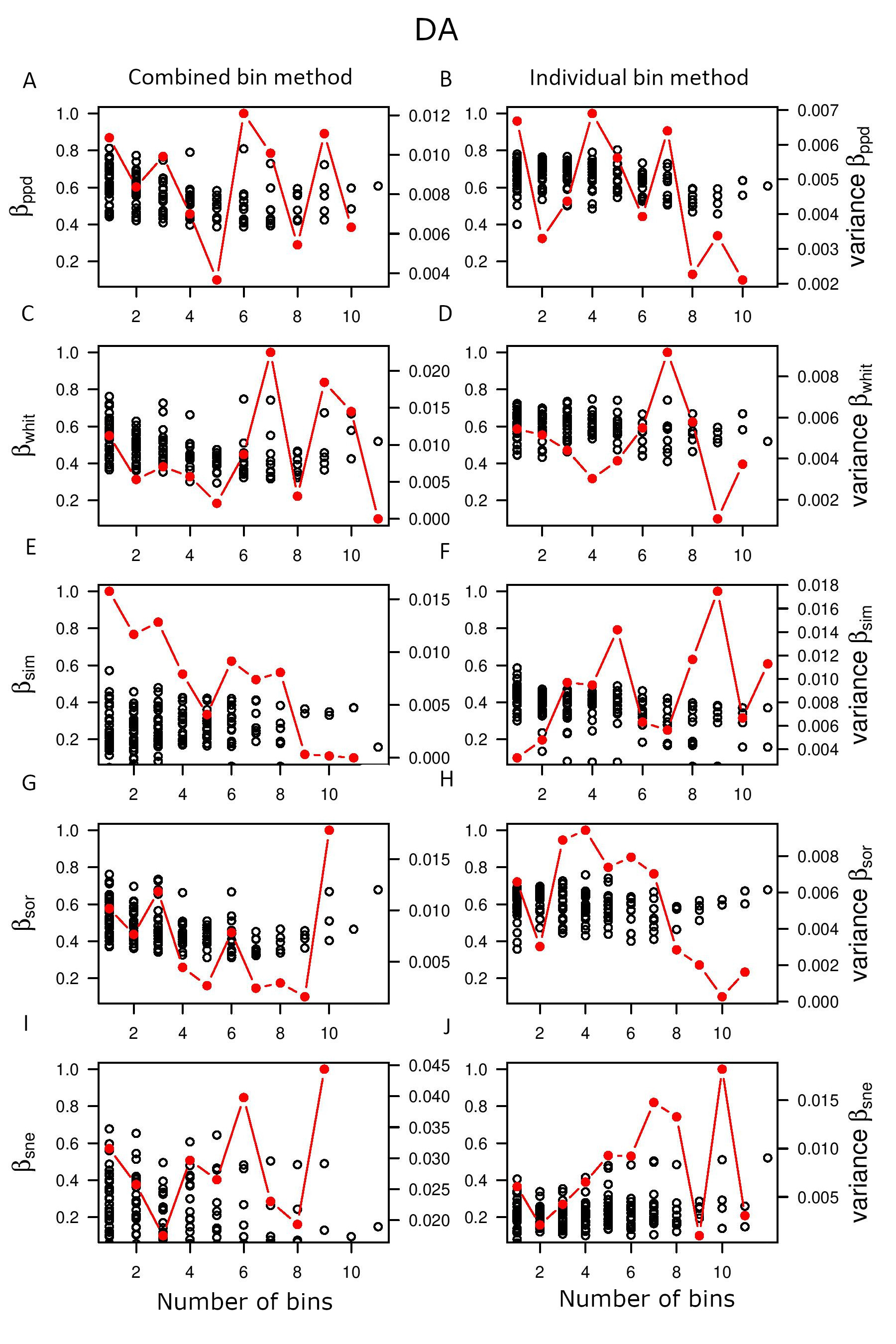
