## Supplementary R script (S3) for "Controls of spatial grain size and environmental variables on observed beta diversity of molluscan assemblage at a regional scale"

```
## ---------------------------
##
## Project: Effects of spatial grain sizes and environmental variables on beta
diversity of molluscan assemblage at a regional scale
##
## ## Author: Madhura Bhattacherjee, Devapriya Chattopadhyay
##
##
##
##
## Last Modified: 2022-10-30
##
## ---------------------------
##
##
##
##
## ---------------------------
require(vegan)
require(sads)
require(fields)
require(dplyr)
require(visreg)
require(ggplot2)
setwd("D:/Beta diversity Chapter")

###Beta diversity code for dead assemblage
master=read.csv("West Coast.csv", header=T)
ta <- tapply(master$Abundance,
list(master$Latitudinal.bin,master$Worms.updated.name), sum)
ta.0 <- ta
ta.0[is.na(ta)] <- 0
rownames(ta.
0)=c("21","20","19","18","17","16","15","14","13","12","11","10","9","8")

#####change the order of the rows from 8-21###############
require(dplyr)
ta.01=arrange(as.data.frame(ta.0), -row_number())
rownames(ta.
01)=c("8","9","10","11","12","13","14","15","16","17","18","19","20","21")

###change it to presence-absence matrix
ta.pa=decostand(ta.01,method = "pa")

##run the environment data
env<-read.delim("Environment.txt", header=T)
rownames(env)=c("21","20","19","18","17","16","15","14","13","12","11","10","9","8")
env[,1]=rownames(env)
env$cumulative= ave(env$Coastline.length..Km.,FUN=cumsum) ###calculating
cumulative coastlength


####calculating beta diversity using bray -curtis ######
#####PPD bray curtis#######
```

```
ta.0 <- as.matrix(ta.pa)
ta.prop <-prop.table(ta.0, margin=1) #change to proportions
ppd <- vegdist(ta.prop, method="bray") #distance matrix
mean(ppd) #mean beta diversity
dppd <- as.matrix(ppd)
dppd[dppd==0] <- NA #replace zeros with NA to calculate means and standard
error
locs <- apply(dppd, 2, FUN=mean, na.rm=T) #mean beta diversity per site with
regard to other sites #Tables 1,2

#####Null model in bray curtis########
###Combined bin method case 1#########
env<-read.delim("Environment.txt", header=T)
rownames(env)=c("21","20","19","18","17","16","15","14","13","12","11","10","9","8")

s=array(0,dim = c(3000,225))
g=array(0,dim = c(3000,225))
betadata=array(0,dim = c(2,225))
meanbeta=array(0,dim = c(3000,225))
veg=array(0,dim = c(3000,225))
meanppd=vector(mode = "integer", length =1)
env_mod=vector(mode = "integer", length =10)
subsamps <- list()
subsamps1<-list()

for(i in 1:50)
{
  env_mod=0
  subsamps1<-list()
  check=sample(1:14,2,replace = F)
  if(check[1]<check[2])
  {
    a=check[1]
    b=check[2]
    if(check[2]-check[1]==1)
    {
      betadata=ta.0[c(a,b),]
      ta.prop <-prop.table(betadata, margin=1)
      veg <- vegdist(ta.prop, method="bray")
      j=1
      x=(a+j)-1
      env_mod=env[x,9]
      out=data.frame(i,j,a,b,as.numeric(veg),x,env_mod)
      subsamps1[[i]] <-out
    }else{
      for ( j in 1:((b-a)-1))
      {
        r=a+j
        t=a+j+1
        s=colSums(ta.0[c(a:r),])
        if(t==b)
        {g=ta.0[t,]}else{
          g=colSums(ta.0[c(t:b),])
        }
```

```
        betadata=rbind(s,g)
        ta.prop <-prop.table(betadata, margin=1)
        veg <- vegdist(ta.prop, method="bray")
        x=(a+j)-1
        env_mod=env_mod+env[x,9]
        out=data.frame(i,j,a,b,as.numeric(veg),x,env_mod)
        subsamps1[[j]] <-out
      }
    }
  }else{
    a=check[2]
    b=check[1]
    if(check[1]-check[2]==1)
    {
      betadata=ta.0[c(a,b),]
      ta.prop <-prop.table(betadata, margin=1)
      veg <- vegdist(ta.prop, method="bray")
      j=1
      x=(a+j)-1
      env_mod=env[x,9]
      oute=data.frame(i,j,a,b,as.numeric(veg),x,env_mod)
      subsamps1[[j]] <-out
    }else{
      a=check[2]
      b=check[1]
      for ( j in 1:((b-a)-1))
      {
        r=a+j
        t=a+j+1
        s=colSums(ta.0[c(a:r),])
        if(t==b)
        {g=ta.0[t,]}else{
          g=colSums(ta.0[c(t:b),])
        }
        betadata=rbind(s,g)
        ta.prop <-prop.table(betadata, margin=1)
        veg <- vegdist(ta.prop, method="bray")
        x=(a+j)-1
        env_mod=env_mod+env[x,9]
        out=data.frame(i,j,a,b,as.numeric(veg),x,env_mod)
        subsamps1[[j]] <-out
      }
    }
  }
  subsamples1 <- do.call("rbind",subsamps1)
  subsamps[[i]]=subsamples1
  subsamples_ppd2<- do.call("rbind",subsamps)
}

####calculating variance for bins########
varbeta_ppd=tapply(subsamples_ppd2$as.numeric.veg., subsamples_ppd2$j, var)

######create bins of coastline length#####
#write.csv(subsamples_ppd2,"subs_ppd_case1.csv")
```

```
library(dplyr)
library(Hmisc)
subsamples<-subsamples_ppd2%>%mutate(Coastlinebins =
cut(subsamples_ppd2$env_mod, breaks =
c(0,200,400,600,800,1000,1200,1400,1600,1800,2000,2200)))
head(subsamples_ppd2,200)
varbeta=tapply(subsamples$as.numeric.veg., subsamples$Coastlinebins, var)
Coastlinebins=c("200","400","600","800","1000","1200","1400","1600","1800","2000","2200")
var=cbind(Coastline.bins=as.numeric(Coastlinebins),varbeta=as.numeric(round(varbeta,
4)))
variance_ppd=as.data.frame(var)

#######Individual bin method case 2#########################
env<-read.delim("Environment.txt", header=T)
row.names(env)=env[,1]# Declare column 1 as the row names#

s=array(0,dim = c(3000,225))
g=array(0,dim = c(3000,225))
betadata=array(0,dim = c(2,225))
meanbeta=array(0,dim = c(3000,225))
veg=array(0,dim = c(3000,225))
meanppd=vector(mode = "integer", length =1)
env_mod=vector(mode = "integer", length =10)
subsamps <- list()
subsamps<-list()

for(i in 1:50)
{
  env_mod=0
  subsamps1<-list()
  check=sample(1:14,2,replace = F)
  if(check[1]<check[2])
  {
    a=check[1]
    b=check[2]
    if(check[2]-check[1]==1)
    {
      betadata=ta.0[c(a,b),]
      ta.prop <-prop.table(betadata, margin=1)
      veg <- vegdist(ta.prop, method="bray")
      meanbeta<-veg
      j=1
      x=(a+j)-1
      env_mod=env[x,9]
      out=data.frame(i,j,a,b,as.numeric(meanbeta),x,env_mod)
      subsamps1[[i]] <-out
    }else{
      for ( j in 1:((b-a)-1))
      {
        r=a+j
        t=a+j+1
        s=colSums(ta.0[c(a:r),])
        if(t==b)
        {g=ta.0[t,]}else{
```

```
          g=ta.0[c(t:b),]
        }
        betadata=rbind(s,g)
        ta.prop <-prop.table(betadata, margin=1)
        veg <- vegdist(ta.prop, method="bray")
        meanbeta=mean(veg)
        x=(a+j)-1
        env_mod=env_mod+env[x,9]
        out=data.frame(i,j,a,b,as.numeric(meanbeta),x,env_mod)
        subsamps1[[j]] <-out
      }
    }
  }else{
    a=check[2]
    b=check[1]
    if(check[1]-check[2]==1)
    {
      betadata=ta.0[c(a,b),]
      ta.prop <-prop.table(betadata, margin=1)
      veg <- vegdist(ta.prop, method="bray")
      meanbeta<-veg
      j=1
      x=(a+j)-1
      env_mod=env[x,9]
      oute=data.frame(i,j,a,b,as.numeric(meanbeta),x,env_mod)
      subsamps1[[j]] <-out
    }else{
      a=check[2]
      b=check[1]
      for ( j in 1:((b-a)-1))
      {
        r=a+j
        t=a+j+1
        s=colSums(ta.0[c(a:r),])
        if(t==b)
        {g=ta.0[t,]}else{
          g=ta.0[c(t:b),]
        }
        betadata=rbind(s,g)
        ta.prop <-prop.table(betadata, margin=1)
        veg <- vegdist(ta.prop, method="bray")
        meanbeta=mean(veg)
        x=(a+j)-1
        env_mod=env_mod+env[x,9]
        out=data.frame(i,j,a,b,as.numeric(meanbeta),x,env_mod)
        subsamps1[[j]] <-out
      }
    }
  }
  subsamples1 <- do.call("rbind",subsamps1)
  subsamps[[i]]=subsamples1
  subsamples_ppd3<- do.call("rbind",subsamps)
}
```

```
####variance for bins####
varbeta_ppd3=tapply(subsamples_ppd3$as.numeric.meanbeta., subsamples_ppd3$j,
var)

######create bins of coastline length#####
#write.csv(subsamples_ppd3,"subs_ppd_case2.csv")
library(dplyr)
subsamples<-subsamples_ppd3%>%mutate(Coastlinebins =
cut(subsamples_ppd3$env_mod, breaks =
c(0,200,400,600,800,1000,1200,1400,1600,1800,2000,2200)))
head(subsamples,200)
varbeta=tapply(subsamples$as.numeric.meanbeta., subsamples$Coastlinebins, var)
Coastlinebins=c("200","400","600","800","1000","1200","1400","1600","1800","2000","2200")
var=cbind(Coastline.bins=as.numeric(Coastlinebins),varbeta=as.numeric(round(varbeta,
4)))
variance2_ppd=as.data.frame(var)



##########################################################################
#####Beta diversity whittaker index############
require(vegan)
ta.pa=decostand(ta.01,method = "pa")
d <- betadiver(ta.pa, "w") ##whittaker index####
dppd1<- as.matrix(d)
as.dist(dppd1)
mean(d)
locs1 <- apply(dppd1, 2, FUN=mean, na.rm=T)

####Nullmodel whittaker index####################
#####Combined bin method####################

env<-read.delim("Environment.txt", header=T)
rownames(env)=c("21","20","19","18","17","16","15","14","13","12","11","10","9","8")

#row.names(env)=env[,1]# Declare column 1 as the row names#
s=array(0,dim = c(3000,225))
g=array(0,dim = c(3000,225))
betadata=array(0,dim = c(2,225))
meanbeta=array(0,dim = c(3000,225))
veg=array(0,dim = c(3000,225))
meanppd=vector(mode = "integer", length =1)
env_mod=vector(mode = "integer", length =10)
subsamps <- list()
subsamps<-list()

for(i in 1:50)
{
  env_mod=0
  subsamps1<-list()
  check=sample(1:14,2,replace = F)
  if(check[1]<check[2])
  {
    a=check[1]
```

```
    b=check[2]
    if(check[2]-check[1]==1)
    {
      betadata=ta.0[c(a,b),]
      d <- betadiver(betadata, "w")##whittaker index####
      veg <- as.dist(d)
      j=1
      x=(a+j)-1
      env_mod=env[x,9]
      out=data.frame(i,j,a,b,as.numeric(veg),x,env_mod)
      subsamps1[[i]] <-out
    }else{
      for ( j in 1:((b-a)-1))
      {
        r=a+j
        t=a+j+1
        s=colSums(ta.0[c(a:r),])
        if(t==b)
        {g=ta.0[t,]}else{
          g=colSums(ta.0[c(t:b),])
        }
        betadata=rbind(s,g)
        d <- betadiver(betadata, "w")##whittaker index####
        veg <- as.dist(d)
        x=(a+j)-1
        env_mod=env_mod+env[x,9]
        out=data.frame(i,j,a,b,as.numeric(veg),x,env_mod)
        subsamps1[[j]] <-out
      }
    }
  }else{
    a=check[2]
    b=check[1]
    if(check[1]-check[2]==1)
    {
      betadata=ta.0[c(a,b),]
      d <- betadiver(betadata, "w")##whittaker index####
      veg <- as.dist(d)
      j=1
      x=(a+j)-1
      env_mod=env[x,9]
      oute=data.frame(i,j,a,b,as.numeric(veg),x,env_mod)
      subsamps1[[j]] <-out
    }else{
      a=check[2]
      b=check[1]
      for ( j in 1:((b-a)-1))
      {
        r=a+j
        t=a+j+1
        s=colSums(ta.0[c(a:r),])
        if(t==b)
        {g=ta.0[t,]}else{
          g=colSums(ta.0[c(t:b),])
```

```
        }
        betadata=rbind(s,g)
        d <- betadiver(betadata, "w")##whittaker index####
        veg <- as.dist(d)
        x=(a+j)-1
        env_mod=env_mod+env[x,9]
        out=data.frame(i,j,a,b,as.numeric(veg),x,env_mod)
        subsamps1[[j]] <-out
      }
    }
  }
  subsamples1 <- do.call("rbind",subsamps1)
  subsamps[[i]]=subsamples1
  subsamples_whit2 <- do.call("rbind",subsamps)
}

###variance for bins##########
varbeta_whit1=tapply(subsamples_whit2$as.numeric.veg., subsamples_whit2$j,
var)

######create bins of coastline length#####
#write.csv(subsamples_whit2,"subs_whit_case1.csv")
library(dplyr)
subsamples_whit2<-subsamples_whit2%>%mutate(Coastlinebins =
cut(subsamples_whit2$env_mod, breaks =
c(0,200,400,600,800,1000,1200,1400,1600,1800,2000,2200)))
head(subsamples_whit2,200)
varbeta=tapply(subsamples_whit2$as.numeric.veg.,
subsamples_whit2$Coastlinebins, var)
write.table(subsamples_whit2$Coastlinebins)
Coastlinebins=c("200","400","600","800","1000","1200","1400","1600","1800","2000","2200")
var=cbind(Coastline.bins=as.numeric(Coastlinebins),varbeta=as.numeric(round(varbeta,
4)))
variance_whit=as.data.frame(var)

####Individual bin method case 2#########
########################clubbing rows from a side only and keeping the
remaining rows##########################################################

env<-read.delim("Environment.txt", header=T)
row.names(env)=env[,1]# Declare column 1 as the row names#

s=array(0,dim = c(3000,225))
g=array(0,dim = c(3000,225))
betadata=array(0,dim = c(2,225))
meanbeta=array(0,dim = c(3000,225))
veg=array(0,dim = c(3000,225))
meanppd=vector(mode = "integer", length =1)
env_mod=vector(mode = "integer", length =10)
subsamps <- list()
subsamps<-list()

for(i in 1:50)
{
```

```
env_mod=0
subsamps1<-list()
check=sample(1:14,2,replace = F)
if(check[1]<check[2])
{
  a=check[1]
  b=check[2]
  if(check[2]-check[1]==1)
  {
    betadata=ta.0[c(a,b),]
    d <- betadiver(betadata, "w")##whittaker index####
    veg <- as.dist(d)
    meanbeta<-veg
    j=1
    x=(a+j)-1
    env_mod=env[x,9]
    out=data.frame(i,j,a,b,as.numeric(meanbeta),x,env_mod)
    subsamps1[[i]] <-out
  }else{
    for ( j in 1:((b-a)-1))
    {
      r=a+j
      t=a+j+1
      s=colSums(ta.0[c(a:r),])
      if(t==b)
      {g=ta.0[t,]}else{
        g=ta.0[c(t:b),]
      }
      betadata=rbind(s,g)
      d <- betadiver(betadata, "w")##whittaker index####
      veg <- as.dist(d)
      meanbeta=mean(veg)
      x=(a+j)-1
      env_mod=env_mod+env[x,9]
      out=data.frame(i,j,a,b,as.numeric(meanbeta),x,env_mod)
      subsamps1[[j]] <-out
    }
  }
}else{
  a=check[2]
  b=check[1]
  if(check[1]-check[2]==1)
  {
    betadata=ta.0[c(a,b),]
    d <- betadiver(betadata, "w")##whittaker index####
    veg <- as.dist(d)
    meanbeta<-veg
    j=1
    x=(a+j)-1
    env_mod=env[x,9]
    oute=data.frame(i,j,a,b,as.numeric(meanbeta),x,env_mod)
    subsamps1[[j]] <-out
  }else{
    a=check[2]
```

```
      b=check[1]
      for ( j in 1:((b-a)-1))
      {
        r=a+j
        t=a+j+1
        s=colSums(ta.0[c(a:r),])
        if(t==b)
        {g=ta.0[t,]}else{
          g=ta.0[c(t:b),]
        }
        betadata=rbind(s,g)
        d <- betadiver(betadata, "w")##whittaker index####
        veg <- as.dist(d)
        meanbeta=mean(veg)
        x=(a+j)-1
        env_mod=env_mod+env[x,9]
        out=data.frame(i,j,a,b,as.numeric(meanbeta),x,env_mod)
        subsamps1[[j]] <-out
      }
    }
  }
  subsamples1 <- do.call("rbind",subsamps1)
  subsamps[[i]]=subsamples1
  subsamples_whit3 <- do.call("rbind",subsamps)
}


varbeta_whit2=tapply(subsamples_whit3$as.numeric.meanbeta.,
subsamples_whit3$j, var)

######create bins of coastline length#####
library(dplyr)
subsamples_whit3<-subsamples_whit3%>%mutate(Coastlinebins =
cut(subsamples_whit3$env_mod, breaks =
c(0,200,400,600,800,1000,1200,1400,1600,1800,2000,2200)))
head(subsamples_whit3,200)
varbeta=tapply(subsamples_whit3$as.numeric.meanbeta.,
subsamples_whit3$Coastlinebins, var)

Coastlinebins=c("200","400","600","800","1000","1200","1400","1600","1800","2000","2200")
var=cbind(Coastline.bins=as.numeric(Coastlinebins),varbeta=as.numeric(round(varbeta,
4)))

variance2_whit=as.data.frame(var)

##############################################
# pairwise beta diversities####
#####simpson######turnover component########
library(betapart)
ta.pa=decostand(ta.01,method = "pa")
pair.s <- beta.pair(ta.pa)
b_sim <- as.matrix(pair.s$beta.sim)
as.dist(b_sim)
mean(b_sim)
```

```
locs_sim <- apply(b_sim, 2, FUN=mean, na.rm=T) #mean beta diversity per site
with regard to other sites #Tables 1,2


#####Null model################
###Combined bin method########
env<-read.delim("Environment.txt", header=T)
rownames(env)=c("21","20","19","18","17","16","15","14","13","12","11","10","9","8")

s=array(0,dim = c(3000,225))
g=array(0,dim = c(3000,225))
betadata=array(0,dim = c(2,225))
meanbeta=array(0,dim = c(3000,225))
veg=array(0,dim = c(3000,225))
meanppd=vector(mode = "integer", length =1)
env_mod=vector(mode = "integer", length =10)
subsamps <- list()
subsamps<-list()

for(i in 1:50)
{
  env_mod=0
  subsamps1<-list()
  check=sample(1:14,2,replace = F)
  if(check[1]<check[2])
  {
    a=check[1]
    b=check[2]
    if(check[2]-check[1]==1)
    {
      betadata=ta.0[c(a,b),]
      betadata.pa=decostand(betadata,method = "pa")
      pair.s <- beta.pair(betadata.pa)
      b_sim <- as.matrix(pair.s$beta.sim)
      veg=as.dist(b_sim)
      j=1
      x=(a+j)-1
      env_mod=env[x,9]
      out=data.frame(i,j,a,b,as.numeric(veg),x,env_mod)
      subsamps1[[i]] <-out
    }else{
      for ( j in 1:((b-a)-1))
      {
        r=a+j
        t=a+j+1
        s=colSums(ta.0[c(a:r),])
        if(t==b)
        {g=ta.0[t,]}else{
          g=colSums(ta.0[c(t:b),])
        }
        betadata=rbind(s,g)
        betadata.pa=decostand(betadata,method = "pa")
        pair.s <- beta.pair(betadata.pa)
        b_sim <- as.matrix(pair.s$beta.sim)
```

```
        veg=as.dist(b_sim)
        x=(a+j)-1
        env_mod=env_mod+env[x,9]
        out=data.frame(i,j,a,b,as.numeric(veg),x,env_mod)
        subsamps1[[j]] <-out
      }
    }
  }else{
    a=check[2]
    b=check[1]
    if(check[1]-check[2]==1)
    {
      betadata=ta.0[c(a,b),]
      betadata.pa=decostand(betadata,method = "pa")
      pair.s <- beta.pair(betadata.pa)
      b_sim <- as.matrix(pair.s$beta.sim)
      veg=as.dist(b_sim)
      j=1
      x=(a+j)-1
      env_mod=env[x,9]
      oute=data.frame(i,j,a,b,as.numeric(veg),x,env_mod)
      subsamps1[[j]] <-out
    }else{
      a=check[2]
      b=check[1]
      for ( j in 1:((b-a)-1))
      {
        r=a+j
        t=a+j+1
        s=colSums(ta.0[c(a:r),])
        if(t==b)
        {g=ta.0[t,]}else{
          g=colSums(ta.0[c(t:b),])
        }
        betadata=rbind(s,g)
        betadata.pa=decostand(betadata,method = "pa")
        pair.s <- beta.pair(betadata.pa)
        b_sim <- as.matrix(pair.s$beta.sim)
        veg=as.dist(b_sim)
        x=(a+j)-1
        env_mod=env_mod+env[x,9]
        out=data.frame(i,j,a,b,as.numeric(veg),x,env_mod)
        subsamps1[[j]] <-out
      }
    }
  }
  subsamples1 <- do.call("rbind",subsamps1)
  subsamps[[i]]=subsamples1
  subsamples_simp2 <- do.call("rbind",subsamps)
}


####variance for bins#####
```

```
varbeta_simp1=tapply(subsamples_simp2$as.numeric.veg., subsamples_simp2$j,
var)

######create bins of coastline length#####
library(dplyr)
#write.csv(subsamples_simp2,"subs_sim_case1.csv")
subsamples_simp2<-subsamples_simp2%>%mutate(Coastlinebins =
cut(subsamples_simp2$env_mod, breaks =
c(0,200,400,600,800,1000,1200,1400,1600,1800,2000,2200)))
head(subsamples_simp2,200)
varbeta=tapply(subsamples_simp2$as.numeric.veg.,
subsamples_simp2$Coastlinebins, var)
Coastlinebins=c("200","400","600","800","1000","1200","1400","1600","1800","2000","2200")
var=cbind(Coastline.bins=as.numeric(Coastlinebins),varbeta=as.numeric(round(varbeta,
4)))
variance_simp=as.data.frame(var)

#####Individual bin method########################
########################clubbing rows from a side only and keeping the
remaining rows#########################################################

env<-read.delim("Environment.txt", header=T)
row.names(env)=env[,1]# Declare column 1 as the row names#

s=array(0,dim = c(3000,225))
g=array(0,dim = c(3000,225))
betadata=array(0,dim = c(2,225))
meanbeta=array(0,dim = c(3000,225))
veg=array(0,dim = c(3000,225))
meanppd=vector(mode = "integer", length =1)
env_mod=vector(mode = "integer", length =10)
subsamps <- list()
subsamps<-list()

for(i in 1:50)
{
  env_mod=0
  subsamps1<-list()
  check=sample(1:14,2,replace = F)
  if(check[1]<check[2])
  {
    a=check[1]
    b=check[2]
    if(check[2]-check[1]==1)
    {
      betadata=ta.0[c(a,b),]
      betadata.pa=decostand(betadata,method = "pa")
      pair.s <- beta.pair(betadata.pa)
      b_sim <- as.matrix(pair.s$beta.sim)
      meanbeta=as.dist(b_sim)
      j=1
      x=(a+j)-1
      env_mod=env[x,9]
      out=data.frame(i,j,a,b,as.numeric(meanbeta),x,env_mod)
```

```
      subsamps1[[i]] <-out
    }else{
      for ( j in 1:((b-a)-1))
      {
        r=a+j
        t=a+j+1
        s=colSums(ta.0[c(a:r),])
        if(t==b)
        {g=ta.0[t,]}else{
          g=ta.0[c(t:b),]
        }
        betadata=rbind(s,g)
        betadata.pa=decostand(betadata,method = "pa")
        pair.s <- beta.pair(betadata.pa)
        b_sim <- as.matrix(pair.s$beta.sim)
        veg=as.dist(b_sim)
        meanbeta=mean(veg)
        x=(a+j)-1
        env_mod=env_mod+env[x,9]
        out=data.frame(i,j,a,b,as.numeric(meanbeta),x,env_mod)
        subsamps1[[j]] <-out
      }
    }
  }else{
    a=check[2]
    b=check[1]
    if(check[1]-check[2]==1)
    {
      betadata=ta.0[c(a,b),]
      betadata.pa=decostand(betadata,method = "pa")
      pair.s <- beta.pair(betadata.pa)
      b_sim <- as.matrix(pair.s$beta.sim)
      meanbeta=as.dist(b_sim)
      j=1
      x=(a+j)-1
      env_mod=env[x,9]
      oute=data.frame(i,j,a,b,as.numeric(meanbeta),x,env_mod)
      subsamps1[[j]] <-out
    }else{
      a=check[2]
      b=check[1]
      for ( j in 1:((b-a)-1))
      {
        r=a+j
        t=a+j+1
        s=colSums(ta.0[c(a:r),])
        if(t==b)
        {g=ta.0[t,]}else{
          g=ta.0[c(t:b),]
        }
        betadata=rbind(s,g)
        betadata.pa=decostand(betadata,method = "pa")
        pair.s <- beta.pair(betadata.pa)
        b_sim <- as.matrix(pair.s$beta.sim)
```

```
        veg=as.dist(b_sim)
        meanbeta=mean(veg)
        x=(a+j)-1
        env_mod=env_mod+env[x,9]
        out=data.frame(i,j,a,b,as.numeric(meanbeta),x,env_mod)
        subsamps1[[j]] <-out
      }
    }
  }
  subsamples1 <- do.call("rbind",subsamps1)
  subsamps[[i]]=subsamples1
  subsamples_simp3 <- do.call("rbind",subsamps)
}


varbeta_simp2=tapply(subsamples_simp3$as.numeric.meanbeta.,
subsamples_simp3$j, var)

######create bins of coastline length#####
#write.csv(subsamples_simp3,"subs_sim_case2.csv")
library(dplyr)
subsamples_simp3<-subsamples_simp3%>%mutate(Coastlinebins =
cut(subsamples_simp3$env_mod, breaks =
c(0,200,400,600,800,1000,1200,1400,1600,1800,2000,2200)))
head(subsamples_simp3,200)
varbeta=tapply(subsamples_simp3$as.numeric.meanbeta.,
subsamples_simp3$Coastlinebins, var)
Coastlinebins=c("200","400","600","800","1000","1200","1400","1600","1800","2000","2200")
var=cbind(Coastline.bins=as.numeric(Coastlinebins),varbeta=as.numeric(round(varbeta,
4)))
variance2=as.data.frame(var)


#############################################################
####################Sorenson#########total dissimilarity
library(betapart)
ta.pa=decostand(ta.01,method = "pa")
pair.s <- beta.pair(ta.pa)
b_sor<- as.matrix(pair.s$beta.sor)
mean(b_sor)
locs_sor <- apply(b_sor, 2, FUN=mean, na.rm=T) #mean beta diversity per site
with regard to other sites


######Null model##############
##########Combined bin method ###################

env<-read.delim("Environment.txt", header=T)
rownames(env)=c("21","20","19","18","17","16","15","14","13","12","11","10","9","8")

s=array(0,dim = c(3000,225))
g=array(0,dim = c(3000,225))
betadata=array(0,dim = c(2,225))
meanbeta=array(0,dim = c(3000,225))
```

```
veg=array(0,dim = c(3000,225))
meanppd=vector(mode = "integer", length =1)
env_mod=vector(mode = "integer", length =10)
subsamps <- list()
subsamps<-list()

for(i in 1:50)
{
  env_mod=0
  subsamps1<-list()
  check=sample(1:14,2,replace = F)
  if(check[1]<check[2])
  {
    a=check[1]
    b=check[2]
    if(check[2]-check[1]==1)
    {
      betadata=ta.0[c(a,b),]
      betadata.pa=decostand(betadata,method = "pa")
      pair.s <- beta.pair(betadata.pa)
      b_sor <- as.matrix(pair.s$beta.sor)
      veg=as.dist(b_sor)
      j=1
      x=(a+j)-1
      env_mod=env[x,9]
      out=data.frame(i,j,a,b,as.numeric(veg),x,env_mod)
      subsamps1[[i]] <-out
    }else{
      for ( j in 1:((b-a)-1))
      {
        r=a+j
        t=a+j+1
        s=colSums(ta.0[c(a:r),])
        if(t==b)
        {g=ta.0[t,]}else{
          g=colSums(ta.0[c(t:b),])
        }
        betadata=rbind(s,g)
        betadata.pa=decostand(betadata,method = "pa")
        pair.s <- beta.pair(betadata.pa)
        b_sor <- as.matrix(pair.s$beta.sor)
        veg=as.dist(b_sor)
        x=(a+j)-1
        env_mod=env_mod+env[x,9]
        out=data.frame(i,j,a,b,as.numeric(veg),x,env_mod)
        subsamps1[[j]] <-out
      }
    }
  }else{
    a=check[2]
    b=check[1]
    if(check[1]-check[2]==1)
    {
      betadata=ta.0[c(a,b),]
```

```
      betadata.pa=decostand(betadata,method = "pa")
      pair.s <- beta.pair(betadata.pa)
      b_sor<- as.matrix(pair.s$beta.sor)
      veg=as.dist(b_sor)
      j=1
      x=(a+j)-1
      env_mod=env[x,9]
      oute=data.frame(i,j,a,b,as.numeric(veg),x,env_mod)
      subsamps1[[j]] <-out
    }else{
      a=check[2]
      b=check[1]
      for ( j in 1:((b-a)-1))
      {
        r=a+j
        t=a+j+1
        s=colSums(ta.0[c(a:r),])
        if(t==b)
        {g=ta.0[t,]}else{
          g=colSums(ta.0[c(t:b),])
        }
        betadata=rbind(s,g)
        betadata.pa=decostand(betadata,method = "pa")
        pair.s <- beta.pair(betadata.pa)
        b_sor <- as.matrix(pair.s$beta.sor)
        veg=as.dist(b_sor)
        x=(a+j)-1
        env_mod=env_mod+env[x,9]
        out=data.frame(i,j,a,b,as.numeric(veg),x,env_mod)
        subsamps1[[j]] <-out
      }
    }
  }
  subsamples1 <- do.call("rbind",subsamps1)
  subsamps[[i]]=subsamples1
  subsamples_sor2 <- do.call("rbind",subsamps)
}


varbeta_sor=tapply(subsamples_sor2$as.numeric.veg., subsamples_sor2$j, var)

######create bins of coastline length#####
#write.csv(subsamples_sor2,"subs_sor_case1.csv")
library(dplyr)
subsamples_sor2<-subsamples_sor2%>%mutate(Coastlinebins =
cut(subsamples_sor2$env_mod, breaks =
c(0,200,400,600,800,1000,1200,1400,1600,1800,2000,2200)))
head(subsamples_sor2,200)
varbeta=tapply(subsamples_sor2$as.numeric.veg., subsamples_sor2$Coastlinebins,
var)
Coastlinebins=c("200","400","600","800","1000","1200","1400","1600","1800","2000","2200")
var=cbind(Coastline.bins=as.numeric(Coastlinebins),varbeta=as.numeric(round(varbeta,
4)))
variance_sor=as.data.frame(var)
```

```
#######Individual bin method#####################################
########################clubbing rows from a side only and keeping the
remaining rows#########################################################

env<-read.delim("Environment.txt", header=T)
row.names(env)=env[,1]

s=array(0,dim = c(3000,225))
g=array(0,dim = c(3000,225))
betadata=array(0,dim = c(2,225))
meanbeta=array(0,dim = c(3000,225))
veg=array(0,dim = c(3000,225))
meanppd=vector(mode = "integer", length =1)
env_mod=vector(mode = "integer", length =10)
subsamps <- list()
subsamps<-list()

for(i in 1:50)
{
  env_mod=0
  subsamps1<-list()
  check=sample(1:14,2,replace = F)
  if(check[1]<check[2])
  {
    a=check[1]
    b=check[2]
    if(check[2]-check[1]==1)
    {
      betadata=ta.0[c(a,b),]
      betadata.pa=decostand(betadata,method = "pa")
      pair.s <- beta.pair(betadata.pa)
      b_sor <- as.matrix(pair.s$beta.sor)
      meanbeta=as.dist(b_sor)
      j=1
      x=(a+j)-1
      env_mod=env[x,9]
      out=data.frame(i,j,a,b,as.numeric(meanbeta),x,env_mod)
      subsamps1[[i]] <-out
    }else{
      for ( j in 1:((b-a)-1))
      {
        r=a+j
        t=a+j+1
        s=colSums(ta.0[c(a:r),])
        if(t==b)
        {g=ta.0[t,]}else{
          g=ta.0[c(t:b),]
        }
        betadata=rbind(s,g)
        betadata.pa=decostand(betadata,method = "pa")
        pair.s <- beta.pair(betadata.pa)
        b_sor <- as.matrix(pair.s$beta.sor)
        veg=as.dist(b_sor)
```

```
        meanbeta=mean(veg)
        x=(a+j)-1
        env_mod=env_mod+env[x,9]
        out=data.frame(i,j,a,b,as.numeric(meanbeta),x,env_mod)
        subsamps1[[j]] <-out
      }
    }
  }else{
    a=check[2]
    b=check[1]
    if(check[1]-check[2]==1)
    {
      betadata=ta.0[c(a,b),]
      betadata.pa=decostand(betadata,method = "pa")
      pair.s <- beta.pair(betadata.pa)
      b_sor <- as.matrix(pair.s$beta.sor)
      meanbeta=as.dist(b_sor)
      j=1
      x=(a+j)-1
      env_mod=env[x,9]
      oute=data.frame(i,j,a,b,as.numeric(meanbeta),x,env_mod)
      subsamps1[[j]] <-out
    }else{
      a=check[2]
      b=check[1]
      for ( j in 1:((b-a)-1))
      {
        r=a+j
        t=a+j+1
        s=colSums(ta.0[c(a:r),])
        if(t==b)
        {g=ta.0[t,]}else{
          g=ta.0[c(t:b),]
        }
        betadata=rbind(s,g)
        betadata.pa=decostand(betadata,method = "pa")
        pair.s <- beta.pair(betadata.pa)
        b_sor <- as.matrix(pair.s$beta.sor)
        veg=as.dist(b_sor)
        meanbeta=mean(veg)
        x=(a+j)-1
        env_mod=env_mod+env[x,9]
        out=data.frame(i,j,a,b,as.numeric(meanbeta),x,env_mod)
        subsamps1[[j]] <-out
      }
    }
  }
  subsamples1 <- do.call("rbind",subsamps1)
  subsamps[[i]]=subsamples1
  subsamples_sor3 <- do.call("rbind",subsamps)
}
```

```
varbeta_sor2=tapply(subsamples_sor3$as.numeric.meanbeta., subsamples_sor3$j,
var)

######create bins of coastline length#####
library(dplyr)
#write.csv(subsamples_sor3,"subsamples_sor_case2.csv")
subsamples_sor3<-subsamples_sor3%>%mutate(Coastlinebins =
cut(subsamples_sor3$env_mod, breaks =
c(0,200,400,600,800,1000,1200,1400,1600,1800,2000,2200)))
head(subsamples_sor3,200)
varbeta=tapply(subsamples_sor3$as.numeric.meanbeta.,
subsamples_sor3$Coastlinebins, var)
Coastlinebins=c("200","400","600","800","1000","1200","1400","1600","1800","2000","2200")
var=cbind(Coastline.bins=as.numeric(Coastlinebins),varbeta=as.numeric(round(varbeta,
4)))
variance2_sor=as.data.frame(var)

#################nestedness component of sorenson###########
library(betapart)
library(vegan)
ta.pa=decostand(ta.01,method = "pa")
pair.s <- beta.pair(ta.pa)
b_sne<- as.matrix(pair.s$beta.sne)
mean(b_sne)
locs_sne <- apply(b_sne, 2, FUN=mean, na.rm=T) #mean beta diversity per site
with regard to other sites #Tables 1,2
x1 = matrix(c(rownames(ta.01)),nrow=14,ncol=1)


####Null model##############
##########Combined bin method################
env<-read.delim("Environment.txt", header=T)
rownames(env)=c("21","20","19","18","17","16","15","14","13","12","11","10","9","8")

s=array(0,dim = c(3000,225))
g=array(0,dim = c(3000,225))
betadata=array(0,dim = c(2,225))
meanbeta=array(0,dim = c(3000,225))
veg=array(0,dim = c(3000,225))
meanppd=vector(mode = "integer", length =1)
env_mod=vector(mode = "integer", length =10)
subsamps <- list()
subsamps<-list()

for(i in 1:50)
{
  env_mod=0
  subsamps1<-list()
  check=sample(1:14,2,replace = F)
  if(check[1]<check[2])
  {
    a=check[1]
    b=check[2]
    if(check[2]-check[1]==1)
```

```
    {
      betadata=ta.0[c(a,b),]
      betadata.pa=decostand(betadata,method = "pa")
      pair.s <- beta.pair(betadata.pa)
      b_sne <- as.matrix(pair.s$beta.sne)
      veg=as.dist(b_sne)
      j=1
      x=(a+j)-1
      env_mod=env[x,9]
      out=data.frame(i,j,a,b,as.numeric(veg),x,env_mod)
      subsamps1[[i]] <-out
    }else{
      for ( j in 1:((b-a)-1))
      {
        r=a+j
        t=a+j+1
        s=colSums(ta.0[c(a:r),])
        if(t==b)
        {g=ta.0[t,]}else{
          g=colSums(ta.0[c(t:b),])
        }
        betadata=rbind(s,g)
        betadata.pa=decostand(betadata,method = "pa")
        pair.s <- beta.pair(betadata.pa)
        b_sne <- as.matrix(pair.s$beta.sne)
        veg=as.dist(b_sne)
        x=(a+j)-1
        env_mod=env_mod+env[x,9]
        out=data.frame(i,j,a,b,as.numeric(veg),x,env_mod)
        subsamps1[[j]] <-out
      }
    }
  }else{
    a=check[2]
    b=check[1]
    if(check[1]-check[2]==1)
    {
      betadata=ta.0[c(a,b),]
      betadata.pa=decostand(betadata,method = "pa")
      pair.s <- beta.pair(betadata.pa)
      b_sne<- as.matrix(pair.s$beta.sne)
      veg=as.dist(b_sne)
      j=1
      x=(a+j)-1
      env_mod=env[x,9]
      oute=data.frame(i,j,a,b,as.numeric(veg),x,env_mod)
      subsamps1[[j]] <-out
    }else{
      a=check[2]
      b=check[1]
      for ( j in 1:((b-a)-1))
      {
        r=a+j
        t=a+j+1
```

```
        s=colSums(ta.0[c(a:r),])
        if(t==b)
        {g=ta.0[t,]}else{
          g=colSums(ta.0[c(t:b),])
        }
        betadata=rbind(s,g)
        betadata.pa=decostand(betadata,method = "pa")
        pair.s <- beta.pair(betadata.pa)
        b_sne <- as.matrix(pair.s$beta.sne)
        veg=as.dist(b_sne)
        x=(a+j)-1
        env_mod=env_mod+env[x,9]
        out=data.frame(i,j,a,b,as.numeric(veg),x,env_mod)
        subsamps1[[j]] <-out
      }
    }
  }
  subsamples1 <- do.call("rbind",subsamps1)
  subsamps[[i]]=subsamples1
  subsamples_sne2 <- do.call("rbind",subsamps)
}



varbeta_sne=tapply(subsamples_sne2$as.numeric.veg., subsamples_sne2$j, var)

######create bins of coastline length#####
#write.csv(subsamples_sne2,"subsamples_sne_case1.csv")
library(dplyr)
subsamples_sne2<-subsamples_sne2%>%mutate(Coastlinebins =
cut(subsamples_sne2$env_mod, breaks =
c(0,200,400,600,800,1000,1200,1400,1600,1800,2000,2200)))
head(subsamples_sne2,200)
varbeta=tapply(subsamples_sne2$as.numeric.veg., subsamples_sne2$Coastlinebins,
var)
Coastlinebins=c("200","400","600","800","1000","1200","1400","1600","1800","2000","2200")
var=cbind(Coastline.bins=as.numeric(Coastlinebins),varbeta=as.numeric(round(varbeta,
4)))
variance_sne=as.data.frame(var)

####Individual bin method##############################
########################clubbing rows from a side only and keeping the
remaining rows#########################################################

env<-read.delim("Environment.txt", header=T)
row.names(env)=env[,1]# Declare column 1 as the row names#

s=array(0,dim = c(3000,225))
g=array(0,dim = c(3000,225))
betadata=array(0,dim = c(2,225))
meanbeta=array(0,dim = c(3000,225))
veg=array(0,dim = c(3000,225))
meanppd=vector(mode = "integer", length =1)
env_mod=vector(mode = "integer", length =10)
```

```
subsamps <- list()
subsamps<-list()

for(i in 1:50)
{
  env_mod=0
  subsamps1<-list()
  check=sample(1:14,2,replace = F)
  if(check[1]<check[2])
  {
    a=check[1]
    b=check[2]
    if(check[2]-check[1]==1)
    {
      betadata=ta.0[c(a,b),]
      betadata.pa=decostand(betadata,method = "pa")
      pair.s <- beta.pair(betadata.pa)
      b_sne <- as.matrix(pair.s$beta.sne)
      meanbeta=as.dist(b_sne)
      j=1
      x=(a+j)-1
      env_mod=env[x,9]
      out=data.frame(i,j,a,b,as.numeric(meanbeta),x,env_mod)
      subsamps1[[i]] <-out
    }else{
      for ( j in 1:((b-a)-1))
      {
        r=a+j
        t=a+j+1
        s=colSums(ta.0[c(a:r),])
        if(t==b)
        {g=ta.0[t,]}else{
          g=ta.0[c(t:b),]
        }
        betadata=rbind(s,g)
        betadata.pa=decostand(betadata,method = "pa")
        pair.s <- beta.pair(betadata.pa)
        b_sne <- as.matrix(pair.s$beta.sne)
        veg=as.dist(b_sne)
        meanbeta=mean(veg)
        x=(a+j)-1
        env_mod=env_mod+env[x,9]
        out=data.frame(i,j,a,b,as.numeric(meanbeta),x,env_mod)
        subsamps1[[j]] <-out
      }
    }
  }else{
    a=check[2]
    b=check[1]
    if(check[1]-check[2]==1)
    {
      betadata=ta.0[c(a,b),]
      betadata.pa=decostand(betadata,method = "pa")
      pair.s <- beta.pair(betadata.pa)
```

```
      b_sne <- as.matrix(pair.s$beta.sne)
      meanbeta=as.dist(b_sne)
      j=1
      x=(a+j)-1
      env_mod=env[x,9]
      oute=data.frame(i,j,a,b,as.numeric(meanbeta),x,env_mod)
      subsamps1[[j]] <-out
    }else{
      a=check[2]
      b=check[1]
      for ( j in 1:((b-a)-1))
      {
        r=a+j
        t=a+j+1
        s=colSums(ta.0[c(a:r),])
        if(t==b)
        {g=ta.0[t,]}else{
          g=ta.0[c(t:b),]
        }
        betadata=rbind(s,g)
        betadata.pa=decostand(betadata,method = "pa")
        pair.s <- beta.pair(betadata.pa)
        b_sne <- as.matrix(pair.s$beta.sne)
        veg=as.dist(b_sne)
        meanbeta=mean(veg)
        x=(a+j)-1
        env_mod=env_mod+env[x,9]
        out=data.frame(i,j,a,b,as.numeric(meanbeta),x,env_mod)
        subsamps1[[j]] <-out
      }
    }
  }
  subsamples1 <- do.call("rbind",subsamps1)
  subsamps[[i]]=subsamples1
  subsamples_sne3 <- do.call("rbind",subsamps)
}


varbeta_sne2=tapply(subsamples_sne3$as.numeric.meanbeta., subsamples_sne3$j,
var)

######create bins of coastline length#####
#write.csv(subsamples_sne3,"subsamples_sne_case2.csv")
library(dplyr)
subsamples_sne3<-subsamples_sne3%>%mutate(Coastlinebins =
cut(subsamples_sne3$env_mod, breaks =
c(0,200,400,600,800,1000,1200,1400,1600,1800,2000,2200)))
head(subsamples_sne3,200)
varbeta=tapply(subsamples_sne3$as.numeric.meanbeta.,
subsamples_sne3$Coastlinebins, var)
Coastlinebins=c("200","400","600","800","1000","1200","1400","1600","1800","2000","2200")
var=cbind(Coastline.bins=as.numeric(Coastlinebins),varbeta=as.numeric(round(varbeta,
4)))
variance2_sne=as.data.frame(var)
```

```
########Table 3.3###########################################
######Correlation values###############
#############################################################################
par(mfrow=c(5,2))
par(mar=c(2,5,1,1))
#par(pty="s")
#####with boxes####
######ppd case 1################
plot(subsamples_ppd2$j,subsamples_ppd2$as.numeric.veg., col="black",
ylim=c(0.1,1),ylab=expression (beta[ppd]),xlab="Number of boxes/ lat bins")
cor.test(subsamples_ppd2$j,subsamples_ppd2$as.numeric.veg.,method="spearman")
####ppd case 2##############
plot(subsamples_ppd3$j,subsamples_ppd3$as.numeric.meanbeta., col="black",
ylim=c(0.1,1),ylab=expression (beta[ppd]),xlab="Number of boxes/ lat bins")
cor.test(subsamples_ppd3$j,subsamples_ppd3$as.numeric.meanbeta.,method="spearman")

############whit case1
plot(subsamples_whit2$j,subsamples_whit2$as.numeric.veg., col="black",
ylim=c(0.1,1),ylab=expression (beta[whit]),xlab="Number of boxes/ lat bins")
cor.test(subsamples_whit2$as.numeric.veg.,subsamples_whit2$j,method="spearman")
#######whit case 2
plot(subsamples_whit3$j,subsamples_whit3$as.numeric.meanbeta., col="black",
ylim=c(0.1,1),ylab=expression (beta[whit]),xlab="Number of boxes/ lat bins")
cor.test(subsamples_whit3$as.numeric.meanbeta.,subsamples_whit3$j,method="spearman")

###simpson case 1
plot(subsamples_simp2$j,subsamples_simp2$as.numeric.veg., col="black",
ylim=c(0,1),ylab=expression (beta[sim]),xlab="Number of boxes/ lat bins")
cor.test(subsamples_simp2$as.numeric.veg.,subsamples_simp2$j,method="spearman")
####simpson case 2
plot(subsamples_simp3$j,subsamples_simp3$as.numeric.meanbeta., col="black",
ylim=c(0,1),ylab=expression (beta[sim]),xlab="Number of boxes/ lat bins")
cor.test(subsamples_simp3$as.numeric.meanbeta.,subsamples_simp3$j,method="spearman")

###sorenson case 1
plot(subsamples_sor2$j,subsamples_sor2$as.numeric.veg., col="black",
ylim=c(0.1,1),ylab=expression (beta[sor]),xlab="Number of boxes/ lat bins")
cor.test(subsamples_sor2$as.numeric.veg.,subsamples_sor2$j,method="spearman")
#sorenson case 2
plot(subsamples_sor3$j,subsamples_sor3$as.numeric.meanbeta., col="black",
ylim=c(0.1,1),ylab=expression (beta[sor]),xlab="Number of boxes/ lat bins")
cor.test(subsamples_sor3$as.numeric.meanbeta.,subsamples_sor3$j,method="spearman")

#sne case 1
plot(subsamples_sne2$j,subsamples_sne2$as.numeric.veg., col="black",
ylim=c(0.1,1),ylab=expression (beta[sne]),xlab="Number of boxes/ lat bins")
cor.test(subsamples_sne2$j,subsamples_sne2$as.numeric.veg.,method="spearman")
##sne case 2
plot(subsamples_sne3$j,subsamples_sne3$as.numeric.meanbeta., col="black",
ylim=c(0.1,1),ylab=expression (beta[sne]),xlab="Number of boxes/ lat bins")
cor.test(subsamples_sne3$j,subsamples_sne3$as.numeric.meanbeta.,method="spearman")
```

```
#############with coastline length##########################################
##ppd case 1
plot(subsamples_ppd2$env_mod,subsamples_ppd2$as.numeric.veg., col="black",
ylim=c(0.5,1),ylab=expression (beta[ppd]),xlab="Coastline length")
cor.test(subsamples_ppd2$env_mod,subsamples_ppd2$as.numeric.veg.,method="spearman")
####ppd case 2
plot(subsamples_ppd3$env_mod,subsamples_ppd3$as.numeric.meanbeta.,
col="black", ylim=c(0.5,1),ylab=expression (beta[ppd]),xlab="Coastline
length")
cor.test(subsamples_ppd3$env_mod,subsamples_ppd3$as.numeric.meanbeta.,method="spearman")

######whit case 1
plot(subsamples_whit2$env_mod,subsamples_whit2$as.numeric.veg., col="black",
ylim=c(0.1,1),ylab=expression (beta[whit]),xlab="Coastline length")
cor.test(subsamples_whit2$env_mod,subsamples_whit2$as.numeric.veg.,method="spearman")
#######whit case 2
plot(subsamples_whit3$env_mod,subsamples_whit3$as.numeric.meanbeta.,
col="black", ylim=c(0.1,1),ylab=expression (beta[whit]),xlab="Coastline
length")
cor.test(subsamples_whit3$env_mod,subsamples_whit3$as.numeric.meanbeta.,method="spearman")

###simpson case 1
plot(subsamples_simp2$env_mod,subsamples_simp2$as.numeric.veg., col="black",
ylim=c(0,1),ylab=expression (beta[sim]),xlab="Coastline length")
cor.test(subsamples_simp2$env_mod,subsamples_simp2$as.numeric.veg.,method="spearman")
#####simpson case 2
plot(subsamples_simp3$env_mod,subsamples_simp3$as.numeric.meanbeta.,
col="black", ylim=c(0,1),ylab=expression (beta[sim]),xlab="Coastline length")
cor.test(subsamples_simp3$env_mod,subsamples_simp3$as.numeric.meanbeta.,method="spearman")

####sorenson case 1
plot(subsamples_sor2$env_mod,subsamples_sor2$as.numeric.veg., col="black",
ylim=c(0,1),ylab=expression (beta[sor]),xlab="Coastline length")
cor.test(subsamples_sor2$env_mod,subsamples_sor2$as.numeric.veg.,method="spearman")
###sorenson casse 2
plot(subsamples_sor3$env_mod,subsamples_sor3$as.numeric.meanbeta.,
col="black", ylim=c(0,1),ylab=expression (beta[sor]),xlab="Coastline length")
cor.test(subsamples_sor3$env_mod,subsamples_sor3$as.numeric.meanbeta.,method="spearman")

####sne case 1
plot(subsamples_sne2$env_mod,subsamples_sne2$as.numeric.veg., col="black",
ylim=c(0,1),ylab=expression (beta[sne]),xlab="Coastline length")
cor.test(subsamples_sne2$env_mod,subsamples_sne2$as.numeric.veg.,method="spearman")

##sne case 2
plot(subsamples_sne3$env_mod,subsamples_sne3$as.numeric.meanbeta.,
col="black", ylim=c(0,1),ylab=expression (beta[sne]),xlab="Coastline length")
cor.test(subsamples_sne3$env_mod,subsamples_sne3$as.numeric.meanbeta.,method="spearman")


######################variance################################################################
#ppd case 1
plot(1:length(varbeta_ppd),varbeta_ppd, pch=19, ylab=expression
(beta[ppd]~variance),xlab="Number of latitude bins/boxes")
```

```
cor.test(1:length(varbeta_ppd),varbeta_ppd,method="spearman")

plot(variance_ppd$Coastline.bins,variance_ppd$varbeta, pch=19, ylab=expression
(beta[ppd]~variance),xlab = "Coastline length")
cor.test(variance_ppd$Coastline.bins,variance_ppd$varbeta,method = "spearman")

###ppd case 2
plot( 1:length(varbeta_ppd3),varbeta_ppd3,pch=19, ylab=expression
(beta[ppd]~variance),xlab="Number of latitude bins/boxes")
cor.test(varbeta_ppd3,1:length(varbeta_ppd3),method = "spearman")

plot(variance2_ppd$Coastline.bins,variance2_ppd$varbeta,pch=19,
ylab=expression (beta[ppd]~variance),xlab = "Coastline length")
cor.test(variance2_ppd$Coastline.bins,variance2_ppd$varbeta,method =
"spearman")

#whit case 1##############
plot(1:length(varbeta_whit1),varbeta_whit1,pch=19, ylab=expression
(beta[whit]~variance),xlab="Number of latitude bins/boxes")
cor.test(1:length(varbeta_whit1),varbeta_whit1,method = "spearman")
#points(subsamples$j, subsamples$as.numeric.veg., ylim=c(0,9))

plot(variance_whit$Coastline.bins,variance_whit$varbeta,pch=19,
ylab=expression (beta[whit]~variance),xlab = "Coastline length")
cor.test(variance_whit$Coastline.bins,variance_whit$varbeta,method =
"spearman")

##whit case 2####################################
plot(1:length(varbeta_whit2),varbeta_whit2, pch=19, ylab=expression
(beta[whit]~variance),xlab="Number of latitude bins/boxes")
cor.test(1:length(varbeta_whit2),varbeta_whit2,method = "spearman")

plot(variance2_whit$Coastline.bins,variance2_whit$varbeta,pch=19,
ylab=expression (beta[whit]~variance),xlab = "Coastline length")
cor.test(variance2_whit$Coastline.bins,variance2_whit$varbeta,method =
"spearman")

######simp case 1###############
cor.test(1:length(varbeta_simp1),varbeta_simp1,method="spearman")
plot(1:length(varbeta_simp1),varbeta_simp1, pch=19, ylab=expression
(beta[simp]~variance),xlab="Number of latitude bins/boxes")

plot(variance_simp$Coastline.bins,variance_simp$varbeta,pch=19,
ylab=expression (beta[simp]~variance),xlab = "Coastline length")
cor.test(variance_simp$Coastline.bins,variance_simp$varbeta,method =
"spearman")

########simp case 2
plot(1:length(varbeta_simp2),varbeta_simp2, pch=19, ylab=expression
(beta[simp]~variance),xlab="Number of latitude bins/boxes")
cor.test(1:length(varbeta_simp2),varbeta_simp2,method = "spearman")

plot(variance2$Coastline.bins,variance2$varbeta,pch=19, ylab=expression
(beta[simp]~variance),xlab = "Coastline length")
```

```
cor.test(variance2$Coastline.bins,variance2$varbeta,method = "spearman")

#######sorenson case 1
cor.test(1:length(varbeta_sor),varbeta_sor,method="spearman")
plot(1:length(varbeta_sor),varbeta_sor, pch=19, ylab=expression
(beta[sor]~variance),xlab="Number of latitude bins/boxes")

plot(variance_sor$Coastline.bins,variance_sor$varbeta,pch=19,ylab=expression
(beta[sor]~variance),xlab = "Coastline length")
cor.test(variance_sor$Coastline.bins,variance_sor$varbeta,method = "spearman")

###sorenson case 2
plot(1:length(varbeta_sor2),varbeta_sor2,pch=19,ylab=expression
(beta[sor]~variance),xlab="Number of latitude bins/boxes")
cor.test(varbeta_sor2,c(1:length(varbeta_sor2)),method = "spearman")

plot(variance2_sor$Coastline.bins,variance2_sor$varbeta,pch=19,
ylab=expression (beta[sor]~variance),xlab = "Coastline length")
cor.test(variance2_sor$Coastline.bins,variance2_sor$varbeta,method =
"spearman")

###sne case 1
cor.test(1:length(varbeta_sne),varbeta_sne,method="spearman")
plot(1:length(varbeta_sne),varbeta_sne, pch=19,ylab=expression
(beta[sne]~variance),xlab="Number of latitude bins/boxes")

plot(variance_sne$Coastline.bins,variance_sne$varbeta,pch=19, ylab=expression
(beta[sne]~variance),xlab = "Coastline length")
cor.test(variance_sne$Coastline.bins,variance_sne$varbeta,method = "spearman")

###sne case 2
plot(1:length(varbeta_sne2),varbeta_sne2,pch=19, ylab=expression
(beta[sne]~variance),xlab="Number of latitude bins/boxes")
cor.test(varbeta_sne2,1:length(varbeta_sne2),method = "spearman")

plot(variance2_sne$Coastline.bins,variance2_sne$varbeta,pch=19,ylab=expression
(beta[sne]~variance),xlab = "Coastline length")
cor.test(variance2_sne$Coastline.bins,variance2_sne$varbeta,method =
"spearman")



#########Figure 3.4###########################
#############with coastline length###########################################
##ppd case 1
par(mfrow=c(5,2))
par(mar=c(2,5,1,3))
plot(subsamples_ppd2$env_mod,subsamples_ppd2$as.numeric.veg.,ylim=c(0.1,1),
xlim=c(0,2500), col="black", ylab=expression (beta[ppd]),xlab="Coastline
length",las=1,cex.lab=1.5)
par(new=TRUE)
plot(variance_ppd$Coastline.bins,variance_ppd$varbeta,
axes=F,type="b",pch=19,col="red",xlab=" ",ylab =" ")
axis(4,ylim=c(0,0.1),las=1)
```

```
####ppd case 2
par(mar=c(2,3,1,6))
plot(subsamples_ppd3$env_mod,subsamples_ppd3$as.numeric.meanbeta.,
xlim=c(0,2500),col="black", ylim=c(0.1,1),ylab=" ",xlab="Coastline
length",las=1)
par(new=TRUE)
plot(variance2_ppd$Coastline.bins,variance2_ppd$varbeta,
axes=F,type="b",pch=19,col="red",xlab=" ",ylab =" ")
axis(4,ylim=c(0,0.1),las=1)
mtext(expression (variance~beta[ppd]),side=4,cex = 1,line=4)

######whit case 1
par(mar=c(2,5,1,3))
plot(subsamples_whit2$env_mod,subsamples_whit2$as.numeric.veg.,xlim=c(0,2500),
col="black", ylim=c(0.1,1),ylab=expression (beta[whit]),xlab="Coastline
length",las=1,cex.lab=1.5)
par(new=TRUE)
plot(variance_whit$Coastline.bins,variance_whit$varbeta,
axes=F,type="b",pch=19,col="red",xlab=" ",ylab =" ")
axis(4,ylim=c(0,0.1),las=1)
#######whit case 2
par(mar=c(2,3,1,6))
plot(subsamples_whit3$env_mod,subsamples_whit3$as.numeric.meanbeta.,xlim=c(0,2500),
col="black", ylim=c(0.1,1),ylab=" ",xlab="Coastline length",las=1)
par(new=TRUE)
plot(variance2_whit$Coastline.bins,variance2_whit$varbeta,
axes=F,type="b",pch=19,col="red",xlab=" ",ylab =" ")
axis(4,ylim=c(0,0.1),las=1)
mtext(expression (variance~beta[whit]),side=4,col="black",cex = 1,line=4)

###simpson case 1
par(mar=c(2,5,1,3))
plot(subsamples_simp2$env_mod,subsamples_simp2$as.numeric.veg.,xlim=c(0,2500),
col="black", ylim=c(0.1,1),ylab=expression (beta[sim]),xlab="Coastline
length",las=1,cex.lab=1.5)
par(new=TRUE)
plot(variance_simp$Coastline.bins,variance_simp$varbeta,
axes=F,type="b",pch=19,col="red",xlab=" ",ylab =" ")
axis(4,ylim=c(0,0.1),las=1)
####simpson case 2
par(mar=c(2,3,1,6))
plot(subsamples_simp3$env_mod,subsamples_simp3$as.numeric.meanbeta.,xlim=c(0,2500),
col="black", ylim=c(0.1,1),ylab=" ",xlab="Coastline length",las=1)
par(new=TRUE)
plot(variance2$Coastline.bins,variance2$varbeta,
axes=F,type="b",pch=19,col="red",xlab=" ",ylab =" ")
axis(4,ylim=c(0,0.1),las=1)
mtext(expression (variance~beta[sim]),side=4,col="black",cex = 1,line=4)

###sorenson case 1
par(mar=c(2,5,1,3))
```

```
plot(subsamples_sor2$env_mod,subsamples_sor2$as.numeric.veg.,xlim=c(0,2500),
col="black", ylab=expression (beta[sor]),xlab="Coastline
length",las=1,cex.lab=1.5)
par(new=TRUE)
plot(variance_sor$Coastline.bins,variance_sor$varbeta,
axes=F,type="b",pch=19,col="red",xlab=" ",ylab =" ")
axis(4,ylim=c(0,0.1),las=1)
###sorenson casse 2
par(mar=c(2,3,1,6))
plot(subsamples_sor3$env_mod,subsamples_sor3$as.numeric.meanbeta.,xlim=c(0,2500),
col="black", ylim=c(0.1,1),ylab=" ",xlab="Coastline length",las=1)
par(new=TRUE)
plot(variance2_sor$Coastline.bins,variance2_sor$varbeta,
axes=F,type="b",pch=19,col="red",xlab=" ",ylab =" ")
axis(4,ylim=c(0,0.1),las=1)
mtext(expression (variance~beta[sor]),side=4,col="black",cex = 1,line=4)

####sne case 1
par(mar=c(2,5,1,3))
plot(subsamples_sne2$env_mod,subsamples_sne2$as.numeric.veg.,
xlim=c(0,2500),col="black", ylim=c(0.1,1),ylab=expression
(beta[sne]),xlab="Coastline length",las=1,cex.lab=1.5)
par(new=TRUE)
plot(variance_sne$Coastline.bins,variance_sne$varbeta,
axes=F,type="b",pch=19,col="red",xlab=" ",ylab =" ")
axis(4,ylim=c(0,0.1),las=1)

##sne case 2
par(mar=c(2,3,1,6))
plot(subsamples_sne3$env_mod,subsamples_sne3$as.numeric.meanbeta.,
xlim=c(0,2500),col="black", ylim=c(0.1,1),ylab=" ",xlab="Coastline
length",las=1)
par(new=TRUE)
plot(variance2_sne$Coastline.bins,variance2_sne$varbeta,
axes=F,type="b",pch=19,col="red",xlab=" ",ylab =" ")
axis(4,ylim=c(0,0.1),las=1)
mtext(expression (variance~beta[sne]),side=4,col="black",cex = 1,line=4)

#########Figure 3.S2##############################################
###############plots##########################
#####with boxes####
par(mfrow=c(5,2))
par(mar=c(2,5,1,3))
######ppd case 1################
plot(subsamples_ppd2$j,subsamples_ppd2$as.numeric.veg., col="black",
ylim=c(0.1,1),ylab=expression (beta[ppd]),xlab="Number of boxes/ lat
bins",las=1,cex.lab=1.5)
par(new=TRUE)
plot(1:length(varbeta_ppd),varbeta_ppd,
axes=F,type="b",pch=19,col="red",xlab=" ",ylab =" ")
axis(4,ylim=c(0,0.05),las=1)

####ppd case 2###############
par(mar=c(2,3,1,6))
```

```
plot(subsamples_ppd3$j,subsamples_ppd3$as.numeric.meanbeta., col="black",
ylim=c(0.1,1),ylab=" ",xlab="Number of boxes/ lat bins",las=1)
par(new=TRUE)
plot(1:length(varbeta_ppd3),varbeta_ppd3,
axes=F,type="b",pch=19,col="red",xlab=" ",ylab =" ")
axis(4,ylim=c(0,0.1),las=1)
mtext(expression (variance~beta[ppd]),side=4,cex = 1,line=4)

############whit case1
par(mar=c(2,5,1,3))
plot(subsamples_whit2$j,subsamples_whit2$as.numeric.veg., col="black",
ylim=c(0.1,1),ylab=expression (beta[whit]),xlab="Number of boxes/ lat
bins",las=1,cex.lab=1.5)
par(new=TRUE)
plot(1:length(varbeta_whit1),varbeta_whit1,
axes=F,type="b",pch=19,col="red",xlab=" ",ylab =" ")
axis(4,ylim=c(0,0.05),las=1)
#######whit case 2
par(mar=c(2,3,1,6))
plot(subsamples_whit3$j,subsamples_whit3$as.numeric.meanbeta., col="black",
ylim=c(0.1,1),ylab=" ",xlab="Number of boxes/ lat bins",las=1)
par(new=TRUE)
plot(1:length(varbeta_whit2),varbeta_whit2,
axes=F,type="b",pch=19,col="red",xlab=" ",ylab =" ")
axis(4,ylim=c(0,0.1),las=1)
mtext(expression (variance~beta[whit]),side=4,col="black",cex = 1,line=4)

###simpson case 1
par(mar=c(2,5,1,3))
plot(subsamples_simp2$j,subsamples_simp2$as.numeric.veg., col="black",
ylim=c(0.1,1),ylab=expression (beta[sim]),xlab="Number of boxes/ lat
bins",las=1,cex.lab=1.5)
par(new=TRUE)
plot(1:length(varbeta_simp1),varbeta_simp1,
axes=F,type="b",pch=19,col="red",xlab=" ",ylab =" ")
axis(4,ylim=c(0,0.05),las=1)
####simpson case 2
par(mar=c(2,3,1,6))
plot(subsamples_simp3$j,subsamples_simp3$as.numeric.meanbeta., col="black",
ylim=c(0.1,1),ylab=" ",xlab="Number of boxes/ lat bins",las=1)
par(new=TRUE)
plot(1:length(varbeta_simp2),varbeta_simp2,
axes=F,type="b",pch=19,col="red",xlab=" ",ylab =" ")
axis(4,ylim=c(0,0.1),las=1)
mtext(expression (variance~beta[sim]),side=4,col="black",cex = 1,line=4)

###sorenson case 1
par(mar=c(2,5,1,3))
plot(subsamples_sor2$j,subsamples_sor2$as.numeric.veg., col="black",
ylim=c(0.1,1),ylab=expression (beta[sor]),xlab="Number of boxes/ lat
bins",las=1,cex.lab=1.5)
par(new=TRUE)
plot(1:length(varbeta_sor),varbeta_sor,
axes=F,type="b",pch=19,col="red",xlab=" ",ylab =" ")
```

```
axis(4,ylim=c(0,0.05),las=1)
#sorenson case 2
par(mar=c(2,3,1,6))
plot(subsamples_sor3$j,subsamples_sor3$as.numeric.meanbeta., col="black",
ylim=c(0.1,1),ylab=" ",xlab="Number of boxes/ lat bins",las=1)
par(new=TRUE)
plot(1:length(varbeta_sor2),varbeta_sor2,
axes=F,type="b",pch=19,col="red",xlab=" ",ylab =" ")
axis(4,ylim=c(0,0.1),las=1)
mtext(expression (variance~beta[sor]),side=4,col="black",cex = 1,line=4)

#sne case 1
par(mar=c(2,5,1,3))
plot(subsamples_sne2$j,subsamples_sne2$as.numeric.veg., col="black",
ylim=c(0.1,1),ylab=expression (beta[sne]),xlab="Number of
bins",las=1,cex.lab=1.5)
par(new=TRUE)
plot(1:length(varbeta_sne),varbeta_sne,
axes=F,type="b",pch=19,col="red",xlab=" ",ylab =" ")
axis(4,ylim=c(0,0.05),las=1)
##sne case 2
par(mar=c(2,3,1,6))
plot(subsamples_sne3$j,subsamples_sne3$as.numeric.meanbeta., col="black",
ylim=c(0.1,1),ylab=" ",xlab="Number of bins",las=1)
par(new=TRUE)
plot(1:length(varbeta_sne2),varbeta_sne2,
axes=F,type="b",pch=19,col="red",xlab=" ",ylab =" ")
axis(4,ylim=c(0,0.1),las=1)
mtext(expression (variance~beta[sne]),side=4,col="black",cex = 1,line=4)

#####Figure 3.5 DA##########################################
###$observed beta values vs coastlength#######
env<-read.delim("Environment.txt", header=T) ##containing details of regional
environmental parameter##
row.names(env)=env[,1]
env[is.na(env)]

env$cumulative= ave(env$Coastline.length..Km.,FUN=cumsum) ###calculating
cumulative coastlength

par(pty="s")
par(mar=c(2,5,1,1))
par(mfrow=c(5,1))
plot(env$cumulative,locs,pch=19,col="black",xlab="Coastline
length",ylab=expression (beta[Obs_ppd]))
cor.test(env$cumulative,locs,method="spearman")

plot(env$cumulative,locs1,pch=19,col="black",xlab="Coastline
length",ylab=expression (beta[Obs_whit]))
cor.test(env$cumulative,locs1,method="spearman")

plot(env$cumulative,locs_sim,pch=19,col="black",xlab="Coastline
length",ylab=expression (beta[Obs_sim]))
cor.test(env$cumulative,locs_sim,method="spearman")
```

```
plot(env$cumulative,locs_sor,pch=19,col="black",xlab="Coastline
length",ylab=expression (beta[Obs_sor]))
cor.test(env$cumulative,locs_sor,method="spearman")

plot(env$cumulative,locs_sne,pch=19,col="black",xlab="Coastline
length",ylab=expression (beta[Obs_sne]))
cor.test(env$cumulative,locs_sne,method="spearman")


############Figure 3.6 DA##################################
#####K-s test between null model values and actual WC beta values#####
#####ppd case 1###########
Dist_tidal2=array(0,c(10000,1))
pvalue_tidal2=array(0,c(10000,1))
for(i in 1:10000)
{
  D5=sample(t(subsamples_ppd2$as.numeric.veg.),14,replace = T)
  D6=sample(t(locs),14,replace=T)
  Dist_tidal2[i]=ks.test(as.matrix(D6),as.matrix(D5))$statistic
  pvalue_tidal2[i]=ks.test(as.matrix(D6),as.matrix(D5))$p.value
}
mean(pvalue_tidal2)
mean(Dist_tidal2)
median(pvalue_tidal2)

#####ppd case 2##########
Dist_rest2=array(0,c(10000,1))
pvalue_rest2=array(0,c(10000,1))
for(i in 1:10000)
{
  D7=sample(t(subsamples_ppd3$as.numeric.meanbeta.),14,replace = T)
  D8=sample(t(locs),14,replace=T)
  Dist_rest2[i]=ks.test(as.matrix(D8),as.matrix(D7))$statistic
  pvalue_rest2[i]=ks.test(as.matrix(D8),as.matrix(D7))$p.value
}
mean(pvalue_rest2)
mean(Dist_rest2)
median(pvalue_rest2)
###########################
######whit case 1
Dist_whit1=array(0,c(10000,1))
pvalue_whit1=array(0,c(10000,1))
for(i in 1:10000)
{
  D1=sample(t(subsamples_whit2$as.numeric.veg.),14,replace = T)
  Dist_whit1[i]=ks.test(t(locs1),as.matrix(D1))$statistic
  pvalue_whit1[i]=ks.test(t(locs1),as.matrix(D1))$p.value
}
median(pvalue_whit1)
hist(pvalue_whit1)
hist(Dist_whit1)
```

```
#######whit case 2
Dist_whit2=array(0,c(10000,1))
pvalue_whit2=array(0,c(10000,1))
for(i in 1:10000)
{
  D1=sample(t(subsamples_whit3$as.numeric.meanbeta.),14,replace = T)
  Dist_whit2[i]=ks.test(t(locs1),as.matrix(D1))$statistic
  pvalue_whit2[i]=ks.test(t(locs1),as.matrix(D1))$p.value
}
median(pvalue_whit2)
hist(pvalue_whit2)
hist(Dist_whit2)


###simpson case 1
Dist_simp1=array(0,c(10000,1))
pvalue_simp1=array(0,c(10000,1))
for(i in 1:10000)
{
  D1=sample(t(subsamples_simp2$as.numeric.veg.),14,replace = T)
  Dist_simp1[i]=ks.test(t(locs_sim),as.matrix(D1))$statistic
  pvalue_simp1[i]=ks.test(t(locs_sim),as.matrix(D1))$p.value
}
median(pvalue_simp1)
hist(pvalue_simp1)
hist(Dist_simp1)


#####simpson case 2
Dist_simp2=array(0,c(10000,1))
pvalue_simp2=array(0,c(10000,1))
for(i in 1:10000)
{
  D1=sample(t(subsamples_simp3$as.numeric.meanbeta.),14,replace = T)
  Dist_simp2[i]=ks.test(t(locs_sim),as.matrix(D1))$statistic
  pvalue_simp2[i]=ks.test(t(locs_sim),as.matrix(D1))$p.value
}
median(pvalue_simp2)
hist(pvalue_simp2)
hist(Dist_simp2)


####sorenson case 1
Dist_sor1=array(0,c(10000,1))
pvalue_sor1=array(0,c(10000,1))
for(i in 1:10000)
{
  D1=sample(t(subsamples_sor2$as.numeric.veg.),14,replace = T)
  Dist_sor1[i]=ks.test(t(locs_sor),as.matrix(D1))$statistic
  pvalue_sor1[i]=ks.test(t(locs_sor),as.matrix(D1))$p.value
}
median(pvalue_sor1)
hist(pvalue_sor1)
hist(Dist_sor1)
```

```
###sorenson casse 2
Dist_sor2=array(0,c(10000,1))
pvalue_sor2=array(0,c(10000,1))
for(i in 1:10000)
{
  D1=sample(t(subsamples_sor3$as.numeric.meanbeta.),14,replace = T)
  Dist_sor2[i]=ks.test(t(locs_sor),as.matrix(D1))$statistic
  pvalue_sor2[i]=ks.test(t(locs_sor),as.matrix(D1))$p.value
}
median(pvalue_sor2)
hist(pvalue_sor2)
hist(Dist_sor2)

####sne case 1
Dist_sne1=array(0,c(10000,1))
pvalue_sne1=array(0,c(10000,1))
for(i in 1:10000)
{
  D1=sample(t(subsamples_sne2$as.numeric.veg.),14,replace = T)
  Dist_sne1[i]=ks.test(t(locs_sne),as.matrix(D1))$statistic
  pvalue_sne1[i]=ks.test(t(locs_sne),as.matrix(D1))$p.value
}
median(pvalue_sne1)
median(Dist_sne1)
hist(pvalue_sne1)
hist(Dist_sne1)


##sne case 2
Dist_sne2=array(0,c(10000,1))
pvalue_sne2=array(0,c(10000,1))
for(i in 1:10000)
{
  D1=sample(t(subsamples_sne3$as.numeric.meanbeta.),14,replace = T)
  Dist_sne2[i]=ks.test(t(locs_sne),as.matrix(D1))$statistic
  pvalue_sne2[i]=ks.test(t(locs_sne),as.matrix(D1))$p.value
}
median(pvalue_sne2)
median(Dist_sne2)
hist(pvalue_sne2)
hist(Dist_sne2)


################histograms####################################################
par(mar=c(2,5,1,1))
par(mfrow=c(5,2))
p_ppd=hist(Dist_tidal2)
p_ppd2=hist(Dist_rest2)
p_whit=hist(Dist_whit1)
p_whit2=hist(Dist_whit2)
p_simp=hist(Dist_simp1)
p_simp2=hist(Dist_simp2)
```

```
p_sor=hist(Dist_sor1)
p_sor2=hist(Dist_sor2)
p_sne=hist(Dist_sne1)
p_sne2=hist(Dist_sne2)


plot(p_ppd,w=10,col=c("deeppink"),xlim = c(0.2,1),ylim =
c(0,5000),cex.axis=1.5,ann=FALSE)
plot(p_ppd2,w=10,col=c("deeppink"),xlim = c(0.2,1),ylim =
c(0,5000),cex.axis=1.5,ann=FALSE)


plot(p_whit,col=c("dodgerblue2"),xlim = c(0.2,1),ylim =
c(0,5000),cex.axis=1.5,ann=FALSE)
plot(p_whit2,col=c("dodgerblue2"),xlim = c(0.2,1),ylim =
c(0,5000),cex.axis=1.5,ann=FALSE)



plot(p_simp,col=c("darkorange2"),xlim = c(0.2,1),ylim =
c(0,5000),cex.axis=1.5,ann=FALSE)
plot(p_simp2,col=c("darkorange2"),xlim = c(0.2,1),ylim =
c(0,5000),cex.axis=1.5,ann=FALSE)


plot(p_sor,col=c("green4"),xlim = c(0.2,1),ylim =
c(0,5000),cex.axis=1.5,ann=FALSE)
plot(p_sor2,col=c("green4"),xlim = c(0.2,1),ylim =
c(0,5000),cex.axis=1.5,ann=FALSE)


plot(p_sne,col=c("yellow"),xlim = c(0.2,1),ylim =
c(0,5000),cex.axis=1.5,ann=FALSE)
plot(p_sne2,col=c("yellow"),xlim = c(0.2,1),ylim =
c(0,5000),cex.axis=1.5,ann=FALSE)



###############beta div vs
environment################################################################################
library(ade4)
library(vegan)
library(MASS)
library(ellipse)
library(FactoMineR)


env=read.csv("Environment - Copy.csv", header=T)##containing details of
regional environmental parameter in correct order##
row.names(env)=env[,1]
env[is.na(env)]=0

env.df=data.frame(env)
prodmean=env.df$Productivity..mgC.m.2.day..
```

```
prodrange=env.df$Productivity.range..mgC.m.2.day.
salmean=env.df$Salinity..unit.less.
salrange=as.numeric(env.df$Salinity.range)
tempmean=env.df$Temp.mean..degree.C.
temprange=env.df$Temperature..deg.C..
#coastlength=env.df$Coastline.length..Km.
#river=env.df$Rivers
#shelfwidth=env.df$Shelf.width.m.
#gradient=env.df$Gradient.degree.
cyclonefrq=env.df$Cyclones
oxygen=env.df$Oxygen.ppm.
shelfarea=env.df$Shelf.area.km2.


###########################################################
require(Hmisc)
env.df=env.df[,-c(1,8,10,11,12,13,15)]
corr=rcorr(as.matrix(env.new),type="spearman") #####for computing correlation
with significances#
pvalues=corr[["P"]]
rhos=corr[["r"]]
write.table(rhos,file="correlations.csv")
write.table(pvalues,file="pvalues.csv")

####with shelf area####################
require(Hmisc)
env.df=env.df[,-c(1,9)]
corr=rcorr(as.matrix(env.df),type="spearman") #####for computing correlation
with significances#
pvalues=corr[["P"]]
rhos=corr[["r"]]
write.table(rhos,file="correlations_new.csv")
write.table(pvalues,file="pvalues_spt30.csv")

#######Figure 3.7#############################################
###############################################################################
ppd1=locs[1:14]



##########correlation of beta diversity with environmental
variables################

par(mfrow=c(4,2))
par(mar=c(4,4,1,1))

plot(prodmean[1:14],ppd1,xlab="Productivity (mean)", ylab=expression
(beta[ppd]), pch=1,ylim=c(0,1),cex.lab=1)
cor.test(prodmean,ppd1,method = "spearman")

plot(prodrange[1:14],ppd1,xlab="Productivity (range)", ylab=expression
(beta[ppd]), ylim=c(0,1),pch=1,cex.lab=1)
cor.test(prodrange,ppd1,method = "spearman")
```

```
plot(salmean[1:14],ppd1, xlab="Salinity (mean)", ylab=expression
(beta[ppd]),ylim=c(0,1),cex.lab=1)
cor.test(salmean,ppd1,method = "spearman")

plot(salrange[1:14],ppd1, xlab="Salinity (range)", ylab=expression
(beta[ppd]),ylim=c(0,1),cex.lab=1)
cor.test(salrange,ppd1,method = "spearman")

#plot(tempmean[1:14],ppd, xlab="Temperature (mean)", ylab="Mean
PPD",ylim=c(0,5))
#cor.test(tempmean,ppd,method = "spearman")

plot(temprange[1:14],ppd1, xlab="Temperature (range)", ylab=expression
(beta[ppd]),ylim=c(0,1),cex.lab=1)
cor.test(temprange,ppd1,method = "spearman")

#plot(shelfwidth[1:14],ppd, xlab="Coastline length", ylab="Mean
PPD",ylim=c(0,5))
#cor.test(shelfwidth,ppd,method = "spearman")

#plot(gradient[1:14],ppd, xlab="Rivers", ylab="Mean PPD",ylim=c(0,5))
#cor.test(gradient,ppd,method = "spearman")

plot(shelfarea[1:14],ppd1, xlab="Shelf area", ylab=expression
(beta[ppd]),ylim=c(0,1),cex.lab=1)
cor.test(shelfarea,ppd1,method = "spearman")

plot(oxygen[1:14],ppd1, xlab="Oxygen", ylab=expression
(beta[ppd]),ylim=c(0,1),cex.lab=1)
cor.test(oxygen,ppd1,method = "spearman")

plot(cyclonefrq[1:14],ppd1, xlab="Cyclones", ylab=expression
(beta[ppd]),ylim=c(0,1),cex.lab=1)
cor.test(cyclonefrq,ppd1,method = "spearman")

##########Table 3.4####################################
####multiple glm########
out=summary(glm(ppd1~prodmean+prodrange+salmean+tempmean+temprange+oxygen+cyclonefrq+shelf

#out=summary(glm(ppd~prodmean+prodrange+salmean+salrange+tempmean+temprange+Oxygen+cyclone
summary(out)
finalglm=rbind(out$coefficients)
#write.table(finalglm, file="GLM output.txt")


#############single glm##############################
## GLM overall_Single 566
model2=glm(ppd~prodmean)
tst2=summary(model2)
model12=glm(ppd~prodrange)
tst12=summary(model12)
model3=glm(ppd~salmean)
tst3=summary(model3)
```

```
model4=glm(ppd~tempmean)
tst4=summary(model4)

model14=glm(ppd~temprange)
tst14=summary(model14)

model5=glm(ppd~oxygen)
tst5=summary(model5)

model6=glm(ppd~cyclonefrq)
tst6=summary(model6)

model16=glm(ppd~shelfarea)
tst16=summary(model16)

#Figure 3.8
DA###########################################################################
#####performing RDA with selective variables##############
#########RDA analysis###################
par(mfrow=c(1,1))
par(mar=c(4,2,2,2))
#spe.rda=rda(ta.01~prodmean+salmean+temprange+cyclonefrq+shelfarea+oxygen,
data=env.df)
spe.rda=rda(ta.01 ~ prodrange + salrange + salmean + tempmean+cyclonefrq,data
= env.df)
anova(spe.rda)
summary(spe.rda)####make sure to run ade4 before vegan apckage for this to
work###########
coef(spe.rda)
adjR2.tbrda <- RsquareAdj (spe.rda)$adj.r.squared
env.new=env.df[,-c(1,2,4,8,9,11)]
sel.fs <- forward.sel (Y = ta.01, X = env.new, adjR2thresh = adjR2.tbrda)
tb_rda.vasc.0 <- rda (ta.01 ~ 1, data = env.df)
sel.osR2 <- ordiR2step (tb_rda.vasc.0 , scope = formula (spe.rda), R2scope =
adjR2.tbrda, direction = 'forward', permutations = 9999)


plot(spe.rda,scaling=2,
col="black",display=c("cn","lc"),xlim=c(-80,80),ylim=c(-30,30))
plot(spe.rda,scaling=2,col="black",xlim=c(-80,80),ylim=c(-30,30))

plot(spe.rda,type="n",xlim=c(-80,80),ylim=c(-30,30))
points(spe.rda,display="lc",labels=rownames(ppd),col="black")
text(spe.rda,display = "cn",col="gray41")
(R2adj=RsquareAdj(spe.rda)$adj.r.squared)
spe.rda.all <- rda(ta.01~ ., data=env)
(R2a.all <- RsquareAdj(spe.rda.all)$adj.r.squared)


ta.prop=ta.prop[-c(15,16),]
require(adespatial)
#env=env[,-c(2,6,10)]
env=env[,-1]
```

```
row.names(env)=row.names(ta.01)
forward.sel(ta.01,env,adjR2thresh=R2adj)
#################################################################
## 6. CANONICAL CORRESPONDENCE ANALYSIS using (cca{vegan})

CCAres<- cca(ta.01 ~ prodrange + salrange + salmean + tempmean+cyclonefrq)
plot(CCAres, display=c("sites", "cn"),col="green")
summary(CCAres)
scores(CCAres)
CCAres$CCA$centroids
CCAres$CCA$v
CCAres$CCA$u

#CCAres<- cca(ta.01 ~ prodrange + salrange + salmean + tempmean+shelfarea+
oxygen+cyclonefrq)
#plot(CCAres, display=c("sites", "cn"),col="green")

CCAres<- cca(ta.01 ~ prodrange + prodmean+ salmean + tempmean+shelfarea+
oxygen+cyclonefrq)
plot(CCAres, display=c("sites", "cn"),col="green")


#######LA beta diversity################################

Sys.setlocale("LC_ALL", "C")
#masterbothcoast=read.delim("Occurence_Master (1).txt", header=T)
masterbothcoast=read.csv("Occurence_Master (1).csv", header=T)
masterlive=masterbothcoast[masterbothcoast$Coast=="West",]
count=c(rep(1,509))
masterlive=cbind(masterlive,count)
ta.live <- tapply(masterlive$count,
list(masterlive$Latitudinal.bin,masterlive$Name), sum)
ta.0 <- ta.live
ta.0[is.na(ta.0)] <- 0
#ta.0=ta.0[,-1]
rownames(ta.
0)=c("22","21","20","19","18","17","16","15","14","13","12","11","10","9","8")

#####change the order of the rows from 8-21##############
require(dplyr)
ta.01=arrange(as.data.frame(ta.0), -row_number())
rownames(ta.
01)=c("8","9","10","11","12","13","14","15","16","17","18","19","20","21","22")

ta.01=ta.01[-15,] ###remove latitude bin 22###

###change it to presence-absence matrix
ta.pa=decostand(ta.01,method = "pa")

##run the environment data
env<-read.delim("Environment.txt", header=T)
rownames(env)=c("21","20","19","18","17","16","15","14","13","12","11","10","9","8")
env[,1]=rownames(env)
```

```
env$cumulative= ave(env$Coastline.length..Km.,FUN=cumsum) ###calculating
cumulative coastlength

####calculating beta diversity using bray -curtis ######
#####PPD bray curtis#######
ta.0 <- as.matrix(ta.pa)
ta.prop <-prop.table(ta.0, margin=1) #change to proportions
ppd <- vegdist(ta.prop, method="bray") #distance matrix
mean(ppd) #mean beta diversity
dppd <- as.matrix(ppd)
dppd[dppd==0] <- NA #replace zeros with NA to calculate means and standard
error
locs <- apply(dppd, 2, FUN=mean, na.rm=T) #mean beta diversity per site with
regard to other sites #Tables 1,2

#####Null model in bray curtis########
###Combined bin method case 1#########
env<-read.delim("Environment.txt", header=T)
rownames(env)=c("21","20","19","18","17","16","15","14","13","12","11","10","9","8")

s=array(0,dim = c(3000,225))
g=array(0,dim = c(3000,225))
betadata=array(0,dim = c(2,225))
meanbeta=array(0,dim = c(3000,225))
veg=array(0,dim = c(3000,225))
meanppd=vector(mode = "integer", length =1)
env_mod=vector(mode = "integer", length =10)
subsamps <- list()
subsamps1<-list()

for(i in 1:50)
{
  env_mod=0
  subsamps1<-list()
  check=sample(1:14,2,replace = F)
  if(check[1]<check[2])
  {
    a=check[1]
    b=check[2]
    if(check[2]-check[1]==1)
    {
      betadata=ta.0[c(a,b),]
      ta.prop <-prop.table(betadata, margin=1)
      veg <- vegdist(ta.prop, method="bray")
      j=1
      x=(a+j)-1
      env_mod=env[x,9]
      out=data.frame(i,j,a,b,as.numeric(veg),x,env_mod)
      subsamps1[[i]] <-out
    }else{
      for ( j in 1:((b-a)-1))
      {
        r=a+j
        t=a+j+1
```

```
        s=colSums(ta.0[c(a:r),])
        if(t==b)
        {g=ta.0[t,]}else{
          g=colSums(ta.0[c(t:b),])
        }
        betadata=rbind(s,g)
        ta.prop <-prop.table(betadata, margin=1)
        veg <- vegdist(ta.prop, method="bray")
        x=(a+j)-1
        env_mod=env_mod+env[x,9]
        out=data.frame(i,j,a,b,as.numeric(veg),x,env_mod)
        subsamps1[[j]] <-out
      }
    }
  }else{
    a=check[2]
    b=check[1]
    if(check[1]-check[2]==1)
    {
      betadata=ta.0[c(a,b),]
      ta.prop <-prop.table(betadata, margin=1)
      veg <- vegdist(ta.prop, method="bray")
      j=1
      x=(a+j)-1
      env_mod=env[x,9]
      oute=data.frame(i,j,a,b,as.numeric(veg),x,env_mod)
      subsamps1[[j]] <-out
    }else{
      a=check[2]
      b=check[1]
      for ( j in 1:((b-a)-1))
      {
        r=a+j
        t=a+j+1
        s=colSums(ta.0[c(a:r),])
        if(t==b)
        {g=ta.0[t,]}else{
          g=colSums(ta.0[c(t:b),])
        }
        betadata=rbind(s,g)
        ta.prop <-prop.table(betadata, margin=1)
        veg <- vegdist(ta.prop, method="bray")
        x=(a+j)-1
        env_mod=env_mod+env[x,9]
        out=data.frame(i,j,a,b,as.numeric(veg),x,env_mod)
        subsamps1[[j]] <-out
      }
    }
  }
  subsamples1 <- do.call("rbind",subsamps1)
  subsamps[[i]]=subsamples1
  subsamples_ppd2<- do.call("rbind",subsamps)
}
```

```
####calculating variance for bins########
varbeta_ppd=tapply(subsamples_ppd2$as.numeric.veg., subsamples_ppd2$j, var)

######create bins of coastline length#####
#write.csv(subsamples_ppd2,"subs_ppd_case1.csv")
library(dplyr)
library(Hmisc)
subsamples<-subsamples_ppd2%>%mutate(Coastlinebins =
cut(subsamples_ppd2$env_mod, breaks =
c(0,200,400,600,800,1000,1200,1400,1600,1800,2000,2200)))
head(subsamples_ppd2,200)
varbeta=tapply(subsamples$as.numeric.veg., subsamples$Coastlinebins, var)
Coastlinebins=c("200","400","600","800","1000","1200","1400","1600","1800","2000","2200")
var=cbind(Coastline.bins=as.numeric(Coastlinebins),varbeta=as.numeric(round(varbeta,
4)))
variance_ppd=as.data.frame(var)

#######Individual bin method case 2#########################
env<-read.delim("Environment.txt", header=T)
row.names(env)=env[,1]# Declare column 1 as the row names#

s=array(0,dim = c(3000,225))
g=array(0,dim = c(3000,225))
betadata=array(0,dim = c(2,225))
meanbeta=array(0,dim = c(3000,225))
veg=array(0,dim = c(3000,225))
meanppd=vector(mode = "integer", length =1)
env_mod=vector(mode = "integer", length =10)
subsamps <- list()
subsamps<-list()

for(i in 1:50)
{
  env_mod=0
  subsamps1<-list()
  check=sample(1:14,2,replace = F)
  if(check[1]<check[2])
  {
    a=check[1]
    b=check[2]
    if(check[2]-check[1]==1)
    {
      betadata=ta.0[c(a,b),]
      ta.prop <-prop.table(betadata, margin=1)
      veg <- vegdist(ta.prop, method="bray")
      meanbeta<-veg
      j=1
      x=(a+j)-1
      env_mod=env[x,9]
      out=data.frame(i,j,a,b,as.numeric(meanbeta),x,env_mod)
      subsamps1[[i]] <-out
    }else{
      for ( j in 1:((b-a)-1))
      {
```

```
      r=a+j
      t=a+j+1
      s=colSums(ta.0[c(a:r),])
      if(t==b)
      {g=ta.0[t,]}else{
        g=ta.0[c(t:b),]
      }
      betadata=rbind(s,g)
      ta.prop <-prop.table(betadata, margin=1)
      veg <- vegdist(ta.prop, method="bray")
      meanbeta=mean(veg)
      x=(a+j)-1
      env_mod=env_mod+env[x,9]
      out=data.frame(i,j,a,b,as.numeric(meanbeta),x,env_mod)
      subsamps1[[j]] <-out
    }
  }
}else{
  a=check[2]
  b=check[1]
  if(check[1]-check[2]==1)
  {
    betadata=ta.0[c(a,b),]
    ta.prop <-prop.table(betadata, margin=1)
    veg <- vegdist(ta.prop, method="bray")
    meanbeta<-veg
    j=1
    x=(a+j)-1
    env_mod=env[x,9]
    oute=data.frame(i,j,a,b,as.numeric(meanbeta),x,env_mod)
    subsamps1[[j]] <-out
  }else{
    a=check[2]
    b=check[1]
    for ( j in 1:((b-a)-1))
    {
      r=a+j
      t=a+j+1
      s=colSums(ta.0[c(a:r),])
      if(t==b)
      {g=ta.0[t,]}else{
        g=ta.0[c(t:b),]
      }
      betadata=rbind(s,g)
      ta.prop <-prop.table(betadata, margin=1)
      veg <- vegdist(ta.prop, method="bray")
      meanbeta=mean(veg)
      x=(a+j)-1
      env_mod=env_mod+env[x,9]
      out=data.frame(i,j,a,b,as.numeric(meanbeta),x,env_mod)
      subsamps1[[j]] <-out
    }
  }
}
```

```
  subsamples1 <- do.call("rbind",subsamps1)
  subsamps[[i]]=subsamples1
  subsamples_ppd3<- do.call("rbind",subsamps)
}

####variance for bins####
varbeta_ppd3=tapply(subsamples_ppd3$as.numeric.meanbeta., subsamples_ppd3$j,
var)

######create bins of coastline length#####
#write.csv(subsamples_ppd3,"subs_ppd_case2.csv")
library(dplyr)
subsamples<-subsamples_ppd3%>%mutate(Coastlinebins =
cut(subsamples_ppd3$env_mod, breaks =
c(0,200,400,600,800,1000,1200,1400,1600,1800,2000,2200)))
head(subsamples,200)
varbeta=tapply(subsamples$as.numeric.meanbeta., subsamples$Coastlinebins, var)
Coastlinebins=c("200","400","600","800","1000","1200","1400","1600","1800","2000","2200")
var=cbind(Coastline.bins=as.numeric(Coastlinebins),varbeta=as.numeric(round(varbeta,
4)))
variance2_ppd=as.data.frame(var)



##########################################################################
#####Beta diversity whittaker index############
require(vegan)
ta.pa=decostand(ta.01,method = "pa")
d <- betadiver(ta.pa, "w") ##whittaker index####
dppd1<- as.matrix(d)
as.dist(dppd1)
mean(d)
locs1 <- apply(dppd1, 2, FUN=mean, na.rm=T)

####Nullmodel whittaker index####################
#####Combined bin method####################

env<-read.delim("Environment.txt", header=T)
rownames(env)=c("21","20","19","18","17","16","15","14","13","12","11","10","9","8")

#row.names(env)=env[,1]# Declare column 1 as the row names#
s=array(0,dim = c(3000,225))
g=array(0,dim = c(3000,225))
betadata=array(0,dim = c(2,225))
meanbeta=array(0,dim = c(3000,225))
veg=array(0,dim = c(3000,225))
meanppd=vector(mode = "integer", length =1)
env_mod=vector(mode = "integer", length =10)
subsamps <- list()
subsamps<-list()

for(i in 1:50)
{
  env_mod=0
```

```
subsamps1<-list()
check=sample(1:14,2,replace = F)
if(check[1]<check[2])
{
  a=check[1]
  b=check[2]
  if(check[2]-check[1]==1)
  {
    betadata=ta.0[c(a,b),]
    d <- betadiver(betadata, "w")##whittaker index####
    veg <- as.dist(d)
    j=1
    x=(a+j)-1
    env_mod=env[x,9]
    out=data.frame(i,j,a,b,as.numeric(veg),x,env_mod)
    subsamps1[[i]] <-out
  }else{
    for ( j in 1:((b-a)-1))
    {
      r=a+j
      t=a+j+1
      s=colSums(ta.0[c(a:r),])
      if(t==b)
      {g=ta.0[t,]}else{
        g=colSums(ta.0[c(t:b),])
      }
      betadata=rbind(s,g)
      d <- betadiver(betadata, "w")##whittaker index####
      veg <- as.dist(d)
      x=(a+j)-1
      env_mod=env_mod+env[x,9]
      out=data.frame(i,j,a,b,as.numeric(veg),x,env_mod)
      subsamps1[[j]] <-out
    }
  }
}else{
  a=check[2]
  b=check[1]
  if(check[1]-check[2]==1)
  {
    betadata=ta.0[c(a,b),]
    d <- betadiver(betadata, "w")##whittaker index####
    veg <- as.dist(d)
    j=1
    x=(a+j)-1
    env_mod=env[x,9]
    oute=data.frame(i,j,a,b,as.numeric(veg),x,env_mod)
    subsamps1[[j]] <-out
  }else{
    a=check[2]
    b=check[1]
    for ( j in 1:((b-a)-1))
    {
      r=a+j
```

```
        t=a+j+1
        s=colSums(ta.0[c(a:r),])
        if(t==b)
        {g=ta.0[t,]}else{
          g=colSums(ta.0[c(t:b),])
        }
        betadata=rbind(s,g)
        d <- betadiver(betadata, "w")##whittaker index####
        veg <- as.dist(d)
        x=(a+j)-1
        env_mod=env_mod+env[x,9]
        out=data.frame(i,j,a,b,as.numeric(veg),x,env_mod)
        subsamps1[[j]] <-out
      }
    }
  }
  subsamples1 <- do.call("rbind",subsamps1)
  subsamps[[i]]=subsamples1
  subsamples_whit2 <- do.call("rbind",subsamps)
}

###variance for bins##########
varbeta_whit1=tapply(subsamples_whit2$as.numeric.veg., subsamples_whit2$j,
var)

######create bins of coastline length#####
#write.csv(subsamples_whit2,"subs_whit_case1.csv")
library(dplyr)
subsamples_whit2<-subsamples_whit2%>%mutate(Coastlinebins =
cut(subsamples_whit2$env_mod, breaks =
c(0,200,400,600,800,1000,1200,1400,1600,1800,2000,2200)))
head(subsamples_whit2,200)
varbeta=tapply(subsamples_whit2$as.numeric.veg.,
subsamples_whit2$Coastlinebins, var)
write.table(subsamples_whit2$Coastlinebins)
Coastlinebins=c("200","400","600","800","1000","1200","1400","1600","1800","2000","2200")
var=cbind(Coastline.bins=as.numeric(Coastlinebins),varbeta=as.numeric(round(varbeta,
4)))
variance_whit=as.data.frame(var)

####Individual bin method case 2#########
########################clubbing rows from a side only and keeping the
remaining rows#########################################################

env<-read.delim("Environment.txt", header=T)
row.names(env)=env[,1]# Declare column 1 as the row names#

s=array(0,dim = c(3000,225))
g=array(0,dim = c(3000,225))
betadata=array(0,dim = c(2,225))
meanbeta=array(0,dim = c(3000,225))
veg=array(0,dim = c(3000,225))
meanppd=vector(mode = "integer", length =1)
env_mod=vector(mode = "integer", length =10)
```

```
subsamps <- list()
subsamps<-list()

for(i in 1:50)
{
  env_mod=0
  subsamps1<-list()
  check=sample(1:14,2,replace = F)
  if(check[1]<check[2])
  {
    a=check[1]
    b=check[2]
    if(check[2]-check[1]==1)
    {
      betadata=ta.0[c(a,b),]
      d <- betadiver(betadata, "w")##whittaker index####
      veg <- as.dist(d)
      meanbeta<-veg
      j=1
      x=(a+j)-1
      env_mod=env[x,9]
      out=data.frame(i,j,a,b,as.numeric(meanbeta),x,env_mod)
      subsamps1[[i]] <-out
    }else{
      for ( j in 1:((b-a)-1))
      {
        r=a+j
        t=a+j+1
        s=colSums(ta.0[c(a:r),])
        if(t==b)
        {g=ta.0[t,]}else{
          g=ta.0[c(t:b),]
        }
        betadata=rbind(s,g)
        d <- betadiver(betadata, "w")##whittaker index####
        veg <- as.dist(d)
        meanbeta=mean(veg)
        x=(a+j)-1
        env_mod=env_mod+env[x,9]
        out=data.frame(i,j,a,b,as.numeric(meanbeta),x,env_mod)
        subsamps1[[j]] <-out
      }
    }
  }else{
    a=check[2]
    b=check[1]
    if(check[1]-check[2]==1)
    {
      betadata=ta.0[c(a,b),]
      d <- betadiver(betadata, "w")##whittaker index####
      veg <- as.dist(d)
      meanbeta<-veg
      j=1
      x=(a+j)-1
```

```
      env_mod=env[x,9]
      oute=data.frame(i,j,a,b,as.numeric(meanbeta),x,env_mod)
      subsamps1[[j]] <-out
    }else{
      a=check[2]
      b=check[1]
      for ( j in 1:((b-a)-1))
      {
        r=a+j
        t=a+j+1
        s=colSums(ta.0[c(a:r),])
        if(t==b)
        {g=ta.0[t,]}else{
          g=ta.0[c(t:b),]
        }
        betadata=rbind(s,g)
        d <- betadiver(betadata, "w")##whittaker index####
        veg <- as.dist(d)
        meanbeta=mean(veg)
        x=(a+j)-1
        env_mod=env_mod+env[x,9]
        out=data.frame(i,j,a,b,as.numeric(meanbeta),x,env_mod)
        subsamps1[[j]] <-out
      }
    }
  }
  subsamples1 <- do.call("rbind",subsamps1)
  subsamps[[i]]=subsamples1
  subsamples_whit3 <- do.call("rbind",subsamps)
}


varbeta_whit2=tapply(subsamples_whit3$as.numeric.meanbeta.,
subsamples_whit3$j, var)

######create bins of coastline length#####
library(dplyr)
subsamples_whit3<-subsamples_whit3%>%mutate(Coastlinebins =
cut(subsamples_whit3$env_mod, breaks =
c(0,200,400,600,800,1000,1200,1400,1600,1800,2000,2200)))
head(subsamples_whit3,200)
varbeta=tapply(subsamples_whit3$as.numeric.meanbeta.,
subsamples_whit3$Coastlinebins, var)

Coastlinebins=c("200","400","600","800","1000","1200","1400","1600","1800","2000","2200")
var=cbind(Coastline.bins=as.numeric(Coastlinebins),varbeta=as.numeric(round(varbeta,
4)))

variance2_whit=as.data.frame(var)

##############################################
# pairwise beta diversities####
#####simpson######turnover component########
library(betapart)
```

```
ta.pa=decostand(ta.01,method = "pa")
pair.s <- beta.pair(ta.pa)
b_sim <- as.matrix(pair.s$beta.sim)
as.dist(b_sim)
mean(b_sim)
locs_sim <- apply(b_sim, 2, FUN=mean, na.rm=T) #mean beta diversity per site
with regard to other sites #Tables 1,2


#####Null model################
###Combined bin method########
env<-read.delim("Environment.txt", header=T)
rownames(env)=c("21","20","19","18","17","16","15","14","13","12","11","10","9","8")

s=array(0,dim = c(3000,225))
g=array(0,dim = c(3000,225))
betadata=array(0,dim = c(2,225))
meanbeta=array(0,dim = c(3000,225))
veg=array(0,dim = c(3000,225))
meanppd=vector(mode = "integer", length =1)
env_mod=vector(mode = "integer", length =10)
subsamps <- list()
subsamps<-list()

for(i in 1:50)
{
  env_mod=0
  subsamps1<-list()
  check=sample(1:14,2,replace = F)
  if(check[1]<check[2])
  {
    a=check[1]
    b=check[2]
    if(check[2]-check[1]==1)
    {
      betadata=ta.0[c(a,b),]
      betadata.pa=decostand(betadata,method = "pa")
      pair.s <- beta.pair(betadata.pa)
      b_sim <- as.matrix(pair.s$beta.sim)
      veg=as.dist(b_sim)
      j=1
      x=(a+j)-1
      env_mod=env[x,9]
      out=data.frame(i,j,a,b,as.numeric(veg),x,env_mod)
      subsamps1[[i]] <-out
    }else{
      for ( j in 1:((b-a)-1))
      {
        r=a+j
        t=a+j+1
        s=colSums(ta.0[c(a:r),])
        if(t==b)
        {g=ta.0[t,]}else{
          g=colSums(ta.0[c(t:b),])
```

```
      }
      betadata=rbind(s,g)
      betadata.pa=decostand(betadata,method = "pa")
      pair.s <- beta.pair(betadata.pa)
      b_sim <- as.matrix(pair.s$beta.sim)
      veg=as.dist(b_sim)
      x=(a+j)-1
      env_mod=env_mod+env[x,9]
      out=data.frame(i,j,a,b,as.numeric(veg),x,env_mod)
      subsamps1[[j]] <-out
    }
  }
}else{
  a=check[2]
  b=check[1]
  if(check[1]-check[2]==1)
  {
    betadata=ta.0[c(a,b),]
    betadata.pa=decostand(betadata,method = "pa")
    pair.s <- beta.pair(betadata.pa)
    b_sim <- as.matrix(pair.s$beta.sim)
    veg=as.dist(b_sim)
    j=1
    x=(a+j)-1
    env_mod=env[x,9]
    oute=data.frame(i,j,a,b,as.numeric(veg),x,env_mod)
    subsamps1[[j]] <-out
  }else{
    a=check[2]
    b=check[1]
    for ( j in 1:((b-a)-1))
    {
      r=a+j
      t=a+j+1
      s=colSums(ta.0[c(a:r),])
      if(t==b)
      {g=ta.0[t,]}else{
        g=colSums(ta.0[c(t:b),])
      }
      betadata=rbind(s,g)
      betadata.pa=decostand(betadata,method = "pa")
      pair.s <- beta.pair(betadata.pa)
      b_sim <- as.matrix(pair.s$beta.sim)
      veg=as.dist(b_sim)
      x=(a+j)-1
      env_mod=env_mod+env[x,9]
      out=data.frame(i,j,a,b,as.numeric(veg),x,env_mod)
      subsamps1[[j]] <-out
    }
  }
}
subsamples1 <- do.call("rbind",subsamps1)
subsamps[[i]]=subsamples1
subsamples_simp2 <- do.call("rbind",subsamps)
```

```
}


####variance for bins#####
varbeta_simp1=tapply(subsamples_simp2$as.numeric.veg., subsamples_simp2$j,
var)

######create bins of coastline length#####
library(dplyr)
#write.csv(subsamples_simp2,"subs_sim_case1.csv")
subsamples_simp2<-subsamples_simp2%>%mutate(Coastlinebins =
cut(subsamples_simp2$env_mod, breaks =
c(0,200,400,600,800,1000,1200,1400,1600,1800,2000,2200)))
head(subsamples_simp2,200)
varbeta=tapply(subsamples_simp2$as.numeric.veg.,
subsamples_simp2$Coastlinebins, var)
Coastlinebins=c("200","400","600","800","1000","1200","1400","1600","1800","2000","2200")
var=cbind(Coastline.bins=as.numeric(Coastlinebins),varbeta=as.numeric(round(varbeta,
4)))
variance_simp=as.data.frame(var)

#####Individual bin method########################
########################clubbing rows from a side only and keeping the
remaining rows#########################################################

env<-read.delim("Environment.txt", header=T)
row.names(env)=env[,1]# Declare column 1 as the row names#

s=array(0,dim = c(3000,225))
g=array(0,dim = c(3000,225))
betadata=array(0,dim = c(2,225))
meanbeta=array(0,dim = c(3000,225))
veg=array(0,dim = c(3000,225))
meanppd=vector(mode = "integer", length =1)
env_mod=vector(mode = "integer", length =10)
subsamps <- list()
subsamps<-list()

for(i in 1:50)
{
  env_mod=0
  subsamps1<-list()
  check=sample(1:14,2,replace = F)
  if(check[1]<check[2])
  {
    a=check[1]
    b=check[2]
    if(check[2]-check[1]==1)
    {
      betadata=ta.0[c(a,b),]
      betadata.pa=decostand(betadata,method = "pa")
      pair.s <- beta.pair(betadata.pa)
      b_sim <- as.matrix(pair.s$beta.sim)
      meanbeta=as.dist(b_sim)
```

```
      j=1
      x=(a+j)-1
      env_mod=env[x,9]
      out=data.frame(i,j,a,b,as.numeric(meanbeta),x,env_mod)
      subsamps1[[i]] <-out
    }else{
      for ( j in 1:((b-a)-1))
      {
        r=a+j
        t=a+j+1
        s=colSums(ta.0[c(a:r),])
        if(t==b)
        {g=ta.0[t,]}else{
          g=ta.0[c(t:b),]
        }
        betadata=rbind(s,g)
        betadata.pa=decostand(betadata,method = "pa")
        pair.s <- beta.pair(betadata.pa)
        b_sim <- as.matrix(pair.s$beta.sim)
        veg=as.dist(b_sim)
        meanbeta=mean(veg)
        x=(a+j)-1
        env_mod=env_mod+env[x,9]
        out=data.frame(i,j,a,b,as.numeric(meanbeta),x,env_mod)
        subsamps1[[j]] <-out
      }
    }
  }else{
    a=check[2]
    b=check[1]
    if(check[1]-check[2]==1)
    {
      betadata=ta.0[c(a,b),]
      betadata.pa=decostand(betadata,method = "pa")
      pair.s <- beta.pair(betadata.pa)
      b_sim <- as.matrix(pair.s$beta.sim)
      meanbeta=as.dist(b_sim)
      j=1
      x=(a+j)-1
      env_mod=env[x,9]
      oute=data.frame(i,j,a,b,as.numeric(meanbeta),x,env_mod)
      subsamps1[[j]] <-out
    }else{
      a=check[2]
      b=check[1]
      for ( j in 1:((b-a)-1))
      {
        r=a+j
        t=a+j+1
        s=colSums(ta.0[c(a:r),])
        if(t==b)
        {g=ta.0[t,]}else{
          g=ta.0[c(t:b),]
        }
```

```
        betadata=rbind(s,g)
        betadata.pa=decostand(betadata,method = "pa")
        pair.s <- beta.pair(betadata.pa)
        b_sim <- as.matrix(pair.s$beta.sim)
        veg=as.dist(b_sim)
        meanbeta=mean(veg)
        x=(a+j)-1
        env_mod=env_mod+env[x,9]
        out=data.frame(i,j,a,b,as.numeric(meanbeta),x,env_mod)
        subsamps1[[j]] <-out
      }
    }
  }
  subsamples1 <- do.call("rbind",subsamps1)
  subsamps[[i]]=subsamples1
  subsamples_simp3 <- do.call("rbind",subsamps)
}


varbeta_simp2=tapply(subsamples_simp3$as.numeric.meanbeta.,
subsamples_simp3$j, var)

######create bins of coastline length#####
#write.csv(subsamples_simp3,"subs_sim_case2.csv")
library(dplyr)
subsamples_simp3<-subsamples_simp3%>%mutate(Coastlinebins =
cut(subsamples_simp3$env_mod, breaks =
c(0,200,400,600,800,1000,1200,1400,1600,1800,2000,2200)))
head(subsamples_simp3,200)
varbeta=tapply(subsamples_simp3$as.numeric.meanbeta.,
subsamples_simp3$Coastlinebins, var)
Coastlinebins=c("200","400","600","800","1000","1200","1400","1600","1800","2000","2200")
var=cbind(Coastline.bins=as.numeric(Coastlinebins),varbeta=as.numeric(round(varbeta,
4)))
variance2=as.data.frame(var)


#############################################################
####################Sorenson#########total dissimilarity
library(betapart)
ta.pa=decostand(ta.01,method = "pa")
pair.s <- beta.pair(ta.pa)
b_sor<- as.matrix(pair.s$beta.sor)
mean(b_sor)
locs_sor <- apply(b_sor, 2, FUN=mean, na.rm=T) #mean beta diversity per site
with regard to other sites


######Null model##############
##########Combined bin method ###################

env<-read.delim("Environment.txt", header=T)
rownames(env)=c("21","20","19","18","17","16","15","14","13","12","11","10","9","8")
```

```
s=array(0,dim = c(3000,225))
g=array(0,dim = c(3000,225))
betadata=array(0,dim = c(2,225))
meanbeta=array(0,dim = c(3000,225))
veg=array(0,dim = c(3000,225))
meanppd=vector(mode = "integer", length =1)
env_mod=vector(mode = "integer", length =10)
subsamps <- list()
subsamps<-list()

for(i in 1:50)
{
  env_mod=0
  subsamps1<-list()
  check=sample(1:14,2,replace = F)
  if(check[1]<check[2])
  {
    a=check[1]
    b=check[2]
    if(check[2]-check[1]==1)
    {
      betadata=ta.0[c(a,b),]
      betadata.pa=decostand(betadata,method = "pa")
      pair.s <- beta.pair(betadata.pa)
      b_sor <- as.matrix(pair.s$beta.sor)
      veg=as.dist(b_sor)
      j=1
      x=(a+j)-1
      env_mod=env[x,9]
      out=data.frame(i,j,a,b,as.numeric(veg),x,env_mod)
      subsamps1[[i]] <-out
    }else{
      for ( j in 1:((b-a)-1))
      {
        r=a+j
        t=a+j+1
        s=colSums(ta.0[c(a:r),])
        if(t==b)
        {g=ta.0[t,]}else{
          g=colSums(ta.0[c(t:b),])
        }
        betadata=rbind(s,g)
        betadata.pa=decostand(betadata,method = "pa")
        pair.s <- beta.pair(betadata.pa)
        b_sor <- as.matrix(pair.s$beta.sor)
        veg=as.dist(b_sor)
        x=(a+j)-1
        env_mod=env_mod+env[x,9]
        out=data.frame(i,j,a,b,as.numeric(veg),x,env_mod)
        subsamps1[[j]] <-out
      }
    }
  }else{
    a=check[2]
```

```
    b=check[1]
    if(check[1]-check[2]==1)
    {
      betadata=ta.0[c(a,b),]
      betadata.pa=decostand(betadata,method = "pa")
      pair.s <- beta.pair(betadata.pa)
      b_sor<- as.matrix(pair.s$beta.sor)
      veg=as.dist(b_sor)
      j=1
      x=(a+j)-1
      env_mod=env[x,9]
      oute=data.frame(i,j,a,b,as.numeric(veg),x,env_mod)
      subsamps1[[j]] <-out
    }else{
      a=check[2]
      b=check[1]
      for ( j in 1:((b-a)-1))
      {
        r=a+j
        t=a+j+1
        s=colSums(ta.0[c(a:r),])
        if(t==b)
        {g=ta.0[t,]}else{
          g=colSums(ta.0[c(t:b),])
        }
        betadata=rbind(s,g)
        betadata.pa=decostand(betadata,method = "pa")
        pair.s <- beta.pair(betadata.pa)
        b_sor <- as.matrix(pair.s$beta.sor)
        veg=as.dist(b_sor)
        x=(a+j)-1
        env_mod=env_mod+env[x,9]
        out=data.frame(i,j,a,b,as.numeric(veg),x,env_mod)
        subsamps1[[j]] <-out
      }
    }
  }
  subsamples1 <- do.call("rbind",subsamps1)
  subsamps[[i]]=subsamples1
  subsamples_sor2 <- do.call("rbind",subsamps)
}


varbeta_sor=tapply(subsamples_sor2$as.numeric.veg., subsamples_sor2$j, var)

######create bins of coastline length#####
#write.csv(subsamples_sor2,"subs_sor_case1.csv")
library(dplyr)
subsamples_sor2<-subsamples_sor2%>%mutate(Coastlinebins =
cut(subsamples_sor2$env_mod, breaks =
c(0,200,400,600,800,1000,1200,1400,1600,1800,2000,2200)))
head(subsamples_sor2,200)
varbeta=tapply(subsamples_sor2$as.numeric.veg., subsamples_sor2$Coastlinebins,
var)
```

```
Coastlinebins=c("200","400","600","800","1000","1200","1400","1600","1800","2000","2200")
var=cbind(Coastline.bins=as.numeric(Coastlinebins),varbeta=as.numeric(round(varbeta,
4)))
variance_sor=as.data.frame(var)

#######Individual bin method######################################
########################clubbing rows from a side only and keeping the
remaining rows#########################################################

env<-read.delim("Environment.txt", header=T)
row.names(env)=env[,1]

s=array(0,dim = c(3000,225))
g=array(0,dim = c(3000,225))
betadata=array(0,dim = c(2,225))
meanbeta=array(0,dim = c(3000,225))
veg=array(0,dim = c(3000,225))
meanppd=vector(mode = "integer", length =1)
env_mod=vector(mode = "integer", length =10)
subsamps <- list()
subsamps<-list()

for(i in 1:50)
{
  env_mod=0
  subsamps1<-list()
  check=sample(1:14,2,replace = F)
  if(check[1]<check[2])
  {
    a=check[1]
    b=check[2]
    if(check[2]-check[1]==1)
    {
      betadata=ta.0[c(a,b),]
      betadata.pa=decostand(betadata,method = "pa")
      pair.s <- beta.pair(betadata.pa)
      b_sor <- as.matrix(pair.s$beta.sor)
      meanbeta=as.dist(b_sor)
      j=1
      x=(a+j)-1
      env_mod=env[x,9]
      out=data.frame(i,j,a,b,as.numeric(meanbeta),x,env_mod)
      subsamps1[[i]] <-out
    }else{
      for ( j in 1:((b-a)-1))
      {
        r=a+j
        t=a+j+1
        s=colSums(ta.0[c(a:r),])
        if(t==b)
        {g=ta.0[t,]}else{
          g=ta.0[c(t:b),]
        }
        betadata=rbind(s,g)
```

```
      betadata.pa=decostand(betadata,method = "pa")
      pair.s <- beta.pair(betadata.pa)
      b_sor <- as.matrix(pair.s$beta.sor)
      veg=as.dist(b_sor)
      meanbeta=mean(veg)
      x=(a+j)-1
      env_mod=env_mod+env[x,9]
      out=data.frame(i,j,a,b,as.numeric(meanbeta),x,env_mod)
      subsamps1[[j]] <-out
    }
  }
}else{
  a=check[2]
  b=check[1]
  if(check[1]-check[2]==1)
  {
    betadata=ta.0[c(a,b),]
    betadata.pa=decostand(betadata,method = "pa")
    pair.s <- beta.pair(betadata.pa)
    b_sor <- as.matrix(pair.s$beta.sor)
    meanbeta=as.dist(b_sor)
    j=1
    x=(a+j)-1
    env_mod=env[x,9]
    oute=data.frame(i,j,a,b,as.numeric(meanbeta),x,env_mod)
    subsamps1[[j]] <-out
  }else{
    a=check[2]
    b=check[1]
    for ( j in 1:((b-a)-1))
    {
      r=a+j
      t=a+j+1
      s=colSums(ta.0[c(a:r),])
      if(t==b)
      {g=ta.0[t,]}else{
        g=ta.0[c(t:b),]
      }
      betadata=rbind(s,g)
      betadata.pa=decostand(betadata,method = "pa")
      pair.s <- beta.pair(betadata.pa)
      b_sor <- as.matrix(pair.s$beta.sor)
      veg=as.dist(b_sor)
      meanbeta=mean(veg)
      x=(a+j)-1
      env_mod=env_mod+env[x,9]
      out=data.frame(i,j,a,b,as.numeric(meanbeta),x,env_mod)
      subsamps1[[j]] <-out
    }
  }
}
subsamples1 <- do.call("rbind",subsamps1)
subsamps[[i]]=subsamples1
subsamples_sor3 <- do.call("rbind",subsamps)
```

```
}


varbeta_sor2=tapply(subsamples_sor3$as.numeric.meanbeta., subsamples_sor3$j,
var)

######create bins of coastline length#####
library(dplyr)
#write.csv(subsamples_sor3,"subsamples_sor_case2.csv")
subsamples_sor3<-subsamples_sor3%>%mutate(Coastlinebins =
cut(subsamples_sor3$env_mod, breaks =
c(0,200,400,600,800,1000,1200,1400,1600,1800,2000,2200)))
head(subsamples_sor3,200)
varbeta=tapply(subsamples_sor3$as.numeric.meanbeta.,
subsamples_sor3$Coastlinebins, var)
Coastlinebins=c("200","400","600","800","1000","1200","1400","1600","1800","2000","2200")
var=cbind(Coastline.bins=as.numeric(Coastlinebins),varbeta=as.numeric(round(varbeta,
4)))
variance2_sor=as.data.frame(var)

#################nestedness component of sorenson###########
library(betapart)
library(vegan)
ta.pa=decostand(ta.01,method = "pa")
pair.s <- beta.pair(ta.pa)
b_sne<- as.matrix(pair.s$beta.sne)
mean(b_sne)
locs_sne <- apply(b_sne, 2, FUN=mean, na.rm=T) #mean beta diversity per site
with regard to other sites #Tables 1,2
x1 = matrix(c(rownames(ta.01)),nrow=14,ncol=1)


####Null model#############
##########Combined bin method###############
env<-read.delim("Environment.txt", header=T)
rownames(env)=c("21","20","19","18","17","16","15","14","13","12","11","10","9","8")

s=array(0,dim = c(3000,225))
g=array(0,dim = c(3000,225))
betadata=array(0,dim = c(2,225))
meanbeta=array(0,dim = c(3000,225))
veg=array(0,dim = c(3000,225))
meanppd=vector(mode = "integer", length =1)
env_mod=vector(mode = "integer", length =10)
subsamps <- list()
subsamps<-list()

for(i in 1:50)
{
  env_mod=0
  subsamps1<-list()
  check=sample(1:14,2,replace = F)
  if(check[1]<check[2])
  {
```

```
    a=check[1]
    b=check[2]
    if(check[2]-check[1]==1)
    {
      betadata=ta.0[c(a,b),]
      betadata.pa=decostand(betadata,method = "pa")
      pair.s <- beta.pair(betadata.pa)
      b_sne <- as.matrix(pair.s$beta.sne)
      veg=as.dist(b_sne)
      j=1
      x=(a+j)-1
      env_mod=env[x,9]
      out=data.frame(i,j,a,b,as.numeric(veg),x,env_mod)
      subsamps1[[i]] <-out
    }else{
      for ( j in 1:((b-a)-1))
      {
        r=a+j
        t=a+j+1
        s=colSums(ta.0[c(a:r),])
        if(t==b)
        {g=ta.0[t,]}else{
          g=colSums(ta.0[c(t:b),])
        }
        betadata=rbind(s,g)
        betadata.pa=decostand(betadata,method = "pa")
        pair.s <- beta.pair(betadata.pa)
        b_sne <- as.matrix(pair.s$beta.sne)
        veg=as.dist(b_sne)
        x=(a+j)-1
        env_mod=env_mod+env[x,9]
        out=data.frame(i,j,a,b,as.numeric(veg),x,env_mod)
        subsamps1[[j]] <-out
      }
    }
  }else{
    a=check[2]
    b=check[1]
    if(check[1]-check[2]==1)
    {
      betadata=ta.0[c(a,b),]
      betadata.pa=decostand(betadata,method = "pa")
      pair.s <- beta.pair(betadata.pa)
      b_sne<- as.matrix(pair.s$beta.sne)
      veg=as.dist(b_sne)
      j=1
      x=(a+j)-1
      env_mod=env[x,9]
      oute=data.frame(i,j,a,b,as.numeric(veg),x,env_mod)
      subsamps1[[j]] <-out
    }else{
      a=check[2]
      b=check[1]
      for ( j in 1:((b-a)-1))
```

```
      {
        r=a+j
        t=a+j+1
        s=colSums(ta.0[c(a:r),])
        if(t==b)
        {g=ta.0[t,]}else{
          g=colSums(ta.0[c(t:b),])
        }
        betadata=rbind(s,g)
        betadata.pa=decostand(betadata,method = "pa")
        pair.s <- beta.pair(betadata.pa)
        b_sne <- as.matrix(pair.s$beta.sne)
        veg=as.dist(b_sne)
        x=(a+j)-1
        env_mod=env_mod+env[x,9]
        out=data.frame(i,j,a,b,as.numeric(veg),x,env_mod)
        subsamps1[[j]] <-out
      }
    }
  }
  subsamples1 <- do.call("rbind",subsamps1)
  subsamps[[i]]=subsamples1
  subsamples_sne2 <- do.call("rbind",subsamps)
}



varbeta_sne=tapply(subsamples_sne2$as.numeric.veg., subsamples_sne2$j, var)

######create bins of coastline length#####
#write.csv(subsamples_sne2,"subsamples_sne_case1.csv")
library(dplyr)
subsamples_sne2<-subsamples_sne2%>%mutate(Coastlinebins =
cut(subsamples_sne2$env_mod, breaks =
c(0,200,400,600,800,1000,1200,1400,1600,1800,2000,2200)))
head(subsamples_sne2,200)
varbeta=tapply(subsamples_sne2$as.numeric.veg., subsamples_sne2$Coastlinebins,
var)
Coastlinebins=c("200","400","600","800","1000","1200","1400","1600","1800","2000","2200")
var=cbind(Coastline.bins=as.numeric(Coastlinebins),varbeta=as.numeric(round(varbeta,
4)))
variance_sne=as.data.frame(var)

####Individual bin method##############################
#######################clubbing rows from a side only and keeping the
remaining rows########################################################

env<-read.delim("Environment.txt", header=T)
row.names(env)=env[,1]# Declare column 1 as the row names#

s=array(0,dim = c(3000,225))
g=array(0,dim = c(3000,225))
betadata=array(0,dim = c(2,225))
meanbeta=array(0,dim = c(3000,225))
```

```
veg=array(0,dim = c(3000,225))
meanppd=vector(mode = "integer", length =1)
env_mod=vector(mode = "integer", length =10)
subsamps <- list()
subsamps<-list()

for(i in 1:50)
{
  env_mod=0
  subsamps1<-list()
  check=sample(1:14,2,replace = F)
  if(check[1]<check[2])
  {
    a=check[1]
    b=check[2]
    if(check[2]-check[1]==1)
    {
      betadata=ta.0[c(a,b),]
      betadata.pa=decostand(betadata,method = "pa")
      pair.s <- beta.pair(betadata.pa)
      b_sne <- as.matrix(pair.s$beta.sne)
      meanbeta=as.dist(b_sne)
      j=1
      x=(a+j)-1
      env_mod=env[x,9]
      out=data.frame(i,j,a,b,as.numeric(meanbeta),x,env_mod)
      subsamps1[[i]] <-out
    }else{
      for ( j in 1:((b-a)-1))
      {
        r=a+j
        t=a+j+1
        s=colSums(ta.0[c(a:r),])
        if(t==b)
        {g=ta.0[t,]}else{
          g=ta.0[c(t:b),]
        }
        betadata=rbind(s,g)
        betadata.pa=decostand(betadata,method = "pa")
        pair.s <- beta.pair(betadata.pa)
        b_sne <- as.matrix(pair.s$beta.sne)
        veg=as.dist(b_sne)
        meanbeta=mean(veg)
        x=(a+j)-1
        env_mod=env_mod+env[x,9]
        out=data.frame(i,j,a,b,as.numeric(meanbeta),x,env_mod)
        subsamps1[[j]] <-out
      }
    }
  }else{
    a=check[2]
    b=check[1]
    if(check[1]-check[2]==1)
    {
```

```
      betadata=ta.0[c(a,b),]
      betadata.pa=decostand(betadata,method = "pa")
      pair.s <- beta.pair(betadata.pa)
      b_sne <- as.matrix(pair.s$beta.sne)
      meanbeta=as.dist(b_sne)
      j=1
      x=(a+j)-1
      env_mod=env[x,9]
      oute=data.frame(i,j,a,b,as.numeric(meanbeta),x,env_mod)
      subsamps1[[j]] <-out
    }else{
      a=check[2]
      b=check[1]
      for ( j in 1:((b-a)-1))
      {
        r=a+j
        t=a+j+1
        s=colSums(ta.0[c(a:r),])
        if(t==b)
        {g=ta.0[t,]}else{
          g=ta.0[c(t:b),]
        }
        betadata=rbind(s,g)
        betadata.pa=decostand(betadata,method = "pa")
        pair.s <- beta.pair(betadata.pa)
        b_sne <- as.matrix(pair.s$beta.sne)
        veg=as.dist(b_sne)
        meanbeta=mean(veg)
        x=(a+j)-1
        env_mod=env_mod+env[x,9]
        out=data.frame(i,j,a,b,as.numeric(meanbeta),x,env_mod)
        subsamps1[[j]] <-out
      }
    }
  }
  subsamples1 <- do.call("rbind",subsamps1)
  subsamps[[i]]=subsamples1
  subsamples_sne3 <- do.call("rbind",subsamps)
}


varbeta_sne2=tapply(subsamples_sne3$as.numeric.meanbeta., subsamples_sne3$j,
var)

######create bins of coastline length#####
#write.csv(subsamples_sne3,"subsamples_sne_case2.csv")
library(dplyr)
subsamples_sne3<-subsamples_sne3%>%mutate(Coastlinebins =
cut(subsamples_sne3$env_mod, breaks =
c(0,200,400,600,800,1000,1200,1400,1600,1800,2000,2200)))
head(subsamples_sne3,200)
varbeta=tapply(subsamples_sne3$as.numeric.meanbeta.,
subsamples_sne3$Coastlinebins, var)
Coastlinebins=c("200","400","600","800","1000","1200","1400","1600","1800","2000","2200")
```

```
var=cbind(Coastline.bins=as.numeric(Coastlinebins),varbeta=as.numeric(round(varbeta,
4)))
variance2_sne=as.data.frame(var)

#########Table 3.2############################
######Correlation values###############
############################################################################
par(mfrow=c(5,2))
par(mar=c(2,5,1,1))
#par(pty="s")
#####with boxes####
######ppd case 1################
plot(subsamples_ppd2$j,subsamples_ppd2$as.numeric.veg., col="black",
ylim=c(0.1,1),ylab=expression (beta[ppd]),xlab="Number of boxes/ lat bins")
cor.test(subsamples_ppd2$j,subsamples_ppd2$as.numeric.veg.,method="spearman")
####ppd case 2###############
plot(subsamples_ppd3$j,subsamples_ppd3$as.numeric.meanbeta., col="black",
ylim=c(0.1,1),ylab=expression (beta[ppd]),xlab="Number of boxes/ lat bins")
cor.test(subsamples_ppd3$j,subsamples_ppd3$as.numeric.meanbeta.,method="spearman")

############whit case1
plot(subsamples_whit2$j,subsamples_whit2$as.numeric.veg., col="black",
ylim=c(0.1,1),ylab=expression (beta[whit]),xlab="Number of boxes/ lat bins")
cor.test(subsamples_whit2$as.numeric.veg.,subsamples_whit2$j,method="spearman")
#######whit case 2
plot(subsamples_whit3$j,subsamples_whit3$as.numeric.meanbeta., col="black",
ylim=c(0.1,1),ylab=expression (beta[whit]),xlab="Number of boxes/ lat bins")
cor.test(subsamples_whit3$as.numeric.meanbeta.,subsamples_whit3$j,method="spearman")

###simpson case 1
plot(subsamples_simp2$j,subsamples_simp2$as.numeric.veg., col="black",
ylim=c(0,1),ylab=expression (beta[sim]),xlab="Number of boxes/ lat bins")
cor.test(subsamples_simp2$as.numeric.veg.,subsamples_simp2$j,method="spearman")
####simpson case 2
plot(subsamples_simp3$j,subsamples_simp3$as.numeric.meanbeta., col="black",
ylim=c(0,1),ylab=expression (beta[sim]),xlab="Number of boxes/ lat bins")
cor.test(subsamples_simp3$as.numeric.meanbeta.,subsamples_simp3$j,method="spearman")

###sorenson case 1
plot(subsamples_sor2$j,subsamples_sor2$as.numeric.veg., col="black",
ylim=c(0.1,1),ylab=expression (beta[sor]),xlab="Number of boxes/ lat bins")
cor.test(subsamples_sor2$as.numeric.veg.,subsamples_sor2$j,method="spearman")
#sorenson case 2
plot(subsamples_sor3$j,subsamples_sor3$as.numeric.meanbeta., col="black",
ylim=c(0.1,1),ylab=expression (beta[sor]),xlab="Number of boxes/ lat bins")
cor.test(subsamples_sor3$as.numeric.meanbeta.,subsamples_sor3$j,method="spearman")

#sne case 1
plot(subsamples_sne2$j,subsamples_sne2$as.numeric.veg., col="black",
ylim=c(0.1,1),ylab=expression (beta[sne]),xlab="Number of boxes/ lat bins")
cor.test(subsamples_sne2$j,subsamples_sne2$as.numeric.veg.,method="spearman")
##sne case 2
plot(subsamples_sne3$j,subsamples_sne3$as.numeric.meanbeta., col="black",
ylim=c(0.1,1),ylab=expression (beta[sne]),xlab="Number of boxes/ lat bins")
```

```
cor.test(subsamples_sne3$j,subsamples_sne3$as.numeric.meanbeta.,method="spearman")


#############with coastline length##########################################
##ppd case 1
plot(subsamples_ppd2$env_mod,subsamples_ppd2$as.numeric.veg., col="black",
ylim=c(0.5,1),ylab=expression (beta[ppd]),xlab="Coastline length")
cor.test(subsamples_ppd2$env_mod,subsamples_ppd2$as.numeric.veg.,method="spearman")
####ppd case 2
plot(subsamples_ppd3$env_mod,subsamples_ppd3$as.numeric.meanbeta.,
col="black", ylim=c(0.5,1),ylab=expression (beta[ppd]),xlab="Coastline
length")
cor.test(subsamples_ppd3$env_mod,subsamples_ppd3$as.numeric.meanbeta.,method="spearman")

######whit case 1
plot(subsamples_whit2$env_mod,subsamples_whit2$as.numeric.veg., col="black",
ylim=c(0.1,1),ylab=expression (beta[whit]),xlab="Coastline length")
cor.test(subsamples_whit2$env_mod,subsamples_whit2$as.numeric.veg.,method="spearman")
#######whit case 2
plot(subsamples_whit3$env_mod,subsamples_whit3$as.numeric.meanbeta.,
col="black", ylim=c(0.1,1),ylab=expression (beta[whit]),xlab="Coastline
length")
cor.test(subsamples_whit3$env_mod,subsamples_whit3$as.numeric.meanbeta.,method="spearman")

###simpson case 1
plot(subsamples_simp2$env_mod,subsamples_simp2$as.numeric.veg., col="black",
ylim=c(0,1),ylab=expression (beta[sim]),xlab="Coastline length")
cor.test(subsamples_simp2$env_mod,subsamples_simp2$as.numeric.veg.,method="spearman")
#####simpson case 2
plot(subsamples_simp3$env_mod,subsamples_simp3$as.numeric.meanbeta.,
col="black", ylim=c(0,1),ylab=expression (beta[sim]),xlab="Coastline length")
cor.test(subsamples_simp3$env_mod,subsamples_simp3$as.numeric.meanbeta.,method="spearman")

####sorenson case 1
plot(subsamples_sor2$env_mod,subsamples_sor2$as.numeric.veg., col="black",
ylim=c(0,1),ylab=expression (beta[sor]),xlab="Coastline length")
cor.test(subsamples_sor2$env_mod,subsamples_sor2$as.numeric.veg.,method="spearman")
###sorenson casse 2
plot(subsamples_sor3$env_mod,subsamples_sor3$as.numeric.meanbeta.,
col="black", ylim=c(0,1),ylab=expression (beta[sor]),xlab="Coastline length")
cor.test(subsamples_sor3$env_mod,subsamples_sor3$as.numeric.meanbeta.,method="spearman")

####sne case 1
plot(subsamples_sne2$env_mod,subsamples_sne2$as.numeric.veg., col="black",
ylim=c(0,1),ylab=expression (beta[sne]),xlab="Coastline length")
cor.test(subsamples_sne2$env_mod,subsamples_sne2$as.numeric.veg.,method="spearman")

##sne case 2
plot(subsamples_sne3$env_mod,subsamples_sne3$as.numeric.meanbeta.,
col="black", ylim=c(0,1),ylab=expression (beta[sne]),xlab="Coastline length")
cor.test(subsamples_sne3$env_mod,subsamples_sne3$as.numeric.meanbeta.,method="spearman")


######################variance########################################################
```

```
#ppd case 1
plot(1:length(varbeta_ppd),varbeta_ppd, pch=19, ylab=expression
(beta[ppd]~variance),xlab="Number of latitude bins/boxes")
cor.test(1:length(varbeta_ppd),varbeta_ppd,method="spearman")

plot(variance_ppd$Coastline.bins,variance_ppd$varbeta, pch=19, ylab=expression
(beta[ppd]~variance),xlab = "Coastline length")
cor.test(variance_ppd$Coastline.bins,variance_ppd$varbeta,method = "spearman")

###ppd case 2
plot( 1:length(varbeta_ppd3),varbeta_ppd3,pch=19, ylab=expression
(beta[ppd]~variance),xlab="Number of latitude bins/boxes")
cor.test(varbeta_ppd3,1:length(varbeta_ppd3),method = "spearman")

plot(variance2_ppd$Coastline.bins,variance2_ppd$varbeta,pch=19,
ylab=expression (beta[ppd]~variance),xlab = "Coastline length")
cor.test(variance2_ppd$Coastline.bins,variance2_ppd$varbeta,method =
"spearman")

#whit case 1#############
plot(1:length(varbeta_whit1),varbeta_whit1,pch=19, ylab=expression
(beta[whit]~variance),xlab="Number of latitude bins/boxes")
cor.test(1:length(varbeta_whit1),varbeta_whit1,method = "spearman")
#points(subsamples$j, subsamples$as.numeric.veg., ylim=c(0,9))

plot(variance_whit$Coastline.bins,variance_whit$varbeta,pch=19,
ylab=expression (beta[whit]~variance),xlab = "Coastline length")
cor.test(variance_whit$Coastline.bins,variance_whit$varbeta,method =
"spearman")

##whit case 2####################################
plot(1:length(varbeta_whit2),varbeta_whit2, pch=19, ylab=expression
(beta[whit]~variance),xlab="Number of latitude bins/boxes")
cor.test(1:length(varbeta_whit2),varbeta_whit2,method = "spearman")

plot(variance2_whit$Coastline.bins,variance2_whit$varbeta,pch=19,
ylab=expression (beta[whit]~variance),xlab = "Coastline length")
cor.test(variance2_whit$Coastline.bins,variance2_whit$varbeta,method =
"spearman")

######simp case 1##############
cor.test(1:length(varbeta_simp1),varbeta_simp1,method="spearman")
plot(1:length(varbeta_simp1),varbeta_simp1, pch=19, ylab=expression
(beta[simp]~variance),xlab="Number of latitude bins/boxes")

plot(variance_simp$Coastline.bins,variance_simp$varbeta,pch=19,
ylab=expression (beta[simp]~variance),xlab = "Coastline length")
cor.test(variance_simp$Coastline.bins,variance_simp$varbeta,method =
"spearman")

########simp case 2
plot(1:length(varbeta_simp2),varbeta_simp2, pch=19, ylab=expression
(beta[simp]~variance),xlab="Number of latitude bins/boxes")
cor.test(1:length(varbeta_simp2),varbeta_simp2,method = "spearman")
```

```
plot(variance2$Coastline.bins,variance2$varbeta,pch=19, ylab=expression
(beta[simp]~variance),xlab = "Coastline length")
cor.test(variance2$Coastline.bins,variance2$varbeta,method = "spearman")

#######sorenson case 1
cor.test(1:length(varbeta_sor),varbeta_sor,method="spearman")
plot(1:length(varbeta_sor),varbeta_sor, pch=19, ylab=expression
(beta[sor]~variance),xlab="Number of latitude bins/boxes")

plot(variance_sor$Coastline.bins,variance_sor$varbeta,pch=19,ylab=expression
(beta[sor]~variance),xlab = "Coastline length")
cor.test(variance_sor$Coastline.bins,variance_sor$varbeta,method = "spearman")

###sorenson case 2
plot(1:length(varbeta_sor2),varbeta_sor2,pch=19,ylab=expression
(beta[sor]~variance),xlab="Number of latitude bins/boxes")
cor.test(varbeta_sor2,c(1:length(varbeta_sor2)),method = "spearman")

plot(variance2_sor$Coastline.bins,variance2_sor$varbeta,pch=19,
ylab=expression (beta[sor]~variance),xlab = "Coastline length")
cor.test(variance2_sor$Coastline.bins,variance2_sor$varbeta,method =
"spearman")

###sne case 1
cor.test(1:length(varbeta_sne),varbeta_sne,method="spearman")
plot(1:length(varbeta_sne),varbeta_sne, pch=19,ylab=expression
(beta[sne]~variance),xlab="Number of latitude bins/boxes")

plot(variance_sne$Coastline.bins,variance_sne$varbeta,pch=19, ylab=expression
(beta[sne]~variance),xlab = "Coastline length")
cor.test(variance_sne$Coastline.bins,variance_sne$varbeta,method = "spearman")

###sne case 2
plot(1:length(varbeta_sne2),varbeta_sne2,pch=19, ylab=expression
(beta[sne]~variance),xlab="Number of latitude bins/boxes")
cor.test(varbeta_sne2,1:length(varbeta_sne2),method = "spearman")

plot(variance2_sne$Coastline.bins,variance2_sne$varbeta,pch=19,ylab=expression
(beta[sne]~variance),xlab = "Coastline length")
cor.test(variance2_sne$Coastline.bins,variance2_sne$varbeta,method =
"spearman")



#Figure 3.3#######################################
#############with coastline length###########################################
##ppd case 1
par(mfrow=c(5,2))
par(mar=c(2,5,1,3))
plot(subsamples_ppd2$env_mod,subsamples_ppd2$as.numeric.veg.,ylim=c(0.1,1),
xlim=c(0,2500), col="black", ylab=expression (beta[ppd]),xlab="Coastline
length",las=1,cex.lab=1.5)
par(new=TRUE)
```

```
plot(variance_ppd$Coastline.bins,variance_ppd$varbeta,
axes=F,type="b",pch=19,col="red",xlab=" ",ylab =" ")
axis(4,ylim=c(0,0.1),las=1)

####ppd case 2
par(mar=c(2,3,1,6))
plot(subsamples_ppd3$env_mod,subsamples_ppd3$as.numeric.meanbeta.,
xlim=c(0,2500),col="black", ylim=c(0.1,1),ylab=" ",xlab="Coastline
length",las=1)
par(new=TRUE)
plot(variance2_ppd$Coastline.bins,variance2_ppd$varbeta,
axes=F,type="b",pch=19,col="red",xlab=" ",ylab =" ")
axis(4,ylim=c(0,0.1),las=1)
mtext(expression (variance~beta[ppd]),side=4,cex = 1,line=4)

######whit case 1
par(mar=c(2,5,1,3))
plot(subsamples_whit2$env_mod,subsamples_whit2$as.numeric.veg.,xlim=c(0,2500),
col="black", ylim=c(0.1,1),ylab=expression (beta[whit]),xlab="Coastline
length",las=1,cex.lab=1.5)
par(new=TRUE)
plot(variance_whit$Coastline.bins,variance_whit$varbeta,
axes=F,type="b",pch=19,col="red",xlab=" ",ylab =" ")
axis(4,ylim=c(0,0.1),las=1)
#######whit case 2
par(mar=c(2,3,1,6))
plot(subsamples_whit3$env_mod,subsamples_whit3$as.numeric.meanbeta.,xlim=c(0,2500),
col="black", ylim=c(0.1,1),ylab=" ",xlab="Coastline length",las=1)
par(new=TRUE)
plot(variance2_whit$Coastline.bins,variance2_whit$varbeta,
axes=F,type="b",pch=19,col="red",xlab=" ",ylab =" ")
axis(4,ylim=c(0,0.1),las=1)
mtext(expression (variance~beta[whit]),side=4,col="black",cex = 1,line=4)

###simpson case 1
par(mar=c(2,5,1,3))
plot(subsamples_simp2$env_mod,subsamples_simp2$as.numeric.veg.,xlim=c(0,2500),
col="black", ylim=c(0.1,1),ylab=expression (beta[sim]),xlab="Coastline
length",las=1,cex.lab=1.5)
par(new=TRUE)
plot(variance_simp$Coastline.bins,variance_simp$varbeta,
axes=F,type="b",pch=19,col="red",xlab=" ",ylab =" ")
axis(4,ylim=c(0,0.1),las=1)
####simpson case 2
par(mar=c(2,3,1,6))
plot(subsamples_simp3$env_mod,subsamples_simp3$as.numeric.meanbeta.,xlim=c(0,2500),
col="black", ylim=c(0.1,1),ylab=" ",xlab="Coastline length",las=1)
par(new=TRUE)
plot(variance2$Coastline.bins,variance2$varbeta,
axes=F,type="b",pch=19,col="red",xlab=" ",ylab =" ")
axis(4,ylim=c(0,0.1),las=1)
mtext(expression (variance~beta[sim]),side=4,col="black",cex = 1,line=4)

###sorenson case 1
```

```
par(mar=c(2,5,1,3))
plot(subsamples_sor2$env_mod,subsamples_sor2$as.numeric.veg.,xlim=c(0,2500),
col="black", ylab=expression (beta[sor]),xlab="Coastline
length",las=1,cex.lab=1.5)
par(new=TRUE)
plot(variance_sor$Coastline.bins,variance_sor$varbeta,
axes=F,type="b",pch=19,col="red",xlab=" ",ylab =" ")
axis(4,ylim=c(0,0.1),las=1)
###sorenson casse 2
par(mar=c(2,3,1,6))
plot(subsamples_sor3$env_mod,subsamples_sor3$as.numeric.meanbeta.,xlim=c(0,2500),
col="black", ylim=c(0.1,1),ylab=" ",xlab="Coastline length",las=1)
par(new=TRUE)
plot(variance2_sor$Coastline.bins,variance2_sor$varbeta,
axes=F,type="b",pch=19,col="red",xlab=" ",ylab =" ")
axis(4,ylim=c(0,0.1),las=1)
mtext(expression (variance~beta[sor]),side=4,col="black",cex = 1,line=4)

####sne case 1
par(mar=c(2,5,1,3))
plot(subsamples_sne2$env_mod,subsamples_sne2$as.numeric.veg.,
xlim=c(0,2500),col="black", ylim=c(0.1,1),ylab=expression
(beta[sne]),xlab="Coastline length",las=1,cex.lab=1.5)
par(new=TRUE)
plot(variance_sne$Coastline.bins,variance_sne$varbeta,
axes=F,type="b",pch=19,col="red",xlab=" ",ylab =" ")
axis(4,ylim=c(0,0.1),las=1)

##sne case 2
par(mar=c(2,3,1,6))
plot(subsamples_sne3$env_mod,subsamples_sne3$as.numeric.meanbeta.,
xlim=c(0,2500),col="black", ylim=c(0.1,1),ylab=" ",xlab="Coastline
length",las=1)
par(new=TRUE)
plot(variance2_sne$Coastline.bins,variance2_sne$varbeta,
axes=F,type="b",pch=19,col="red",xlab=" ",ylab =" ")
axis(4,ylim=c(0,0.1),las=1)
mtext(expression (variance~beta[sne]),side=4,col="black",cex = 1,line=4)

###############plots##########################
############Figure 3.S1##########################
#####with boxes####
par(mfrow=c(5,2))
par(mar=c(2,5,1,3))
######ppd case 1################
plot(subsamples_ppd2$j,subsamples_ppd2$as.numeric.veg., col="black",
ylim=c(0.1,1),ylab=expression (beta[ppd]),xlab="Number of boxes/ lat
bins",las=1,cex.lab=1.5)
par(new=TRUE)
plot(1:length(varbeta_ppd),varbeta_ppd,
axes=F,type="b",pch=19,col="red",xlab=" ",ylab =" ")
axis(4,ylim=c(0,0.05),las=1)

####ppd case 2##############
```

```
par(mar=c(2,3,1,6))
plot(subsamples_ppd3$j,subsamples_ppd3$as.numeric.meanbeta., col="black",
ylim=c(0.1,1),ylab=" ",xlab="Number of boxes/ lat bins",las=1)
par(new=TRUE)
plot(1:length(varbeta_ppd3),varbeta_ppd3,
axes=F,type="b",pch=19,col="red",xlab=" ",ylab =" ")
axis(4,ylim=c(0,0.1),las=1)
mtext(expression (variance~beta[ppd]),side=4,cex = 1,line=4)

############whit case1
par(mar=c(2,5,1,3))
plot(subsamples_whit2$j,subsamples_whit2$as.numeric.veg., col="black",
ylim=c(0.1,1),ylab=expression (beta[whit]),xlab="Number of boxes/ lat
bins",las=1,cex.lab=1.5)
par(new=TRUE)
plot(1:length(varbeta_whit1),varbeta_whit1,
axes=F,type="b",pch=19,col="red",xlab=" ",ylab =" ")
axis(4,ylim=c(0,0.05),las=1)
#######whit case 2
par(mar=c(2,3,1,6))
plot(subsamples_whit3$j,subsamples_whit3$as.numeric.meanbeta., col="black",
ylim=c(0.1,1),ylab=" ",xlab="Number of boxes/ lat bins",las=1)
par(new=TRUE)
plot(1:length(varbeta_whit2),varbeta_whit2,
axes=F,type="b",pch=19,col="red",xlab=" ",ylab =" ")
axis(4,ylim=c(0,0.1),las=1)
mtext(expression (variance~beta[whit]),side=4,col="black",cex = 1,line=4)

###simpson case 1
par(mar=c(2,5,1,3))
plot(subsamples_simp2$j,subsamples_simp2$as.numeric.veg., col="black",
ylim=c(0.1,1),ylab=expression (beta[sim]),xlab="Number of boxes/ lat
bins",las=1,cex.lab=1.5)
par(new=TRUE)
plot(1:length(varbeta_simp1),varbeta_simp1,
axes=F,type="b",pch=19,col="red",xlab=" ",ylab =" ")
axis(4,ylim=c(0,0.05),las=1)
####simpson case 2
par(mar=c(2,3,1,6))
plot(subsamples_simp3$j,subsamples_simp3$as.numeric.meanbeta., col="black",
ylim=c(0.1,1),ylab=" ",xlab="Number of boxes/ lat bins",las=1)
par(new=TRUE)
plot(1:length(varbeta_simp2),varbeta_simp2,
axes=F,type="b",pch=19,col="red",xlab=" ",ylab =" ")
axis(4,ylim=c(0,0.1),las=1)
mtext(expression (variance~beta[sim]),side=4,col="black",cex = 1,line=4)

###sorenson case 1
par(mar=c(2,5,1,3))
plot(subsamples_sor2$j,subsamples_sor2$as.numeric.veg., col="black",
ylim=c(0.1,1),ylab=expression (beta[sor]),xlab="Number of boxes/ lat
bins",las=1,cex.lab=1.5)
par(new=TRUE)
```

```
plot(1:length(varbeta_sor),varbeta_sor,
axes=F,type="b",pch=19,col="red",xlab=" ",ylab =" ")
axis(4,ylim=c(0,0.05),las=1)
#sorenson case 2
par(mar=c(2,3,1,6))
plot(subsamples_sor3$j,subsamples_sor3$as.numeric.meanbeta., col="black",
ylim=c(0.1,1),ylab=" ",xlab="Number of boxes/ lat bins",las=1)
par(new=TRUE)
plot(1:length(varbeta_sor2),varbeta_sor2,
axes=F,type="b",pch=19,col="red",xlab=" ",ylab =" ")
axis(4,ylim=c(0,0.1),las=1)
mtext(expression (variance~beta[sor]),side=4,col="black",cex = 1,line=4)

#sne case 1
par(mar=c(2,5,1,3))
plot(subsamples_sne2$j,subsamples_sne2$as.numeric.veg., col="black",
ylim=c(0.1,1),ylab=expression (beta[sne]),xlab="Number of
bins",las=1,cex.lab=1.5)
par(new=TRUE)
plot(1:length(varbeta_sne),varbeta_sne,
axes=F,type="b",pch=19,col="red",xlab=" ",ylab =" ")
axis(4,ylim=c(0,0.05),las=1)
##sne case 2
par(mar=c(2,3,1,6))
plot(subsamples_sne3$j,subsamples_sne3$as.numeric.meanbeta., col="black",
ylim=c(0.1,1),ylab=" ",xlab="Number of bins",las=1)
par(new=TRUE)
plot(1:length(varbeta_sne2),varbeta_sne2,
axes=F,type="b",pch=19,col="red",xlab=" ",ylab =" ")
axis(4,ylim=c(0,0.1),las=1)
mtext(expression (variance~beta[sne]),side=4,col="black",cex = 1,line=4)

######Figure 3.5 LA part#####################
###observed beta values vs coastlength#######
env<-read.delim("Environment.txt", header=T) ##containing details of regional
environmental parameter##
row.names(env)=env[,1]
env[is.na(env)]

env$cumulative= ave(env$Coastline.length..Km.,FUN=cumsum) ###calculating
cumulative coastlength

par(pty="s")
par(mar=c(2,5,1,1))
par(mfrow=c(5,1))
plot(env$cumulative,locs,pch=19,col="black",xlab="Coastline
length",ylab=expression (beta[Obs_ppd]))
cor.test(env$cumulative,locs,method="spearman")

plot(env$cumulative,locs1,pch=19,col="black",xlab="Coastline
length",ylab=expression (beta[Obs_whit]))
cor.test(env$cumulative,locs1,method="spearman")
```

```
plot(env$cumulative,locs_sim,pch=19,col="black",xlab="Coastline
length",ylab=expression (beta[Obs_sim]))
cor.test(env$cumulative,locs_sim,method="spearman")

plot(env$cumulative,locs_sor,pch=19,col="black",xlab="Coastline
length",ylab=expression (beta[Obs_sor]))
cor.test(env$cumulative,locs_sor,method="spearman")

plot(env$cumulative,locs_sne,pch=19,col="black",xlab="Coastline
length",ylab=expression (beta[Obs_sne]))
cor.test(env$cumulative,locs_sne,method="spearman")

###############beta div vs
environment###############################################################################
library(ade4)
library(vegan)
library(MASS)
library(ellipse)
library(FactoMineR)

#####performing RDA with selective variables##############


env=read.csv("Environment - Copy.csv", header=T)##containing details of
regional environmental parameter in correct order##
row.names(env)=env[,1]
env[is.na(env)]=0

env.df=data.frame(env)
prodmean=env.df$Productivity..mgC.m.2.day..
prodrange=env.df$Productivity.range..mgC.m.2.day.
salmean=env.df$Salinity..unit.less.
salrange=as.numeric(env.df$Salinity.range)
tempmean=env.df$Temp.mean..degree.C.
temprange=env.df$Temperature..deg.C..
#coastlength=env.df$Coastline.length..Km.
#river=env.df$Rivers
#shelfwidth=env.df$Shelf.width.m.
#gradient=env.df$Gradient.degree.
cyclonefrq=env.df$Cyclones
oxygen=env.df$Oxygen.ppm.
shelfarea=env.df$Shelf.area.km2.


######Table 3.S1####################
###########################################################
require(Hmisc)
env.df=env.df[,-c(1,8,10,11,12,13,15)]
corr=rcorr(as.matrix(env.new),type="spearman") #####for computing correlation
with significances#
pvalues=corr[["P"]]
rhos=corr[["r"]]
write.table(rhos,file="correlations.csv")
write.table(pvalues,file="pvalues.csv")
```

```
####with shelf area###################
require(Hmisc)
env.df=env.df[,-c(1,9)]
corr=rcorr(as.matrix(env.df),type="spearman") #####for computing correlation
with significances#
pvalues=corr[["P"]]
rhos=corr[["r"]]
write.table(rhos,file="correlations_new.csv")
write.table(pvalues,file="pvalues_spt30.csv")


#########Table
3.4#######################################################################
####multiple glm########
out=summary(glm(ppd1~prodmean+prodrange+salmean+tempmean+temprange+oxygen+cyclonefrq+shelf
#out=summary(glm(ppd~prodmean+prodrange+salmean+salrange+tempmean+temprange+Oxygen+cyclone
summary(out)
finalglm=rbind(out$coefficients)
write.csv(finalglm, file="GLM output live.csv")


############single glm#############################
## GLM overall_Single 566
model2=glm(ppd1~prodmean)
tst2=summary(model2)
tst2
model12=glm(ppd1~prodrange)
tst12=summary(model12)
tst12

model3=glm(ppd1~salmean)
tst3=summary(model3)
tst3

model4=glm(ppd1~tempmean)
tst4=summary(model4)
tst4

model14=glm(ppd1~temprange)
tst14=summary(model14)
tst14

model5=glm(ppd1~oxygen)
tst5=summary(model5)
tst5

model6=glm(ppd1~cyclonefrq)
tst6=summary(model6)
tst6

model16=glm(ppd1~shelfarea)
tst16=summary(model16)
tst16
```

```
#####Figure 3.6 LA ########################################
#####K-s test between null model values and actual WC beta values#####
#####ppd case 1###########
Dist_tidal2=array(0,c(10000,1))
pvalue_tidal2=array(0,c(10000,1))
for(i in 1:10000)
{
  D5=sample(t(subsamples_ppd2$as.numeric.veg.),14,replace = T)
  D6=sample(t(locs),14,replace=T)
  Dist_tidal2[i]=ks.test(as.matrix(D6),as.matrix(D5))$statistic
  pvalue_tidal2[i]=ks.test(as.matrix(D6),as.matrix(D5))$p.value
}
mean(pvalue_tidal2)
mean(Dist_tidal2)
median(pvalue_tidal2)

#####ppd case 2##########
Dist_rest2=array(0,c(10000,1))
pvalue_rest2=array(0,c(10000,1))
for(i in 1:10000)
{
  D7=sample(t(subsamples_ppd3$as.numeric.meanbeta.),14,replace = T)
  D8=sample(t(locs),14,replace=T)
  Dist_rest2[i]=ks.test(as.matrix(D8),as.matrix(D7))$statistic
  pvalue_rest2[i]=ks.test(as.matrix(D8),as.matrix(D7))$p.value
}
mean(pvalue_rest2)
mean(Dist_rest2)
median(pvalue_rest2)
###########################
######whit case 1
Dist_whit1=array(0,c(10000,1))
pvalue_whit1=array(0,c(10000,1))
for(i in 1:10000)
{
  D1=sample(t(subsamples_whit2$as.numeric.veg.),14,replace = T)
  Dist_whit1[i]=ks.test(t(locs1),as.matrix(D1))$statistic
  pvalue_whit1[i]=ks.test(t(locs1),as.matrix(D1))$p.value
}
median(pvalue_whit1)
hist(pvalue_whit1)
hist(Dist_whit1)


#######whit case 2
Dist_whit2=array(0,c(10000,1))
pvalue_whit2=array(0,c(10000,1))
for(i in 1:10000)
{
  D1=sample(t(subsamples_whit3$as.numeric.meanbeta.),14,replace = T)
  Dist_whit2[i]=ks.test(t(locs1),as.matrix(D1))$statistic
  pvalue_whit2[i]=ks.test(t(locs1),as.matrix(D1))$p.value
```

```
}
median(pvalue_whit2)
hist(pvalue_whit2)
hist(Dist_whit2)


###simpson case 1
Dist_simp1=array(0,c(10000,1))
pvalue_simp1=array(0,c(10000,1))
for(i in 1:10000)
{
  D1=sample(t(subsamples_simp2$as.numeric.veg.),14,replace = T)
  Dist_simp1[i]=ks.test(t(locs_sim),as.matrix(D1))$statistic
  pvalue_simp1[i]=ks.test(t(locs_sim),as.matrix(D1))$p.value
}
median(pvalue_simp1)
hist(pvalue_simp1)
hist(Dist_simp1)


#####simpson case 2
Dist_simp2=array(0,c(10000,1))
pvalue_simp2=array(0,c(10000,1))
for(i in 1:10000)
{
  D1=sample(t(subsamples_simp3$as.numeric.meanbeta.),14,replace = T)
  Dist_simp2[i]=ks.test(t(locs_sim),as.matrix(D1))$statistic
  pvalue_simp2[i]=ks.test(t(locs_sim),as.matrix(D1))$p.value
}
median(pvalue_simp2)
hist(pvalue_simp2)
hist(Dist_simp2)


####sorenson case 1
Dist_sor1=array(0,c(10000,1))
pvalue_sor1=array(0,c(10000,1))
for(i in 1:10000)
{
  D1=sample(t(subsamples_sor2$as.numeric.veg.),14,replace = T)
  Dist_sor1[i]=ks.test(t(locs_sor),as.matrix(D1))$statistic
  pvalue_sor1[i]=ks.test(t(locs_sor),as.matrix(D1))$p.value
}
median(pvalue_sor1)
hist(pvalue_sor1)
hist(Dist_sor1)


###sorenson casse 2
Dist_sor2=array(0,c(10000,1))
pvalue_sor2=array(0,c(10000,1))
for(i in 1:10000)
{
  D1=sample(t(subsamples_sor3$as.numeric.meanbeta.),14,replace = T)
```

```
  Dist_sor2[i]=ks.test(t(locs_sor),as.matrix(D1))$statistic
  pvalue_sor2[i]=ks.test(t(locs_sor),as.matrix(D1))$p.value
}
median(pvalue_sor2)
hist(pvalue_sor2)
hist(Dist_sor2)

####sne case 1
Dist_sne1=array(0,c(10000,1))
pvalue_sne1=array(0,c(10000,1))
for(i in 1:10000)
{
  D1=sample(t(subsamples_sne2$as.numeric.veg.),14,replace = T)
  Dist_sne1[i]=ks.test(t(locs_sne),as.matrix(D1))$statistic
  pvalue_sne1[i]=ks.test(t(locs_sne),as.matrix(D1))$p.value
}
median(pvalue_sne1)
median(Dist_sne1)
hist(pvalue_sne1)
hist(Dist_sne1)


##sne case 2
Dist_sne2=array(0,c(10000,1))
pvalue_sne2=array(0,c(10000,1))
for(i in 1:10000)
{
  D1=sample(t(subsamples_sne3$as.numeric.meanbeta.),14,replace = T)
  Dist_sne2[i]=ks.test(t(locs_sne),as.matrix(D1))$statistic
  pvalue_sne2[i]=ks.test(t(locs_sne),as.matrix(D1))$p.value
}
median(pvalue_sne2)
median(Dist_sne2)
hist(pvalue_sne2)
hist(Dist_sne2)


################histograms####################################################
par(mar=c(2,5,1,1))
par(mfrow=c(5,2))
p_ppd=hist(Dist_tidal2)
p_ppd2=hist(Dist_rest2)
p_whit=hist(Dist_whit1)
p_whit2=hist(Dist_whit2)
p_simp=hist(Dist_simp1)
p_simp2=hist(Dist_simp2)
p_sor=hist(Dist_sor1)
p_sor2=hist(Dist_sor2)
p_sne=hist(Dist_sne1)
p_sne2=hist(Dist_sne2)


plot(p_ppd,w=10,col=c("deeppink"),xlim = c(0.2,1),ylim =
c(0,5000),cex.axis=1.5,ann=FALSE)
```

```
plot(p_ppd2,w=10,col=c("deeppink"),xlim = c(0.2,1),ylim =
c(0,5000),cex.axis=1.5,ann=FALSE)


plot(p_whit,col=c("dodgerblue2"),xlim = c(0.2,1),ylim =
c(0,5000),cex.axis=1.5,ann=FALSE)
plot(p_whit2,col=c("dodgerblue2"),xlim = c(0.2,1),ylim =
c(0,5000),cex.axis=1.5,ann=FALSE)



plot(p_simp,col=c("darkorange2"),xlim = c(0.2,1),ylim =
c(0,5000),cex.axis=1.5,ann=FALSE)
plot(p_simp2,col=c("darkorange2"),xlim = c(0.2,1),ylim =
c(0,5000),cex.axis=1.5,ann=FALSE)


plot(p_sor,col=c("green4"),xlim = c(0.2,1),ylim =
c(0,5000),cex.axis=1.5,ann=FALSE)
plot(p_sor2,col=c("green4"),xlim = c(0.2,1),ylim =
c(0,5000),cex.axis=1.5,ann=FALSE)


plot(p_sne,col=c("yellow"),xlim = c(0.2,1),ylim =
c(0,5000),cex.axis=1.5,ann=FALSE)
plot(p_sne2,col=c("yellow"),xlim = c(0.2,1),ylim =
c(0,5000),cex.axis=1.5,ann=FALSE)



##############Figure 3.7 LA
plots#################################################################
ppd1=locs[1:14]



##########correlation of beta diversity with environmental
variables###############

par(mfrow=c(4,2))
par(mar=c(4,4,1,1))

plot(prodmean[1:14],ppd1,xlab="Productivity (mean)", ylab=expression
(beta[ppd]), pch=1,ylim=c(0,1),cex.lab=1)
cor.test(prodmean,ppd1,method = "spearman")

plot(prodrange[1:14],ppd1,xlab="Productivity (range)", ylab=expression
(beta[ppd]), ylim=c(0,1),pch=1,cex.lab=1)
cor.test(prodrange,ppd1,method = "spearman")

plot(salmean[1:14],ppd1, xlab="Salinity (mean)", ylab=expression
(beta[ppd]),ylim=c(0,1),cex.lab=1)
cor.test(salmean,ppd1,method = "spearman")
```

```
plot(salrange[1:14],ppd1, xlab="Salinity (range)", ylab=expression
(beta[ppd]),ylim=c(0,1),cex.lab=1)
cor.test(salrange,ppd1,method = "spearman")

#plot(tempmean[1:14],ppd, xlab="Temperature (mean)", ylab="Mean
PPD",ylim=c(0,5))
#cor.test(tempmean,ppd,method = "spearman")

plot(temprange[1:14],ppd1, xlab="Temperature (range)", ylab=expression
(beta[ppd]),ylim=c(0,1),cex.lab=1)
cor.test(temprange,ppd1,method = "spearman")

#plot(shelfwidth[1:14],ppd, xlab="Coastline length", ylab="Mean
PPD",ylim=c(0,5))
#cor.test(shelfwidth,ppd,method = "spearman")

#plot(gradient[1:14],ppd, xlab="Rivers", ylab="Mean PPD",ylim=c(0,5))
#cor.test(gradient,ppd,method = "spearman")

plot(shelfarea[1:14],ppd1, xlab="Shelf area", ylab=expression
(beta[ppd]),ylim=c(0,1),cex.lab=1)
cor.test(shelfarea,ppd1,method = "spearman")

plot(oxygen[1:14],ppd1, xlab="Oxygen", ylab=expression
(beta[ppd]),ylim=c(0,1),cex.lab=1)
cor.test(oxygen,ppd1,method = "spearman")

plot(cyclonefrq[1:14],ppd1, xlab="Cyclones", ylab=expression
(beta[ppd]),ylim=c(0,1),cex.lab=1)
cor.test(cyclonefrq,ppd1,method = "spearman")

####Figure 3.8 LA plots##########################
#########RDA analysis###################
par(mfrow=c(1,1))
par(mar=c(4,2,2,2))

spe.rda=rda(ta.01 ~ prodrange + salrange +salmean+cyclonefrq+shelfarea,data =
env.df)
anova(spe.rda)
adjR2.tbrda <- RsquareAdj (spe.rda)$adj.r.squared
env.new=env.df[,-c(1,2,7,8,9,10,11)]
require(adespatial)
sel.fs <- forward.sel (Y = ta.01, X = env, adjR2thresh = adjR2.tbrda)
tb_rda.vasc.0 <- rda (ta.01 ~ 1, data = env.df)
sel.osR2 <- ordiR2step (tb_rda.vasc.0 , scope = formula (spe.rda), R2scope =
adjR2.tbrda, direction = 'forward', permutations = 9999)


plot(spe.rda,scaling=2,col="black",display=c("cn","lc"),xlim=c(-5,5),ylim=c(-5,5))
plot(spe.rda,scaling=2,col="black",xlim=c(-80,80),ylim=c(-30,30))

plot(spe.rda,type="n",xlim=c(-80,80),ylim=c(-30,30))
points(spe.rda,display="lc",labels=rownames(ppd),col="black")
text(spe.rda,display = "cn",col="gray41")
```

```
(R2adj=RsquareAdj(spe.rda)$adj.r.squared)
spe.rda.all <- rda(ta.01~ ., data=env)
(R2a.all <- RsquareAdj(spe.rda.all)$adj.r.squared)

ta.prop=ta.prop[-c(15,16),]
require(adespatial)
#env=env[,-c(2,6,10)]
env=env[,-1]
row.names(env)=row.names(ta.01)
forward.sel(ta.01,env,adjR2thresh=R2adj)
###################################################################
## 6. CANONICAL CORRESPONDENCE ANALYSIS using (cca{vegan})

CCAres<- cca(ta.01 ~ prodrange + salrange + salmean + tempmean+cyclonefrq)
plot(CCAres, display=c("sites", "cn"),col="green")
summary(CCAres)
scores(CCAres)
CCAres$CCA$centroids
CCAres$CCA$v
CCAres$CCA$u

#CCAres<- cca(ta.01 ~ prodrange + salrange + salmean + tempmean+shelfarea+
oxygen+cyclonefrq)
#plot(CCAres, display=c("sites", "cn"),col="green")

CCAres<- cca(ta.01 ~ prodrange + prodmean+ salmean + tempmean+shelfarea+
oxygen+cyclonefrq)
plot(CCAres, display=c("sites", "cn"),col="green")
```
