## Supplementary Table (Table S1) for "Controls of spatial grain size and environmental variables on observed beta diversity of molluscan assemblage at a regional scale"

Table S1. Significance (p-values) of Spearman rank correlation test between environmental variables. The significant results are in bold.

|  | Productivity (mean) | Productivity (range) | Salinity (mean) | Salinity (range) | Temperature (mean) | Temperature (range) | Oxygen | Cyclones | Shelf area |
| --- | --- | --- | --- | --- | --- | --- | --- | --- | --- |
| **Productivity (mean)** | 0.692 | 0.059 | **0.001** | **0.002** | **0.003** | 0.923 | 0.817 | 0.061 | 0.081 |
| **Productivity (range)** | NA | **0.002** | 0.056 | **0.045** | 0.533 | 0.056 | 0.852 | 0.538 | **0.005** |
| **Salinity (mean)** |  | NA | **0.011** | **0.000** | 0.274 | 0.375 | 0.970 | 0.874 | **0.000** |
| **Salinity (range)** |  |  | NA | **0.002** | 0.049 | 0.180 | 0.573 | 0.224 | 0.015 |
| **Temperature (mean)** |  |  |  | NA | 0.056 | 0.817 | 0.887 | 0.278 | **0.004** |
| **Temperature (range)** |  |  |  |  | NA | 0.203 | 0.887 | 0.240 | 0.197 |
| **Oxygen** |  |  |  |  |  | NA | 0.185 | 0.809 | 0.533 |
| **Cyclones** |  |  |  |  |  |  | NA | 0.910 | 0.817 |
| **Shelf area** |  |  |  |  |  |  |  | NA | 0.320 |
